# Spinal microcircuits go through multiphasic homeostatic compensations in a mouse model of motoneuron degeneration

**DOI:** 10.1101/2024.04.10.588918

**Authors:** Filipe Nascimento, M. Görkem Özyurt, Kareen Halablab, Gardave Singh Bhumbra, Guillaume Caron, Marcin Bączyk, Daniel Zytnicki, Marin Manuel, Francesco Roselli, Rob Brownstone, Marco Beato

## Abstract

In many neurological conditions, early-stage neural circuit adaption can preserve relatively normal behaviour. In some diseases, spinal motoneurons progressively degenerate yet movement is initially preserved. We therefore investigated whether these neurons and associated microcircuits adapt in a mouse model of progressive motoneuron degeneration. Using a combination of *in vitro* and *in vivo* electrophysiology and super-resolution microscopy, we found that, early in the disease, neurotransmission in a key pre-motor circuit, the recurrent inhibition mediated by Renshaw cells, is reduced by half due to impaired quantal size associated with decreased glycine receptor density. This impairment is specific, and not a widespread feature of spinal inhibitory circuits. Furthermore, it recovers at later stages of disease. Additionally, an increased probability of release from proprioceptive afferents leads to increased monosynaptic excitation of motoneurons. We reveal that in motoneuron degenerative conditions, spinal microcircuits undergo specific multiphasic homeostatic compensations that may contribute to preservation of force output.

## Introduction

At the outset of all but the most rapidly progressing neurodegenerative diseases, the nervous system is remarkably resilient. Specifically, in these early stages of disease progression, neurons and their circuits adapt their properties and wiring to ensure that neural circuit output is maintained within a certain homeostatic physiological range^1^. For example, in Alzheimer’s disease, re-balancing of excitation/inhibition in hippocampal and cortical circuits is responsible for delaying disease progression^2^. In Parkinson’s disease, adaptive changes in the activity of basal ganglia neurons may lead to rather normal behaviour^3^ until at least 30% of substantia nigra compacta dopaminergic neurons have died^4^. That is, as cell death progresses, neural circuits adapt such that behaviour is maintained.

It is evident that such homeostatic mechanisms take place even when the neurons affected are motoneurons^1,5^. In animal models of amyotrophic lateral sclerosis (ALS), there are early changes to motoneuron properties well before substantial motoneuron death^5–11^, but weakness is only detectable when as many as 70% of motoneurons innervating a given muscle have died^12,13^. There are likely many factors underlying this homeostatic regulation involving, for example, neuromuscular junctions^14–16^ and motoneuron properties^5–11^. Yet, for movement to be maintained, it would seem likely that there are as yet unidentified changes to pre-motor spinal cord circuits.

Circuits in the spinal cord integrate information from peripheral and central (distant and local) neurons to ensure that the timing, pattern, and degree of motoneuron activity is such that muscle contraction is behaviourally relevant^17–21^. Since a proportion of spinal interneurons form synapses directly with motoneurons (excitatory, inhibitory, or modulatory^17–21^), and others receive inputs directly from motoneuron axons^22–23,24^, these interneuronal circuits would likely undergo changes as motoneurons die^25,26^. Furthermore, motoneurons are also monosynaptically excited and disynaptically inhibited by activity in proprioceptive afferents – what happens to these “simple” reflex pathways when their targets degenerate?

That is, pre-motor microcircuits function less like an immutable motherboard than a symphony: they are plastic and can adapt to pathophysiological circumstances to maintain adequate motor output^1,27^. In other words, as some drop out, remaining members of the orchestra adapt in order for the symphony of movement to continue.

Given that it is conceivable that these changes to neurons and circuits can themselves lead to disease progression (e.g. through excitotoxicity), it is particularly important to define these changes when considering novel therapeutic strategies aimed at breaking a cycle that, although physiologically adaptive, may in fact be maladaptive^1^.

Here we sought to define how these fundamental spinal microcircuits change in early stages of a mouse model of ALS. We investigated how such changes evolve over the early course of the disease, starting from a point in time prior to the development of overt weakness and extending into the period when there are clear signs of motor weakness. We used the mutant SOD1G93A (mSOD1) mouse model of ALS and performed a combination of *in vitro* and *in vivo* electrophysiology and super-resolution microscopy. We report that in young juvenile mice, there is a dramatic reduction (∼50%) in synaptic strength of recurrent inhibition in the most vulnerable motoneurons. This decrease resulted from impairment in quantal size at motoneuron contacts, associated with a reduced surface postsynaptic glycine receptor (GlyRs) distribution. Remarkably, this finding was confined to the recurrent inhibitory synapses, since group I inhibitory pathways were unchanged. We also detected enhanced Ia monosynaptic input to motoneurons, resulting from an increased probability of transmitter release. In older mice when the pathology has substantially advanced (2-3 months), the initial reduction in the strength of recurrent inhibition was reversed. Our results indicate that pre-motor spinal circuits adapt along the time course of the disease in a non-monotonic manner by changing synaptic drive to motoneurons.

## RESULTS

### Intrinsic properties and firing output are not altered in motoneurons from early juvenile mSOD1 mice

Intrinsic neuronal properties are crucial to the integration of synaptic inputs and to shaping appropriate output, which can impact network function^28^. In neurological conditions, changes in the functional properties of important cell components such as ion channels involved in setting resting conductances and gain can reverse or enhance the effect of alterations in the synaptic wiring of associated microcircuits^29^. In the mSOD1 mouse, motoneuron intrinsic excitability is thought to be affected, with previous reports discussing the possibility of both “hypo” and “hyper” excitability states from embryonic to late-stage adult periods^5–11^. Thus, we first defined subthreshold and firing properties of lumbar motoneurons from our *in vitro* recordings, in order to understand if and how motoneurons themselves are affected in early juvenile mSOD1 mice.

We targeted ventrolateral motoneurons obtained from oblique slices (P15-25; Figure 1A), and classified each as either “slow” or “fast” based on their initial firing profile being immediate or delayed, respectively^22,30,31^ (Figure 1B). Multiple properties (Figures 1C-1F) from these two groups were compared between mSOD1 and Wild Type (WT) motoneurons. Apart from higher rheobase in early firing motoneurons from mutants (43% increase with medium effect size), most subthreshold and repetitive firing properties were similar between genotypes (Figures 1G and S1A-S1G). Analysis of action potential properties, revealed only a 14% smaller fAHP amplitude with medium effect size in early firing motoneurons from mSOD1 mice, and a ∼20% shorter mAHP half-width with a small-medium effect size in late firing mutant motoneurons (Figures 1H and S2A-S2K). Together, the analysis of 18 different cellular properties revealed no large impairment in either immediate or delayed firing motoneurons in early juvenile mSOD1 mice. We then proceeded to probe spinal network function to understand if early alterations in pre-motor circuits could precede motoneuron dysfunction.

**Figure 1.**
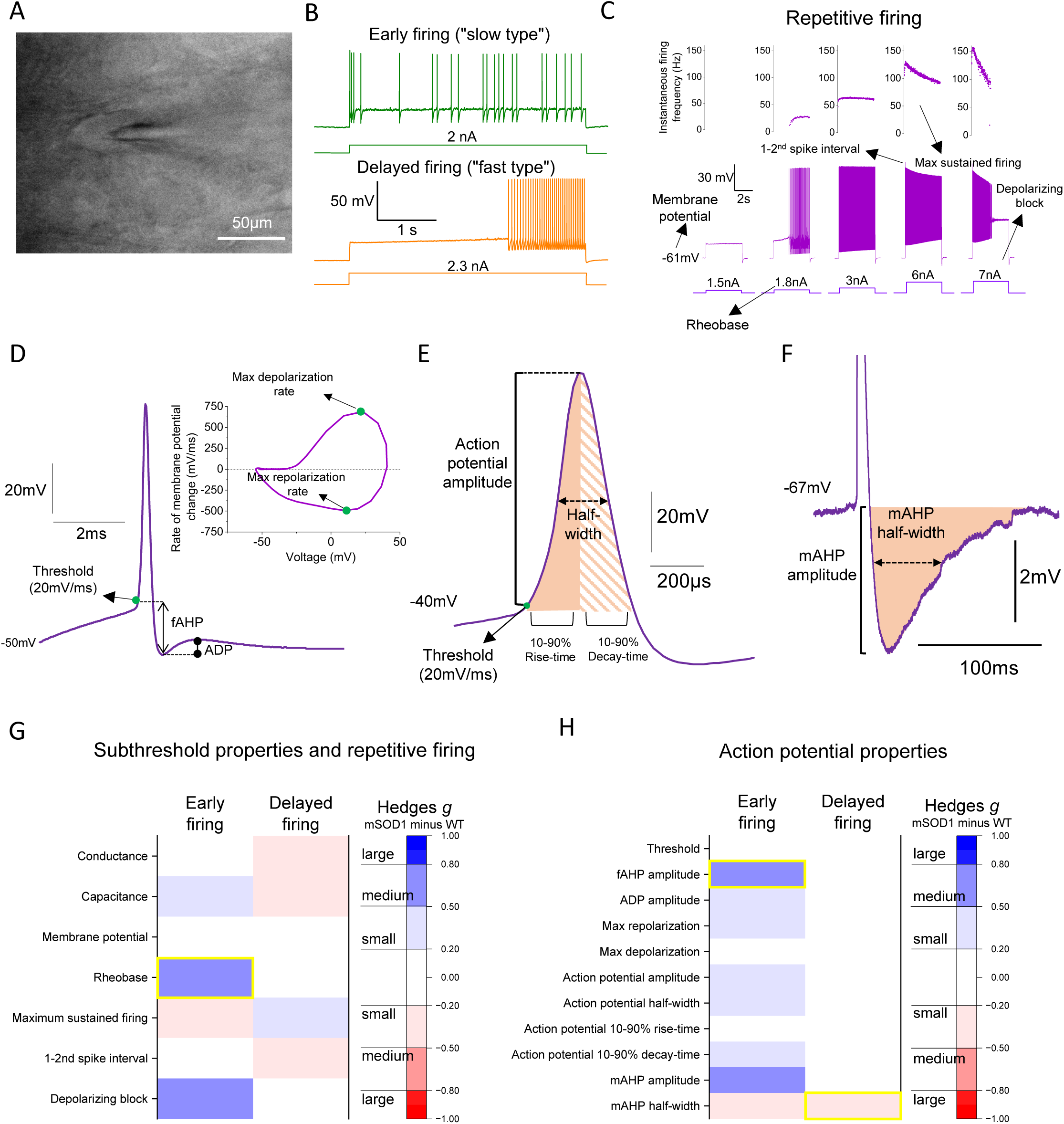
Motoneuron active and passive properties are not substantially altered in early juvenile mSOD1 mice. **(A)** Differential contrast imaging (DIC) image of early juvenile motoneurons (P21) from oblique slices. **(B)** Early and delayed firing profiles used to distinguish between “slow” and “fast” motoneurons. **(C)** Motoneuron response to increasing steps of injected current used to study repetitive firing properties (for simplicity, the scales of the y axis of the last two instantaneous firing plots do not include the value of 1-2^nd^ spike interval). Examples of **(D)** an individual action potential with respective voltage derivative (dV/dt) against voltage plot, **(E)** amplitude, rise and decay time parameters and **(F)** mAHP analyses used to extract information on action potential properties. Heatmaps illustrating absolute mean value of bootstrapped Hedges’ *g* effect size comparisons between early and delayed firing motoneurons from WT and mSOD1 mice for **(G)** subthreshold and repetitive firing and **(H)** spike properties. Yellow boxes highlight comparisons for which bootstrapped 95% confidence interval did not include 0. See also Figures S1-S2 and Tables S1-S2.

### Recurrent excitation is preserved but recurrent inhibition is halved in delayed firing motoneurons in early juvenile mSOD1 mice

For the study of spinal microcircuits, we initially focused on recurrent circuits. These were accessed in oblique slices from P15-25 animals through stimulation of the ventral roots, which excites motor axons that excite other motoneurons^22^ or other groups of interneurons: excitatory V3 interneurons^32^, ventral spinocerebellar tract neurons^33^ or inhibitory Renshaw cells, both projecting back to motoneurons, forming recurrent excitatory or inhibitory loops, respectively^34^.

We first measured recurrent excitation by stimulating the ventral roots and recording an evoked excitatory postsynaptic current (EPSC) near the reversal voltage for Cl^-^ and with pharmacological blockade of inhibition (Figures 2A and 2B). Both absolute and scaled (to cell conductance) recurrent excitation were comparable between genotypes (Figures 2C and S3A). Additionally, recurrent excitation did not correlate with motoneuron conductance and capacitance and the kinetics of the evoked current were similar between WT and mSOD1 mice (Figures S3C-S3E, S3I and S3J).

**Figure 2.**
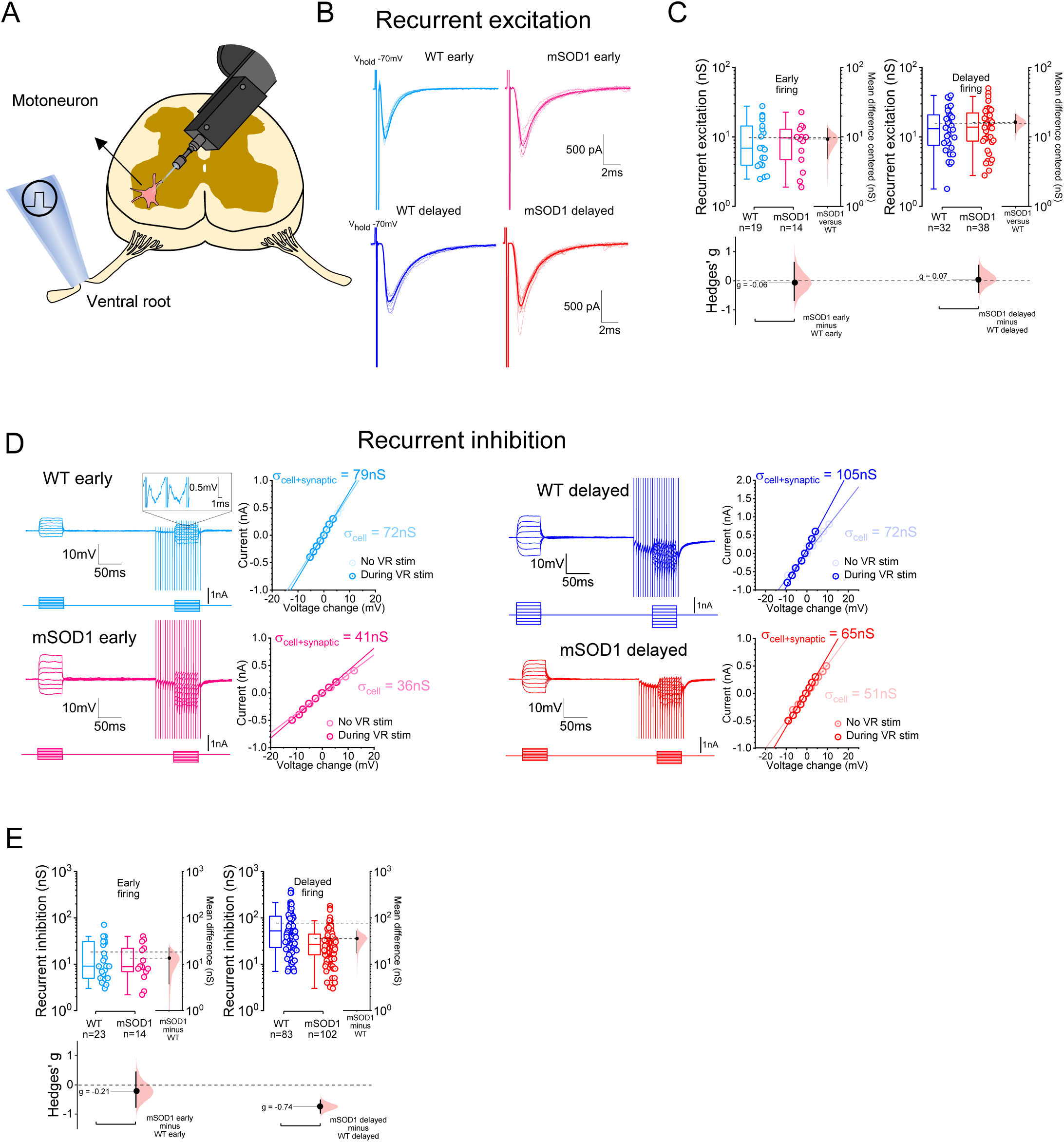
Recurrent inhibition is halved in early juvenile mSOD1 animals. **(A)** Schematic of the oblique spinal cord slice preparation used to obtain whole-cell patch clamp recordings from motoneurons. **(B)** Examples of ventral root-evoked recurrent excitatory EPSCs recorded from motoneurons (at 3-5x the threshold required for an initial synaptic response). Estimation plots for **(C)** absolute recurrent excitation. **(D)** Examples of current-voltage responses obtained before and during high-frequency ventral root stimulation (200 Hz) used to measure recurrent inhibition. Zoomed-in box (WT early trace) illustrates example IPSPs evoked during the train. Plots for **(E)** absolute synaptic conductances for recurrent inhibition. Estimation plots with all individual values and respective box-plots shown along with respective bootstrapped mean difference and bootstrapped Hedges’ *g*. See also Table S3.

Next, we examined recurrent inhibition by comparing motoneuron conductance at rest with the conductance during a high-frequency (200 Hz) ventral root stimulation period during which recurrent inhibition reaches a steady-state^35^ (Figure 2D). Absolute and scaled inhibitory conductances were reduced by ∼50% in delayed firing motoneurons from mSOD1 mice with a medium-large effect sizes (Figures 2E and S3B). For early firing motoneurons, absolute conductances remained unchanged, but scaled conductances were ∼36% lower in mSOD1 with a medium effect size, although with a difference with a large confidence interval. The size of recurrent inhibition correlated with cell capacitance in control mice but not in mutants (Figures S3F-S3H), hinting that perhaps inhibitory conductances in mSOD1 mice are especially reduced in large motoneurons that are known to be more vulnerable to disease progression in ALS and mSOD1 mice^36^. These results indicate that while recurrent excitation is not substantially affected in early juvenile mSOD1 mice, recurrent inhibition mediated by Renshaw cells is preferentially reduced in delayed firing motoneurons from mutants. Given that this is a disynaptic circuit, we then proceeded to pinpoint the locus of synaptic impairment: motoneuron and/or Renshaw cell.

### Motoneuron input to Renshaw cells is preserved in early juvenile mSOD1 mice

To test if synaptic function was compromised at motoneuron-Renshaw cell contacts, we performed whole-cell patch clamp recordings from identified Renshaw cells. For this, we crossed mSOD1 animals with mice that express enhanced green fluorescent protein (EGFP) under the control of the glycine transporter 2 (GlyT2) promoter^37^, thus allowing the targeting of Renshaw cells, identified as cells located in the most ventral area of lamina VIII of oblique slices that express the fluorescent reporter and that receive excitation following ventral root stimulation^34,38^ (Figures 3A-3C).

**Figure 3.**
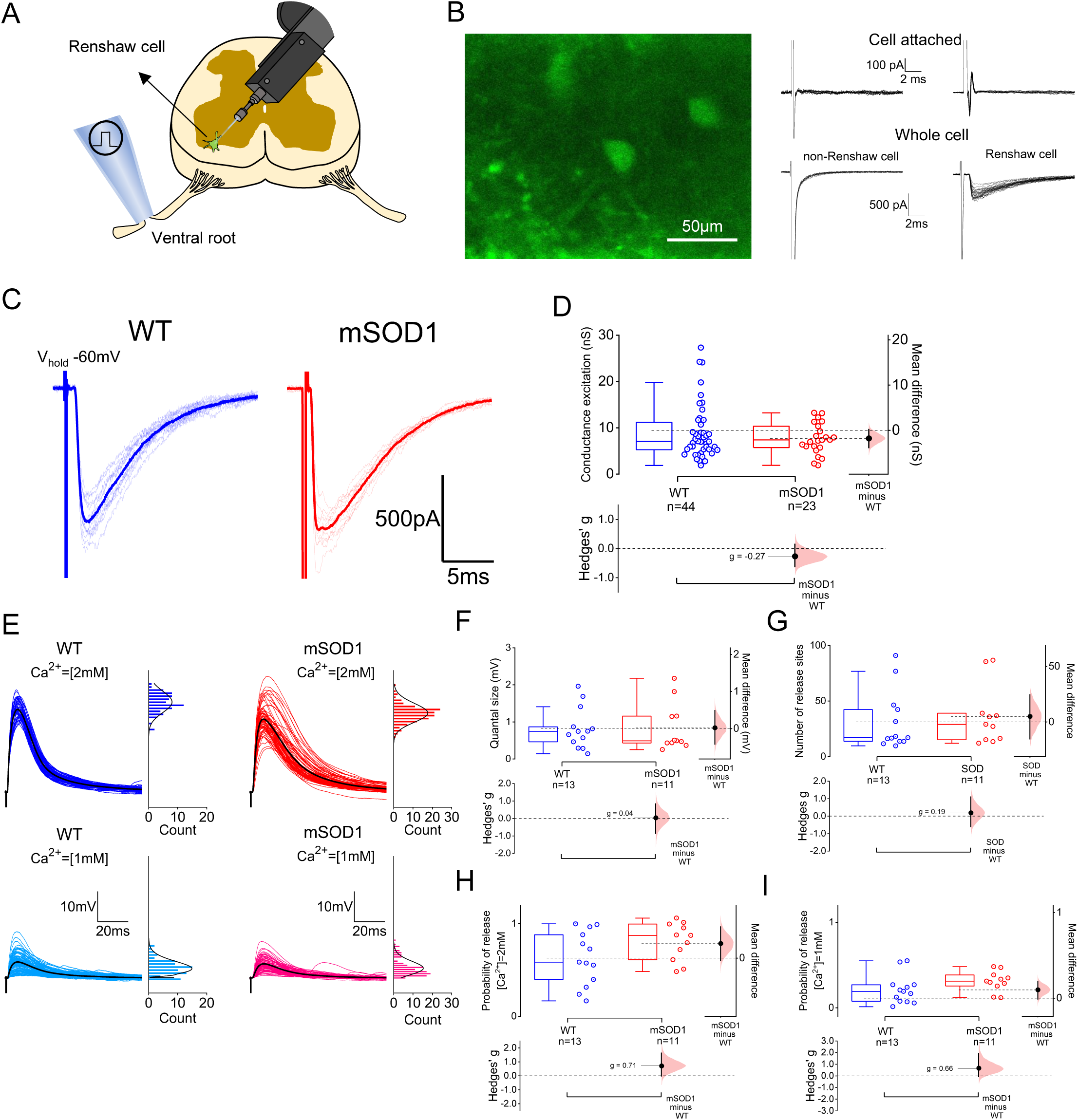
Motoneuron input to Renshaw cells is preserved in early juvenile mSOD1 mice. **(A)** Schematic of the oblique spinal cord slice preparation now used to target Renshaw cells, identified as **(B)** GlyT2 EGFP+ cells located in the most ventral area of lamina VIII that **(C)** receive ventral root-evoked excitation. Group data obtained for **(D)** absolute synaptic conductances for both WT and mSOD1 mice. **(E)** Representative traces showing EPSPs recorded from Renshaw cells in the presence of 2mM (top) and 1mM (bottom) of Ca^2+^, next to respective histogram count, that were used to perform BQA (sweeps were baselined for representation purposes and black IPSP represents averaged trace). Group plots showing data obtained from BQA on parameters such as **(F)** quantal size, **(G)** number of release sites, and probabilities of release with **(H)** 2mM and **(I)** 1mM of extracellular Ca^2+^. Estimation plots with all individual values and respective box-plots shown along with respective bootstrapped mean difference and bootstrapped Hedges’ *g*. See also Table S4.

We evaluated the synaptic drive Renshaw cells receive from motoneurons by estimating the absolute and scaled conductances from the ventral root-evoked EPSCs, and found no differences between groups (Figures 3D and S4C). Intrinsic properties from Renshaw cells were similar, but the rise times of evoked EPSCs were ∼30% slower with a large effect size in mSOD1 mice (Figures S4A-S4B and S4D-S4H). Additionally, we developed a discrete-grid exact inference implementation of Bayesian Quantal Analysis (BQA) (see analytical derivations in Supplemental information) to further characterize and compare quantal parameters between groups. Ventral-root evoked excitatory postsynaptic potentials (EPSPs) obtained from Renshaw cells in the presence of 2 and 1mM of extracellular Ca2+ were used for BQA (Figure 3E). We found that quantal size, number of release sites, and probability of release obtained from BQA, were comparable between WT and mSOD1 mice (Figures 3F-3I), thus further confirming that changes at motoneuron-Renshaw cell synapses are not the cause for impaired recurrent inhibition observed in motoneurons.

### The reduction of Renshaw cell inhibition of large motoneurons is due to decreases in quantal size and glycine receptor clustering

To examine if the locus of impairment was at Renshaw cell-motoneuron contacts, we ran BQA by recording root-evoked inhibitory postsynaptic potentials (IPSPs) obtained from disease susceptible delayed firing motoneurons, slightly above the reversal potential for inhibition (see methods), in the presence of 2 and 4mM of extracellular Ca^2+^ (Figure 4A). We found that while the number of release sites and probabilities of release were unchanged, the quantal size in mSOD1 was reduced by ∼30% with a large effect size (Figures 4B-4E). To further support these findings, we recorded asynchronous IPSCs (aIPSCs) from large motoneurons (putative delayed-firing) following 200 Hz ventral root stimulation in the presence of high concentrations of Sr^2+^, which is known to desynchronize and prolong pre-synaptic vesicular release^39^ (Figure 4F). Prior to replacing extracellular Ca^2+^ with Sr^2+^, in a subset of experiments, we recorded some ventral root-evoked IPSCs, and those from mSOD1 mice had a slower rise and decay phases (Figures S5A-S5C). When Sr^2+^ was added, we observed that the amplitude of aIPSCs was smaller in mSOD1 mice, with a ∼30% reduction for absolute aIPSC conductance with a small-medium effect size (Figure 4G), and ∼50% decrease for scaled aIPSC conductances (Figure S5D). The results from BQA and aIPSCs are both consistent with a decrease in quantal size at Renshaw cell-motoneuron contacts that could be responsible for an early reduction in recurrent inhibition in mSOD1 mice.

**Figure 4.**
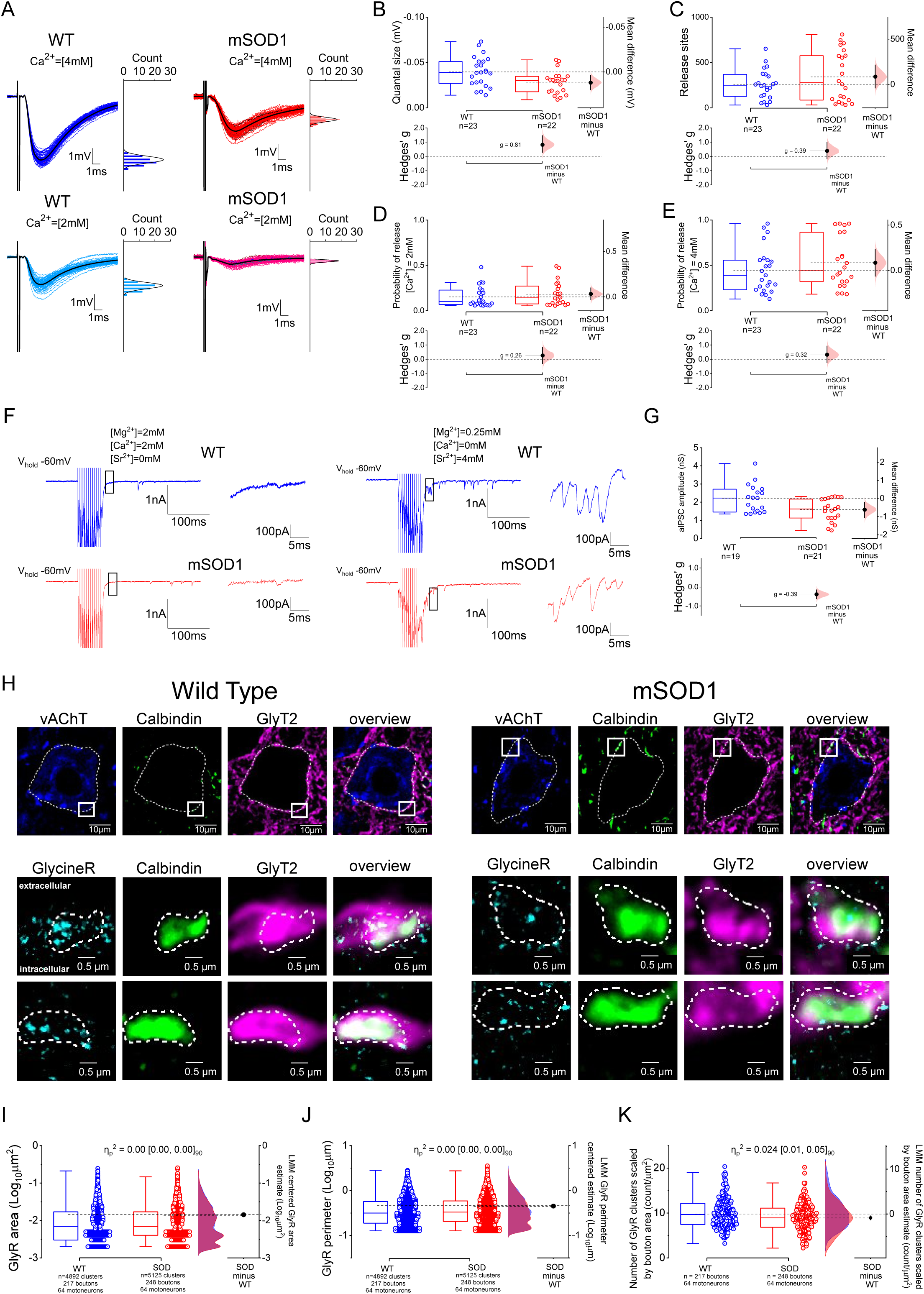
Impairment in recurrent inhibition in early juvenile mSOD1 mice is due to a reduction in quantal size at Renshaw cell to motoneurons contacts which is associated with decreased number of postsynaptic glycine receptors per bouton. **(A)** Examples of IPSPs (baseline adjusted for representation) to ventral root-stimulation obtained in the presence of 4mM and 2mM of extracellular Ca^2+^, next to respective histogram counts. BQA parameters for **(B)** quantal size, **(C)** number of release sites and probability of release for **(D)** 2mM and **(E)** 4mM of extracellular Ca^2+^. **(F)** Examples of voltage-clamp motoneuron responses to 200 Hz ventral root stimulation without (left) and with 4mM of Sr^2+^ (right), a large ion that extends synaptic release thus allowing to detect asynchronous IPSCs (aIPSCs) following extracellular stimulation (see zoomed in window). **(G)** Estimation plots for of aIPSC amplitude conductance. **(H)** Examples of P21 mice identified Renshaw cell boutons (GlyT2^+^ and Calbindin^+^) juxtaposed to motoneurons (vAChT), with labelled clusters of GlyR (GlycineR) for both control (left) and mSOD1 (right) mice. Top row shows motoneuron somata. The boxes in the top row indicate the position of the two boutons highlighted in the bottom raw (represented rotated). Group data for GlyR **(I)** area, **(J)** perimeter and **(K)** number per bouton. Estimation plots with all individual values and respective box-plots shown with respective bootstrapped mean difference and bootstrapped Hedges’ *g* for **(B-E)** plots and Kernel smooth distribution with respective Linear Mixed Model (LMM) estimates shown for **(I-K)** plots. Hierarchical bootstrap used for **(G)** with mean amplitude per motoneuron used in box-plots. See also Tables S5-S6 and Figure S7.

To determine whether the reduction in IPSCs is specific to Renshaw cells, or widespread (i.e. affecting all motoneuron glycinergic synapses), we recorded glycinergic miniature IPSCs (mIPSCs) in large motoneurons (Figure S6A). Although the absolute mIPSC amplitude was reduced by ∼28% with a small-medium effect size, scaled mIPSCs were not different between groups as well as their rise and decay-times. However mIPSC inter-event interval was increased by ∼20% in mSOD1 mice with a small-medium effect size (Figures S6B-S6F). The change in mIPSCs frequency does not explain alterations in quantal size, thus indicating that reduction in quantal size in recurrent inhibition is likely not a generalized feature of pre-motor glycinergic synapses.

Given that changes in quantal size are usually associated with postsynaptic alterations, we next used super-resolution microscopy to quantify GlyRs opposing glycinergic terminals originating from Renshaw cells (identified by their immunoreactivity for GlyT2 and for Calbindin, Calb^+^) (Figure 4H). We observed no differences in GlyR cluster area or perimeter, but the number of GlyR clusters per bouton was ∼10% smaller in mSOD1 mice with a small-medium effect size (Figures 4I-4K). These results suggest that reduced GlyR distribution is contributing to the reduction in quantal size responsible for reduced recurrent inhibition in large motoneurons in mSOD1 mice.

### Monosynaptic Ia excitation is increased in early juvenile mSOD1 mice due to higher probability of release from Ia terminals but disynaptic group I afferent inhibition is not affected

Having characterized recurrent motor circuits, we then diverted our focus towards sensory-related spinal pathways. For this, we used a novel ventral horn-partially ablated spinal cord preparation from early juvenile mice (P14-21) containing both L4 and L5 segments and dorsal roots intact, permitting the measurement of group I afferent-related responses such as monosynaptic Ia excitation (Figures 5A-5B) and disynaptic Ia/Ib inhibition (Figures 5F-5G) through whole cell-patch clamp recordings^35^, targeting large motoneurons (see methods). We first compared intrinsic neuronal properties (resting conductance and capacitance), since in this preparation we preferentially record from the dorsolateral motor nuclei whereas in slices the recordings were from ventrolateral motoneurons^35^; and we found no differences between WT and mSOD1 mice (Figures S8A and S8B).

**Figure 5.**
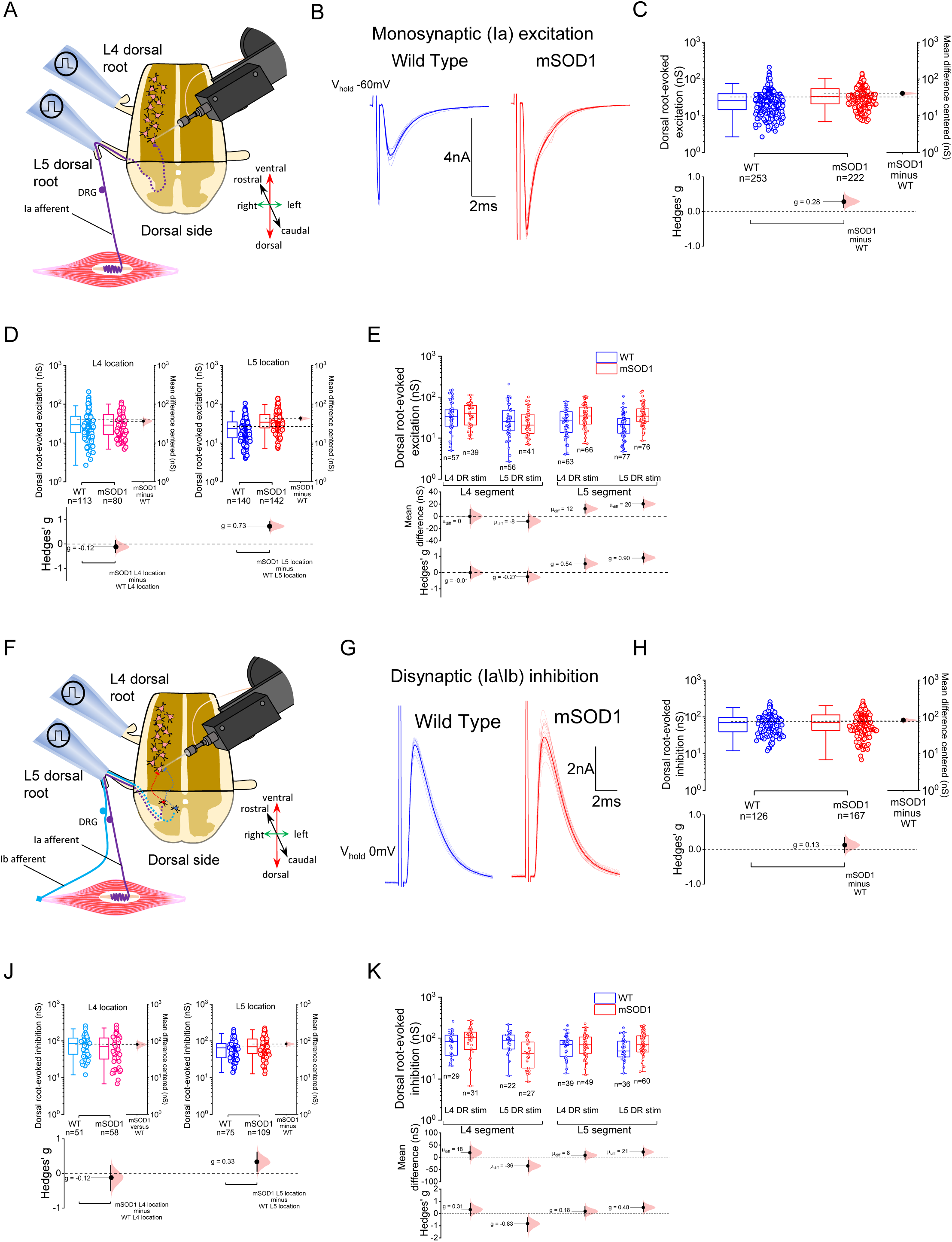
Monosynaptic Ia excitation is increased in early juvenile mSOD1 mice but disynaptic Ia/Ib inhibition remains unchanged. **(A)** Schematic of the ventral horn-partially ablated *in vitro* longitudinal spinal cord preparation with L4 and L5 segments and roots intact, used to study monosynaptic Ia excitation. **(B)** Example of monosynaptic EPSCs obtained following dorsal root stimulation (at 1.5-3x the threshold required to evoke an initial synaptic response). Group data for absolute dorsal root-evoked excitation for **(C)** all responses obtained, responses split by **(D)** location and **(E)** according to stimulated root and location. **(F)** Representation of group I afferent inhibitory pathways (Ia/Ib) studied in *vitro*, with **(G)** examples of disynaptic IPSCs obtained following dorsal root stimulation. Data obtained on absolute synaptic conductance for **(H)** all responses, responses grouped by **(I)** location and **(H)** organized by stimulated root and segment. Estimation plots with all individual values and respective box-plots shown along with respective bootstrapped mean difference and bootstrapped Hedges’ *g*. See also Table S11-S17. DRG – dorsal root ganglion

We next recorded dorsal root-evoked EPSCs obtained close to the voltage equilibrium for inhibition (see methods), and detected a small effect size increase in absolute (∼20%) but not in scaled conductance in mSOD1 mice, indicating that despite the larger absolute conductance observed in SOD1 mice, the effect of Ia excitation on motoneurons would be similar across genotypes (Figures 5C and S9A). Additionally, the magnitude of the Ia EPSC was not proportional to cell size, since the responses correlated weakly with resting motoneuron conductance and capacitance (Figures S8G-S8I). An advantage of this longitudinal preparation is the identification of the position of the postsynaptic cell so that the recordings could be partitioned on the basis of both anatomical location (L4 or L5 segment) and stimulated dorsal root (L4 or L5).This allowed to determine whether observations are generalizable to the lumbar cord or whether there are differences between different segments and roots given the caudal-rostral nature of the progression of the disease in mSOD1 mice^40^. Our analysis showed that mSOD1 motoneurons from L5 tend to receive increased absolute (∼65% more, medium-large effect size) an scaled (∼50% more, small-medium effect size) Ia excitation (Figures 5D-5E and S9B-S9D). To investigate short-term plasticity of Ia synapses, a second EPSC was evoked 30ms after the first, which revealed a 4% reduction in the paired-pulse ratio in mSOD1 mice with small-medium effect size (Figures S10A-S10D). The dorsal root-evoked EPSC had a slightly faster (4%) rise time in mSOD1 mice with a small effect size, more evident in motoneurons in L5 segments (7% faster, with a small-medium effect size), with the decay time remaining unchanged (Figures S11A-S11G). These data indicate that the synaptic strength of monosynaptic Ia excitation is increased in early juvenile mSOD1 mice, with an effect preferentially detected on motoneurons in caudal lumbar segments, and that short-term plasticity of Ia afferent synapses is also altered.

To study disynaptic Ia/Ib inhibition, we recorded dorsal root-evoked IPSCs at holding voltages near the estimated reversal for excitation (Figure 5F), and found no differences between genotypes in both absolute and scaled inhibitory conductances when plotting all root responses (Figures 5H and S9E). Splitting data by location and stimulating root, revealed that, although absolute inhibitory conductances were ∼20% higher in motoneurons from mSOD1 located in L5 regions with a small-medium effect size, the net effect shown by scaled synaptic conductances was similar between groups (Figures 5J-5K and S9F-S9H). We also found that responses to L5 dorsal root stimulation from motoneurons located in L4 segments had smaller absolute (40% decrease with large effect size) and scaled (30% reduction with medium effect size) synaptic conductances in mSOD1 mice, perhaps indicative of a subtle impairment in the strength of ascending inter-segmental Ia/Ib inhibitory inputs or a product of the anatomical spread of that small number of responses. Short-term plasticity of Ia/Ib inhibition, given by the paired-pulse ratio of evoked IPSCs, was marginally altered in mSOD1 mice (3% change with small-medium effect size), due to an increased ratio found predominantly in paired root responses obtained from L5 segments (4% increase with small-medium effect size; Figures S10E-S10H). Additionally, the evoked IPSC rise and decay times tended to be faster in mSOD1 mice by 8% and 15%, respectively, with small-medium effect sizes, but only in responses obtained from cells located within the L4 segment area (Figures S11H-S11M). These data on sensory-evoked disynaptic Ia/Ib inhibition indicate that, unlike the observations obtained on recurrent inhibition, there is no substantial impairment in the synaptic strength to lumbar motoneurons in early juvenile mSOD1 mice.

From the dorsal-root evoked EPSCs, we also obtained information on post-activation depression which is an activity-dependent long-term depression of Ia afferents that can last several seconds^41–43^. Decrease in long-term depression has been reported in people with ALS^25^. From our conditioning protocol (see methods), no differences in post-activation depression existed between genotypes when comparing all root responses. However, we detected a ∼10% smaller ratio with small-medium effect size, in L5 root responses obtained from mSOD1 motoneurons from L5 segments (Figures S12A-S12D).

The above results indicate the presence of early synaptic alterations in Ia afferents in mSOD1 animals. To ascertain the nature of the synaptic dysfunction, we performed BQA on monosynaptic dorsal root-evoked EPSCs (Figure 6A). We systematically targeted large motoneurons located in the more caudal regions (L5 segment) in which differences in Ia excitation synaptic conductance were more pronounced. When comparing experimental groups, BQA revealed that quantal size and number of release sites were similar between mSOD1 and WT animals mice (Figures 6B and 6C). However, the probabilities of release in both 2 and 4mM of Ca^2+^ were respectively 50% and ∼30% higher in mSOD1 mice, differences with a large and quasi-large effect sizes (Figures 6D and 6E). This finding could explain why monosynaptic afferent excitation is increased in mSOD1 mice. Furthermore, large probabilities of release are also known to decrease the paired-pulse ratio^44–46^, which could also account for the decrease in short-term plasticity of Ia excitation.

**Figure 6.**
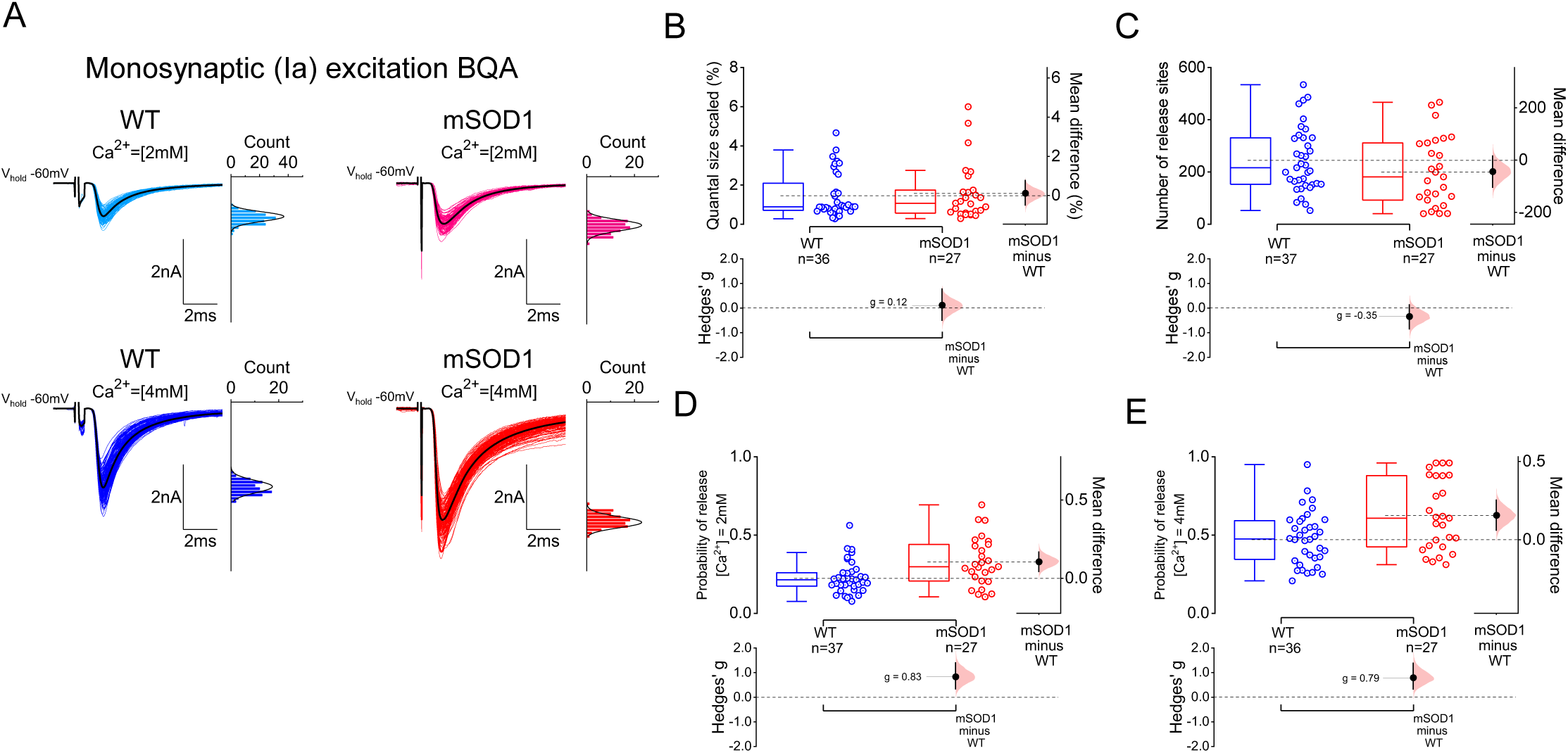
Increase in monosynaptic Ia excitation in early juvenile mSOD1 mice is associated to higher probability of release from Ia afferents. **(A)** Examples of EPSCs obtained in the presence of 2mM (top) and 4mM (bottom) of extracellular Ca^2+^, with respective histogram counts next to traces (black sweep represents averaged trace). BQA estimates for **(B)** quantal size scaled, **(C)** number of release sites and probabilities of release **(D-E)**. Estimation plots with all individual values and respective box-plots shown along with respective bootstrapped mean difference and bootstrapped Hedges’ *g*. See also Table S18.

In summary, data obtained on group I afferents microcircuits identified an increase in the synaptic drive from Ia afferents to lumbar motoneurons (specially to those located more caudally) due to a higher probability of release from the afferents, whereas Ia/Ib disynaptic inhibition is not substantially affected in early juvenile mSOD1 mice.

### Early reduction in recurrent inhibition is compensated in later, advanced stages of disease progression in mSOD1 mice

The previous data was collected from mice that have reached mature stages of motor development (P14-25) but at the very early stages of disease. Furthermore, the time-course of abnormalities in mSOD1 mice might not be unidirectional, with lumbar motoneuron properties undergoing oscillatory alterations throughout disease progression^5^. Whether these oscillations are unique to motoneurons themselves or they also involve pre-motor spinal circuitry is unknown, but it has been hypothesized that spinal microcircuits might undergo multiphasic homeostatic compensations to balance motoneuron output in ALS^1^.

In fact, our observed increase in Ia excitation, is completely reversed a few weeks later just before denervation started in mSOD1 mice^47^. Also, while synaptic stripping of glycinergic inputs has been detected in early-juvenile mSOD1 mice^48^, Renshaw cells start to display a specific type of compensatory sprouting to motoneurons in the second and third month of adult life, which fades considerably at later stages^49^. Furthermore, our GlyR imaging revealed that decreased cluster density still persists in older P45 mutant animals (Figure S7K). We therefore asked whether there is reversal of the initial reduction in quantal size in recurrent inhibitory circuits at later stages of disease. Since *in vitro* preparations are not suitable for studying recurrent circuits in older animals (>P30) as the synaptic drive is heavily reduced in older tissue due to poor motoneuron viability^35^, we performed two alternative independent measurements of recurrent inhibition at later time-points of disease in mSOD1 mice: 1) *in vivo* sharp electrode recordings from motoneurons and 2) EMG recordings.

*In vivo* intracellular recordings were obtained from P48-56 mixed-background mSOD1 mice (see methods). Motoneurons were identified by the presence of an antidromic response to either L4 or L5 ventral root stimulation, with the root that did not elicit an antidromic spike used to evoke recurrent inhibition (Figure 7A). No differences in evoked synaptic conductance were found between WT and mSOD1 animals (Figures 7B and S13D), which contrasted with the patch data from early juvenile mice. Since *in vivo* recordings targeted distal muscle-innervating motoneurons (dorsolateral nuclei), and slice recordings focused on proximal-innervating motoneurons (ventrolateral nuclei), we also estimated recurrent inhibition in the dorsolateral motor nuclei from 2-3 weeks old mice (Figures S13A-S13C). Although we observed a reduction the absolute evoked conductance in early firing motoneurons from mSOD1 mice (∼60% with large effect size), the net effect, given by the scaled conductance was not different. However, for delayed firing motoneurons we observed impairments in both absolute (29% reduction with small-medium effect size) and scaled (40% decrease with medium effect size) conductances in mutant early juveniles.

**Figure 7.**
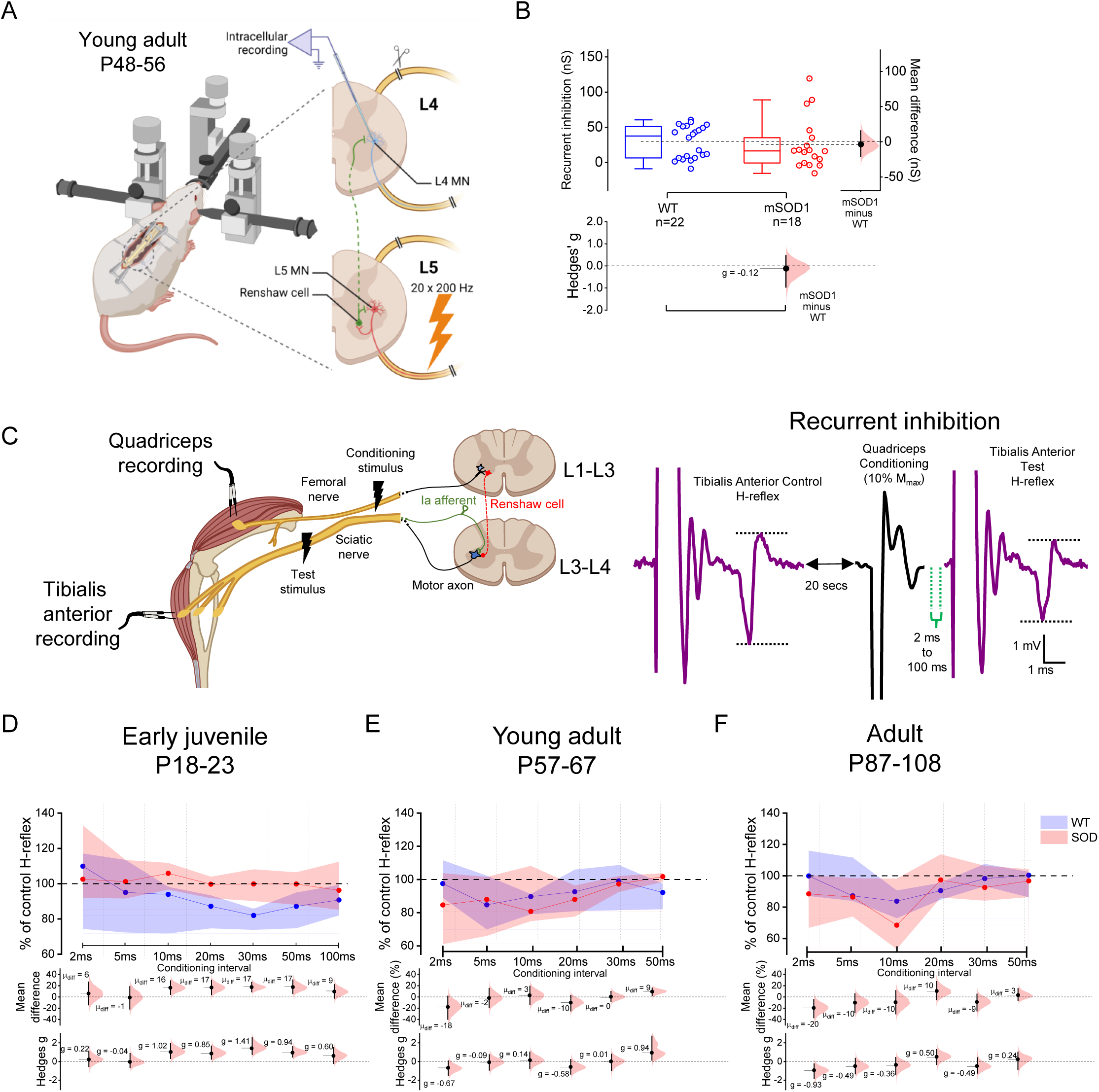
The initial reduction in recurrent inhibition in early juvenile mSOD1 mice is compensated at later adult stages. **(A)** Summary of the recording setup used to perform *in vivo* sharp-electrode recordings from motoneurons from P48-56 mice, with cells identified through antidromic stimulation of the L4 or L5 ventral roots and recurrent inhibition estimated by stimulating the adjacent root. Estimation plots for **(B)** absolute conductances for recurrent inhibition from *in vivo* motoneuron recordings. Schematic and example traces illustrating the EMG recordings used to obtain motor and H-reflex responses from quadriceps and tibialis anterior muscles, and the conditioning protocols used to estimate **(C)** recurrent inhibition. Data obtained for the different age-ranges tested for **(D-F)** recurrent inhibition. Estimation plots with box-plots with individual values shown for *in vivo* sharp-electrode recurrent inhibition, and box-plots shown as median (dot) and interquartile range (shaded area) for EMG-estimated recurrent inhibition, along with respective bootstrapped mean difference and bootstrapped Hedges’ *g*. See also Tables S20 and S23.

Although *in vitro* and *in vivo* motoneuron recordings were obtained from congenic and mixed background mSOD1 mice, respectively, we would expect the results to be comparable given the resemblance in disease progression between the lines^50^. However, to further improve comparability between time-points we performed an additional independent measurement to test for differences in recurrent inhibition in congenic mSOD1 -EMG recordings.

We performed EMG recordings from the Tibialis Anterior (TA) and Quadriceps (Q) of anaesthetised mice from three different age groups: 1) early juvenile (P18-23), same age as the *in vitro* recordings; 2) young adult (P57-67) an age range similar to that of the *in vivo* motoneuron recordings and a period of disease progression during which significant denervation has occurred^51^; 3) adult animals (P87-108), a stage at which mSOD1 mice start displaying clear signs of motor impairment^52^.

We first evaluated neuromuscular impairment, and found no difference in maximal H-reflex (H_max_), but decreasing maximum motor (M_max_) and H_max_/M_max_ responses in both TA and Q muscles as age progressed, with a ∼19% reduction in TA M_max_ (medium effect size) but not in Q of early juvenile (P18-23) mutant mice, indicative of the distal to proximal progressive pathogenesis in mSOD1 mice^53^ (Figures S14A-S14D).

We next performed EMG recordings to also study post-activation depression of Ia afferents, known to be affected in ALS^25^. This depression was investigated using paired-pulse H-reflex conditioning (see methods; Figure S15A), and we detected impairments in mSOD1 mice at different pulse intervals in all ages test: in early juvenile (P18-23) animals, we detected a 18% reduction at 5 sec conditioning interval with a large effect size; in young adults (P56-67) we found a ∼20% reduction at 500 ms with a large effect size; for 3-months-old adults (P87-108) 200 ms and 1 sec conditionings were reduced by ∼20% with large effect sizes (Figures S15B-S15D). Although the intervals affected vary between age-groups, which could be attributed to the inter-individual variability of post-activation depression^54^, our EMG data indicate that the Ia depression is altered in mSOD1 mice.

Finally, we used EMG recordings to infer about time-dependent alterations in recurrent inhibition and improve comparability between our in vitro and in vivo motoneuron datasets. To do this, we measured heteronymous recurrent inhibition by conditioning TA H-reflexes with femoral nerve stimulation using a range of different inter-stimulus intervals (see methods; Figure 7C). In early juvenile (P18-23) mSOD1 mice, we observed reductions of ∼20% with large effect sizes, in heteronymous inhibition obtained at 10, 20, 30 and 50 ms conditioning intervals (Figure 7D). In contrast, in young adult (P57-67) mice, we observed no differences between groups, and for adult animals (P87-108) there was only an increase in H-reflex conditioning (∼20%, large effect size) in mutant mice at very short (2ms) conditioning intervals (Figures 7E-F). These data indicate that, similar to the *in vitro* data, recurrent inhibition is impaired in early juvenile mSOD1 mice, and similar to the *in vivo* recordings, recurrent inhibition then recovers in young adult animals (P57-67). The data from a conditioning interval (2ms) hints that some component of recurrent inhibition might actually be exacerbated in mSOD1 animals that display clear signs of weakness (P87-108), a feature which has been observed in early-stage patients^55^.

## Discussion

In this work, we show that, in an animal model with progressive motoneuron degeneration, pre-motor spinal microcircuits such as recurrent inhibition and monosynaptic Ia excitation are altered months before cell death and onset of motor abnormalities. These alterations are not unidirectional, with an initial reduction in the synaptic strength of recurrent inhibition being compensated at later stages of disease progression, and an increment in Ia monosynaptic excitation being reversed just a few weeks later^47^ (Figure 8).

**Figure 8.**
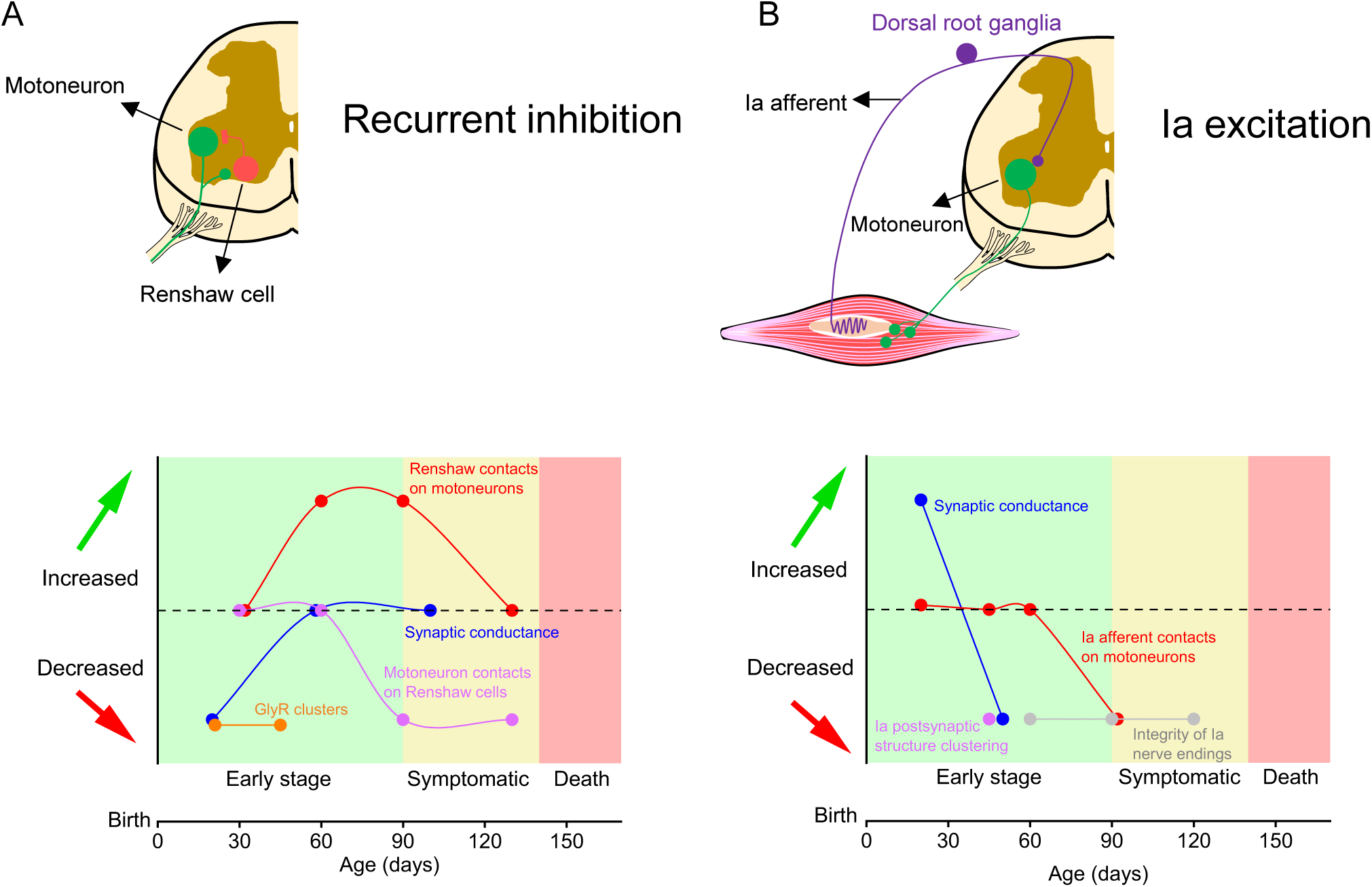
Homeostatic responses in spinal microcircuits are multiphasic throughout the course of disease progression in mSOD1 mice. Summary of identified synaptic alterations in **(A)** recurrent inhibition and **(B)** Ia monosynaptic excitation in mSOD1 mice obtained from this work and previous studies^47,49,69^.

### Motoneuron electrophysiological properties are relatively preserved in the initial stages of disease

Motoneuron intrinsic excitability in mSOD1 mice has been a heated topic of discussion, with studies putting forward the idea that motoneurons can be either “hyperexcitable”^7,30^ or “hypoexcitable”^6,8^. In the current work, we detected differential effects on “slow” vs “fast” type motoneurons: there was an increase in rheobase and fAHP in “slow type,” and a longer mAHP in “fast type” motoneurons. That is, there are alterations in some type-matched motoneuron electrical properties in 2-3 week old mSOD1 mice (Figure 1).

In contrast to our findings, previous *in vivo* recordings from P30-60 mSOD1 mice have identified increased motoneuron conductance and hyperpolarized resting potential^5^, and studies that used *in vitro* spinal cord preparations from embryonic or early postnatal (<P12) mSOD1 mice have reported decreases in input conductance and rheobase^9–11,30,56^. Some of the discrepancies with our findings could be due to 1) differences in the time-points of data acquisition (the changes to motoneurons have been hypothesised to be multiphasic^5^), 2) mSOD1 strains used (e.g. congenic *vs* mixed-background), and 3) contrasts between *in vitro* vs *in vivo* electrophysiology methodologies. But we also note that our dataset includes a large number of recorded cells per genotype (∼100 motoneurons per genotype) and distinguishes between putative slow and fast motoneurons, which are differentially vulnerable to ALS progression^55^. We conclude that at the early-stage in which we obtained our *in vitro* recordings (2-3 weeks old mature mice), alterations to electrophysiological properties of either fast or slow motoneurons are not a prominent feature in mSOD1 mice.

### Early-stage neuronal plasticity involves alterations in inhibitory and excitatory spinal microcircuits

We found the strength of recurrent inhibition of motoneurons reduced by ∼50% in delayed firing (vulnerable) motoneurons from early juvenile mSOD1 mice. This was not due to alterations in synaptic connectivity between motoneurons and Renshaw cells, but to a decrease in quantal size at Renshaw cell to motoneuron synapses. This impairment was associated with a ∼10% reduction in the density of GlyR clusters opposing Renshaw cell boutons on motoneurons, as shown by super resolution microscopy. Due to the resolution limit of our STED imaging we cannot exclude that GlyR cluster area might also be altered. Decreases in recurrent inhibition were preferentially detected in large motoneurons, so it is possible that, although we targeted cell bodies >300µm^2^, our imaging results might be slightly underestimating the reduction in GlyR density. Nevertheless, we note that similar decreases in GlyR cluster density have been shown to reduce GlyR currents by ∼40% in motoneuron cultures^57^. Furthermore, the kinetics of the GlyR-mediated recurrent inhibition IPSC were slower in mSOD1 mice. These findings highlight early-stage structural and functional alterations of GlyRs clustered opposite Renshaw cell boutons in mSOD1 mice.

Interestingly, the impairment in recurrent inhibition is not a feature of all pre-motor glycinergic synapses. We found that the synaptic strength of *in vitro* group I inhibition (Ia/Ib) was largely unaffected in 2-3 week mSOD1 animals, and the kinetics of the dorsal-root evoked IPSCs were actually faster in mSOD1 mice. This indicates that early alterations in inhibitory spinal microcircuits are synapse specific.

We also found that monosynaptic Ia excitation to lower lumbar motoneurons was initially increased in early juvenile mSOD1 mice, but was reduced a few weeks later^47^, at the onset of denervation. Alterations, not only in motoneuron properties^5^, but also in spinal microcircuits^1^ have been suggested to occur prior to clear signs of weakness and substantial motoneuron death. It is possible that the need to maintain motor output within a certain physiological range leads to early-stage increases in the synaptic strength of Ia excitation so that motoneurons can produce the relatively normal output needed for behaviour. And in fact, improvement of Ia synaptic strength in young adult mSOD1 mice has been shown to have beneficial effects on biochemical disease markers^47^.

Quantal analysis revealed that an increase in the probability of release from Ia terminals to lower lumbar motoneurons is responsible for enhancement of the proprioceptive Ia afferent drive to motoneurons. Ca^2+^ influx, buffering and sensitivity determines the probability of release^58^, and pathophysiology in mSOD1 mice is characterized by Ca^2+^ dysfunction with elevated intracellular levels and mishandled Ca^2+^ buffering^59^. It is plausible that Ca^2+^ accumulation in the mSOD1 Ia terminal led to the enhanced presynaptic release, accompanied by the reduced paired pulse ratio that we observed. Interestingly, loss of Ia short-term plasticity was also reported in young adult mSOD1 animals, along with postsynaptic disruption of Ia structures^47^.

Our data show that quantal size at the Ia synapse is not altered in 2-3 week old mSOD1 mice, while post-synaptic impairment occurs in young adult mutant mice^47^, possibly because early changes in probability of release can scale postsynaptic activity by shaping the arrangement and expression of receptor subunits of AMPA receptors^44,60^. Increased tonic and/or evoked activity from Ia proprioceptive afferents to motoneurons, although perhaps initially acting as homeostatic compensation to preserve neuromuscular activity, will later act as a maladaptive plasticity event by promoting a gradual degradation of postsynaptic receptor arrangements in mSOD1 mice thus reducing the synaptic strength of Ia excitation and contributing to disease pathobiochemistry^47^ (Figure 8B).

Of note, the observed increase in Ia excitation was found preferentially in motoneurons located in more caudal lumbar segments (L5 region). ALS can spread in a cell-to-cell “domino-like” manner, with a contiguous caudal-to-rostral spinal progression of disease pathology occurring in spinal onset patients^40,61^; this is also a typical feature observed in mSOD1 mice^53^. Although the motor nuclei we studied (dorsolateral) spread evenly across L4 and L5 regions, we still detected microcircuit alterations that are segment-specific, which might reflect the nature of neurological disease progression or the order through which microcircuit homeostasis operates. Together, the changes to monosynaptic Ia excitation depend on the time-window of observation and segment studied, likely reflecting the multiphasic and multifocal features of ALS progression^61^.

### Specificity of multiphasic compensation of Renshaw cell mediated recurrent inhibition

We have shown that pathophysiology in mSOD1 mice leads to early impairment of recurrent inhibition, but no striking changes in disynaptic group I inhibition received by motoneurons, even though we could not differentiate between group Ia and Ib mediated inhibition. Although we detected a reduction in mSOD1 responses obtained from L5 motoneuron responses to L4 root stimulation, which might be an indication of the start of impairments in disynaptic Ia/Ib inhibition, it is however possible that generalized early synaptic impairments in reciprocal inhibitory circuits might be masked by the enhanced synaptic drive from Ia afferents to Ia inhibitory interneurons induced by the observed increase in release probability at Ia synapse onto motoneurons.In addition to our physiological findings, there is anatomical evidence showing that Renshaw cells and other inhibitory spinal interneurons undergo differential degenerative processes in mSOD1 mice. Studies have shown that Renshaw cells are relatively spared in mSOD1 mice with their numbers reduced only at late stages of disease progression^49,62^. On the other hand, V1 interneurons, of which Renshaw cells constitute less than 10%, with the rest being comprised of cell groups such as Ia inhibitory interneurons^63^, are reduced by 25% from the second postnatal month^48^.

It is interesting to speculate that Ia afferent synaptic drive could be a key determinant, via excitotoxic pathways, to this differential susceptibility. Although an increase in Ia afferent drive may initially be helpful in preserving interneuron function, it may contribute to progressive cell loss^48,67^. To this end, elimination of Ia afferent synapses has been previously shown to improve motoneuron survival and decrease muscle denervation, which correlated with a delayed onset of signs of motor weakness and extended survival^67^. And unlike for other V1 interneurons (e.g. Ia inhibitory interneurons), Ia contacts on Renshaw cells are sparse and immature^64–66^ – as such, they may be protected from afferent-induced excitotoxicity in mSOD1 mice.

Interestingly, we found that the initial reduction of recurrent inhibition is compensated at later stages of disease progression in mSOD1 mice. What could be the mechanism(s) underlying transient changes in recurrent inhibition? Given that the distribution of GlyR clusters at Renshaw cell-motoneuron synapses was also found to be reduced in young adult (P45) mice (Figure S7K), it is unlikely that recovery of GlyR cluster density can explain the recovery of recurrent inhibition. On the other hand, it has been shown that at ∼2-3 months of age in mSOD1 mice, Renshaw cell axons sprout, increasing their innervation of lumbar motoneurons^49^. Collateral sprouting is a typical feature of spared axons during injury/disease and can promote functional homeostasis^68^. In mSOD1 mice, sprouting seems to be relatively specific to Renshaw cells, since overall pre-motor glycinergic inputs are found to be substantially retracted from motoneurons in P45 mSOD1 mice^48^. Renshaw cell activity could be maintained by sufficient innervation by motoneurons - synaptic drive from one active motoneuron is usually enough to activate multiple Renshaw cells^34^. We suggest that reduced quantal size due to decreased GlyR cluster density is responsible for an initial impairment in recurrent inhibition in mSOD1 mice, which is then compensated by sprouting from Renshaw cells, thus leading to recovery of the function of the circuit (Figure 8A).

## Conclusion

We have identified a multiphasic time-course of homeostatic adaptations in spinal microcircuits in an animal model of progressive motor neuron degeneration: 1) increase in Ia excitation due to higher probability of release, which reverses at the onset of denervation^47^, and 2) a postsynaptic reduction of Renshaw cell-motoneuron inhibition associated with a GlyR cluster deficit, which is compensated at later stages^49^. This non-monotonic feature of microcircuit homeostasis provides new insights into disease pathophysiology, tools for following disease progression, and, ultimately, potential new avenues for circuit therapies.

## METHODS

### Animals

Experiments carried out at University College London were performed in accordance with the UK Animals (Scientific Procedures) Act 1986 and were approved by the university’s review committees and conformed to UK Home Office regulations, under project licence PPL Number: 70/9098). *In vivo* sharp electrode electrophysiological recordings performed in Paris were conducted in animals bred and housed in the BioMedTech animal facility at Université Paris Cité, with experimental procedures approved by the Paris Descartes University ethics committee (CEEA34; authorization numbers CEEA34.MM.064.12 and APAFIS ≠16338) which followed the European Directives (86/609/CEE and 2010-63-UE) and the French legislation on the protection of animals used for scientific purposes. Experiments performed at the University of Rhode Island (URI) were authorized by URI IACUC (protocol AN2021-018) and conducted in accordance with the Guide for the Care and Use of Laboratory Animals^70^. The immunohistochemical analysis were performed at Ulm University in compliance with institutional guidelines (Tierforschungszentrum, Ulm) and German animal protection laws, approved by Regierungspräsidium Tübingen (Tübingen, Germany), under license no. 1440 and o.217-9. With the exception of the procedures performed at Université Paris Cité and URI (*in vivo* motoneuron recordings), congenic C57BL/6J mSOD1 mice were used for all the experiments. C57BL/6J mSOD1 male mice (Jackson laboratory, stock N° 004435) were purchased from The Jackson Laboratory (Bar Harbor, ME, USA) and bred with non-transgenic C57BL/6J female mice, with a mating pair originating 2-3 litters from which mSOD1 males were used for further mating with non-mutant females. The congenic progeny was used for further breeding for up to 3-5 generations and the transgene copy number was constantly monitored to avoid deviation from typical ALS phenotype. The mSOD1 mice were also crossed with C57BL/6J mice that express enhanced green fluorescent protein (EGFP) under the control of the promoter of the neuronal glycine transporter GlyT2^37^, allowing the targeting of glycinergic interneurons in ALS mice. For the sharp electrode recordings performed in Paris and URI, B6SJL mSOD1 male mice (Jackson laboratory, stock N° 002726) were bred with non-transgenic B6SJL females, originating mutant mice and healthy controls on a mixed background. Although slight differences exist between congenic and mixed-background mSOD1 lines regarding disease onset and survival, these are on the timescale of days, with the time course of pathological progression being fairly similar between both models^50^.

### *In vitro* spinal cord electrophysiology

#### Oblique slices

For *in vitro* electrophysiology we used both male and female early juvenile mice (P14-25). These animals were beyond weight-bearing stage and exhibited characteristics of motor behaviors associated with adulthood such as walking, running and jumping^71^. Unlike other ALS mouse models, mSOD1 mice have no substantial sex-specific vulnerability^12,72^ and since we did not observe any differences between male and female mice in our electrophysiological data, data from both sexes are pooled together. Mice were anaesthetised with an intraperitoneal injection of a mixture of ketamine/xylazine (100 mg/kg and 10 mg/kg, respectively) and decapitated. The vertebral column was quickly extracted, pinned ventral side up in a chamber filled with ice-cold artificial cerebrospinal fluid (aCSF) of identical composition used for recordings (in mM) as follows: 113 NaCl, 3 KCl, 25 NaHCO_3_, 1 NaH_2_PO_4_, 2 CaCl_2_, 2 MgCl_2_, and 11 D-glucose continuously gassed with 95% O_2_ and 5% CO_2_. Spinal vertebrae were cut, and the spinal cord isolated from lower-thoracic to upper sacral segments. For obtaining oblique slices (Figure 2A), the cord was glued to an agar (7%) block prepared with distilled water and 0.1% methyl blue, to increase the contrast during the microscope guided slicing procedure, positioned at a 45° angle, with the ventral side with intact roots attached facing the slicing blade. This was immersed in a vibratome chamber (Leica VT1200) with ice-cold aCSF (∼2°C) comprising (in mM): 130 K-gluconate, 15 KCl, 0.05 EGTA, 20 HEPES, 25 D-glucose, 3 kynurenic acid, 2 Na-pyruvate, 3 Myo-inositol, 1 Na-L-ascorbate, pH 7.4 with NaOH^73^. Slices from L3 to L5 segments were obtained (350 µm thick), transferred to a chamber with normal extracellular solution for incubation at 37°C for 30-45 min, and were then maintained at room temperature continuously bubbled with a 95/5% O_2_/CO_2_ mixture.

#### Dorsal horn ablated and ventral horn-partially ablated longitudinal spinal cords

To obtain dorsal horn-ablated or ventral horn-partially ablated *in vitro* preparations, we glued the intact cord longitudinally to the agar with either dorsal or ventral side facing up and followed procedures recently described^35^: we aligned the vibratome blade with the midpoint between the start of the ventral commissure white matter and lower end of the central canal (for ventral horn partial ablation) or with the top of the central canal (for dorsal horn removal), and the spinal cord was slowly sectioned (0.02 mm/s). This originated a coronal spinal cord section with the dorsal or ventral side intact, containing L3-L5 segments and roots (see figure 6A and Figure S14A). The longitudinal *in vitro* preparation was then incubated in extracellular solution at 37°C for 30-45 min before being used for experiments. The spinal cord of juvenile mice is very susceptible to structural damage and anoxia, so in order to obtain viable tissue for *in vitro* recordings we relied on quick laminectomy and slicing, with the vibratome slicing commencing maximum ∼10min after decapitation^35^. All *in vitro* recordings were performed at near physiological temperature (31°C).

#### Imaging of spinal cord tissue

In oblique slices, motoneurons were clearly identifiable due to their large soma and anatomical clustering in the ventrolateral and dorsolateral regions (Figure 1A), whereas in longitudinal preparations they were distributed along the lateral rostro-caudal surface^35^. Functional identification was additionally performed, as motoneurons receive a characteristic monosynaptic excitation and disynaptic inhibition from motor efferents and sensory afferents following stimulation of ventral or dorsal roots respectively^22,35^. Putative Renshaw cells were identified by their location in the most ventral part of lamina VIII and by the expression of EGFP (Figure 3A-B). Their identity was confirmed during recordings, by the presence of an extracellular spike before establishing whole-cell mode and/or an excitatory postsynaptic current (EPSC) in response to ventral root stimulation^38,74^ (Figure 3B).

Cells were visualized using an Eclipse E600FN Nikon microscope (Nikon, Japan) containing a double port that allowed simultaneous imaging of infrared differential interface contrast (DIC) images through a digital camera (Nikon, DS-Qi1Mc), and fluorescence through either 1) a laser scanning confocal unit (D-Eclipse C1, Nikon) containing two laser lines (λ=488 and 561 nm) or 2) an epifluorescence turret (Nikon NI-FLTs) containing dichroic filters for EGFP. For epifluorescence, excitation was delivered through a 488 nm light-emitting diode (LED) (Opto LED, Cairns Instruments, UK), whose emission was detected through a charge-coupled device (CCD) camera (Retiga XR, QImaging, UK).

#### Recording setup and pipette intracellular mediums

Whole-cell recordings were performed using either an Axopatch 200B amplifier or a MultiClamp 700B (Molecular Devices). Signals were filtered at 5 kHz and acquired at 50 kHz using a Digidata 1440A A/D board (Molecular Devices) and Clampex 10 software (Molecular Devices). Glass pipettes from borosilicate thick glass (GC150F, Harvard Apparatus, UK) were pulled using a Flaming-Brown puller (P1000, Sutter Instruments, USA) and polished to a resistance of ∼1-3 MΩ for motoneuron or ∼3-4 MΩ for Renshaw cell recordings using a MF2 Narishige Microforge. Patch pipettes were filled with an intracellular solution containing (in mM) 125 K-gluconate, 6 KCl, 10 HEPES, 0.1 EGTA, 2 Mg-ATP, pH 7.3 with KOH, and osmolarity of 290 to 310 mOsm. To enable recordings of ventral root-evoked synaptic currents from Renshaw cells, we added 3 mM QX-314-Br in our glass pipettes to block any unclamped spikes evoked by antidromic motoneuron activation. To facilitate recordings of asynchronous (aIPSCs) and miniature inhibitory postsynaptic currents (mIPSCs) we used a high Cl^-^ intracellular solution with estimated reversal for Cl^-^ set at ∼0 mV that contained (in mM) 140 CsCl, 9 NaCl, 1 MgCl_2_, 10 HEPES, 1 EGTA, 4 Mg-ATP, pH 7.3 with KOH, and osmolarity of 290 to 310 mOsm. For the recordings of dorsal root-evoked inward and outward synaptic currents from motoneurons from ventral horn-partially ablated longitudinal preparations, we used a Cs-gluconate based intracellular solution containing (in mM) 125 Cs-gluconate, 4 NaCl, 0.5 CaCl_2_, 5 EGTA, 10 HEPES, 2 Mg-ATP, pH 7.3 with CsOH, and osmolarity of 290 to 310 mOsm. Although the use of Cs-gluconate precludes the measuring of firing properties, it improves space-clamp when in voltage clamp mode, which largely benefits the recordings of the large sensory-evoked currents (∼2-10 nA) from mature motoneurons, especially when holding motoneurons at the reversal potential for excitatory currents^35^.

#### Motoneuron electrophysiological properties

Motoneuron capacitance and resistance were estimated either in current clamp, from the voltage change to a brief (100 ms) current step (50 to 200 pA), or in voltage clamp, from the current response to a voltage step (5 mV). Whenever possible, motoneurons were distinguished based on their initial firing profile (Figure 1B) following injection of 4 sec-long increasing steps of current until they started to spike. Cells exhibiting early firing are strongly associated with smaller, putative slow motoneurons whereas delayed firing responses are associated with larger motoneurons that innervate fast motor units^30^. From the responses to these increasing steps of current (Figure 1C) we were able to extract further information on motoneuron firing output: membrane potential represents the resting voltage before any current injection; rheobase is reported as the current step at which repetitive action potentials were first observed; maximum sustained firing frequency is the averaged instantaneous firing frequency from the last 2 seconds of the step that elicited the fastest repetitive firing until the end of the current step; 1-2^nd^ spike interval is the instantaneous firing frequency between the first and second spikes from the step that elicited the fastest repetitive firing frequency; and depolarization block was defined as the current measured from the first step in which motoneurons stopped firing repetitively during the 4 sec-long pulse.

From the initial 2-3 evoked spikes from the step that triggered initial firing, we extracted additional information on action potential kinetics (Figure 3D-E): threshold was estimated as the voltage at which the derivative of the action potential reaches 20 mV/ms; spike amplitude was considered as the difference between threshold and peak and we also considered spike half-width, rise and decay times measured between 10-90% of amplitude, and maximum repolarization and depolarization rates; the fast afterhyperpolarization phase (fAHP) was taken as the difference between the threshold and the end of the repolarization phase; afterdepolarization phase (ADP) was measured as the voltage amplitude between the peak of the fAHP and the most positive voltage value that immediately follows the end of the repolarization phase. In some instances the initial spikes were distanced enough to allow the measurement of the duration of the medium afterhyperpolarization phase (mAHP; Figure 3F) during the current step injection, a much slower AHP lasting several milliseconds, whose amplitude and half-width were estimated from the most negative point to a stable baseline value similar to the pre-spike voltage.

#### Microcircuit electrophysiology

To study efferent and afferent-related microcircuits, ventral and dorsal roots were stimulated via a suction electrode with the tip cut to match the thickness and length of the root. An isolated constant current stimulator (DS3, Digitimer, UK) was used to stimulate the roots at an intensity fixed at 3-5x (for ventral roots) or 1-3x (for dorsal roots) the threshold for evoking an initial synaptic response in the recorded cell. The intensity of the dorsal-root stimulation is adjusted to preferentially recruit the thickest nerve fibers (group I afferents) and therefore obtain responses associated with monosynaptic (Ia) excitation and disynaptic (Ia/Ib) inhibition^35^. For root responses in longitudinal preparations, usually both L4 and L5 responses were obtained from each motoneuron when possible. For recordings of aIPSCs, the strength of the ventral root stimulation was adjusted in each experiment (∼1-3x threshold) to allow the clear identification of individual asynchronous release events (Figure 4F). For analysis and representative figures of root-evoked excitatory (EPSCs) or inhibitory postsynaptic currents (IPSCs), traces were baselined and a single or double exponential was used to correct for the stimulation artefact^22^. The conductances (σ) of the root-evoked excitatory (EPSCs) or inhibitory postsynaptic currents (IPSCs) were calculated at the holding voltage assuming a reversal of 0 mV for excitatory and -60 for inhibitory conductances (except for high Cl^-^ intracellular solutions) taking into account a correction for the junction potential (∼15 mV for all intracellular solutions):

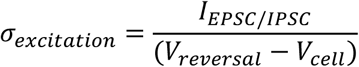

Holding voltage was usually -60 mV for EPSCs and 0 mV for IPSCs recordings, with series resistances in the range of 2 to 10 MΩ compensated by 40% to 80%. Given the range of measured capacitances in motoneurons, the amount of uncompensated series resistance gives rise to a filtering cutoff frequency in the range 0.04-9 kHz. . In some cases cells were hyperpolarized to prevent action antidromic or orthodromic action potentials. In addition to recording near the equilibrium for Cl^-^, for most excitatory ventral root-evoked responses and all dorsal root-evoked currents obtained for Bayesian quantal analysis (BQA), EPSCs were recorded in the presence of strychnine (1 µM) and gabazine (3 µM). For root-evoked EPSCs and IPSCs, in addition to the synaptic conductance, we also extracted information on the rise and decay phases taken from 10-90% of their amplitude. For dorsal root responses, stimulation was performed at 33 Hz in order to measure the paired-pulse ratio of evoked responses (Figures S10A and E). In voltage-clamp recordings, the tail of the stimulation artefact will contaminate the onset of the neuron’s response to dorsal or ventral root stimulation. To remove this contamination, we fitted the end of the artefact with a double or single exponential function, which was then subtracted from the recorded trace. The root responses were obtained from 2-4 week old mature animals at 31°C, leading to responses ∼50% faster than those obtained at room temperature^35^, which may give the impression that both monosynaptic and disynaptic root responses have similar jitter. We performed latency and jitter comparisons between dorsal root-evoked EPSCs and IPSCs obtained from the same cell and root, and for this we considered the onset of the synaptic response as the time-point after the root stimulation in which the derivative of the response reached 5 times the standard deviation of the baseline noise (Figure S8C). Only responses in which a double exponential was used to correct for the stimulation artefact without introducing substantial noise to the derivative were used for comparisons, which revealed that dorsal root-evoked disynaptic IPSCs have larger jitter and latency than monosynaptic EPSCs^35^ (Figure S8D-F). Given that many factors can affect the comparison of latencies between neurons (e.g. length and stretch of root, suction electrode, stimulation artefact), we did not make latency and jitter comparisons between cells and experimental groups.

For this study we tried to obtain ∼10 or more stable responses for analysis, with individual values averaged and reported per cell. Since motoneurons have variable conductance and thus the amount of inputs received might be proportional to their cell conductance, absolute synaptic conductances were scaled and shown as a percentage relative to the resting conductance of the recorded cell:

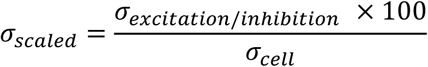

In the absence of differences in resting input conductance between experimental groups, we considered clear alterations in both absolute and scaled conductance as support for differences between mSOD1 and WT mice. For simplicity, plots for scaled conductances are represented in the supplementary figures and tables.

In order to study recurrent inhibition we used an approach that 1) would allow to measure motoneuron firing output and thus characterize its initial firing profile, and 2) could be employed for both *in vivo* and *in vitro* motoneuron recordings. Recurrent inhibition was measured in the presence of D-2-amino-5-phosphonopentanoic acid (APV, 50 µM), 1,2,3,4-tetrahydrobenzo(f)quinoxaline-7-sulphonamide (NBQX, 3 µM) and gabazine (3 µM), which is sufficient to completely block recurrent excitation without affecting the contribution of Renshaw cells drive to motoneurons^22^. We initially estimated resting cell conductance through a current-voltage relationship obtained by injecting a series of steps of current at resting voltage, which was then subtracted from the conductance calculated during ventral root stimulation (100 ms) at a frequency of 200 Hz in which recurrent inhibition reaches a steady-state voltage (Figure 2E). The contribution of Renshaw cells to motoneuron conductance during high-frequency ventral root stimulation is maximum, thus the difference between the conductance during this period and motoneuron resting conductance was considered as measurement of the strength of recurrent inhibition. This method has been recently used, and was shown to be as efficient at estimating recurrent inhibition as the measurement of absolute synaptic currents in voltage clamp^35^. When measuring inhibition evoked by dorsal root stimulation, this method was not suitable, because it requires blockade of glutamate receptors in order to isolate the disynaptic inhibitory input from the monosynaptic excitation, which would have also blocked direct transmission to the Ia/b interneurons that are responsible for the disynaptic inhibition to motoneurons. We therefore used a Cs-gluconate based intracellular solution in order to clamp motoneurons at the reversal for excitation and inhibition and measure monosynaptic Ia excitation and disynaptic Ia/Ib inhibition respectively.

#### Bayesian quantal analysis (BQA) implementation with improved discrete-grid exact inference

BQA was used to estimate quantal parameters from motoneuron-Renshaw, Renshaw-motoneuron and Ia afferent-motoneuron synapses. BQA provides estimates for number of release sites (n) and quantal size (q), by modelling the amplitude distribution of responses at all observed probabilities of release^75^. In the present study the discrete-grid exact inference implementation of BQA was improved to confer increased computational robustness for the estimation of quantal parameters particularly wherein numbers of evoked responses exhibited considerable variation recorded at different release probabilities. Source code for the improved method is available for download at https://github.com/Bhumbra/Pyclamp, and a detailed description of the new implementation is provided in supplemental information.

Root-evoked currents or potentials were recorded in the presence of different concentrations of extracellular Ca^2+^ (1, 2 or 4 mM) in order to modulate the release probability. To estimate the quantal parameters at Renshaw-motoneuron synapses, we decided not to perform acquisition of ventral root-evoked IPSCs in voltage clamp as these would prove challenging since it would require clamping large currents (several nA in amplitude) at a holding voltage that would maximize the electromotive force for Cl^-^ (∼0 mV), but that is far from the resting membrane potential. Although for BQA on Renshaw-motoneuron connections (in neonates) we have previously used non-saturating concentrations of strychnine in addition to a high intracellular Cl^-^ based solution^74^ in order to reduce the size of the evoked currents, employing such strategy would affect the quantal size estimate (q) which would not be comparable between groups. Therefore, to facilitate the BQA experiments for the estimation of the quantal parameters at Renshaw-motoneuron synapses in oblique slices (Figure 4A), we recorded stable ventral root-evoked inhibitory postsynaptic potentials (IPSPs) slightly above the reversal for Cl^-^ (figure 5a), and corrected for fluctuations in the membrane potential by applying the correction factor below to the IPSP amplitude in each sweep:

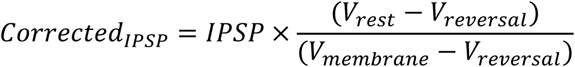

Where V_reversal_ (-75 mV) was calculated from the Cl^-^ equilibrium potential imposed by our intra-and extra-cellular solutions, V_rest_ is -60mV and V_membrane_ is the membrane potential at which the recordings were obtained.

For motoneuron-Renshaw BQA we recorded root-evoked EPSPs obtained at resting voltage (∼-60 mV), and therefore for quantal estimation we did not add any correction factor since the resting membrane is far away from the reversal for excitation (figure 3E). BQA for Ia afferent-motoneuron was performed in voltage clamp in the presence of gabazine and strychnine, and at holding potentials close to resting membrane potential usually ranging between -60-70mV (Figure 6A). If series resistance increased by >10% experiments were stopped.

#### Asynchronous and spontaneous synaptic release

To obtain and quantify aIPSCs derived from Renshaw cells, we replaced extracellular concentrations of Ca^2+^ with Sr^2+^, which is known to desynchronize and prolong the duration of presynaptic neurotransmitter release^39^. In the presence of 4mM of Sr^2+^, we were able to measure asynchronous events that immediately followed evoked current responses to repetitive (200Hz) ventral root stimulation^76^ in the presence of NBQX, APV and gabazine (Figure 4F). For each motoneuron we determined the amplitude of ∼100 glycinergic aIPSCs acquired at holding voltages close to -60mV. Prior to the replacement of extracellular Ca^2+^ with Sr^2+^, we recorded individual ventral root-evoked IPSCs and estimated their rise and decay times measured between 10-90% of their amplitude (Figure S5). Recordings of glycinergic mIPSCs (∼200-1000 per motoneuron) were performed in gap-free mode with NBQX, APV and gabazine at holding voltages of or near -60mV, and we considered their amplitude, inter-event interval and rise and decay times taken from 20-80% of their amplitude. Unlike root evoked currents whose signal is several magnitudes larger than the background noise and for which 10-90% amplitude is a good and less conservative proxy for estimating EPSC/IPSC kinetics, mIPSCs are quite small in amplitude and thus can be affected by baseline noise and therefore 20-80% is a more robust gauge for estimating rise and decay times. For mIPSCs analysis we defined the detection criteria as any spontaneous event larger than 1.5x the baseline noise with a waveform characteristic of a synaptic event (rise phase >0.1ms and decay >0.5ms).

### *In vivo* electrophysiology

#### Motoneuron intracellular recordings

Procedures were performed as previously described^47^ with minor adjustments. Mice were anesthetized with a mixture of Fentanyl/Midazolam/Dormitor (dose 50 µg/kg / 5 mg/kg / 0.5 mg/kg; maintenance 5 µg/kg / 0.5 mg/kg / 50 ug/kg). In these experiments, the dorsal S1-L4 roots were cut proximally to the spinal cord to remove any source of sensory-driven excitation onto motoneurons, and the ventral roots L6, L5 and L4 were cut as distally as possible, dissected free and placed on individual bipolar stimulation electrodes. After impalement, motoneurons were identified as belonging to one of the roots through their antidromic action potential, then, one of the adjacent roots was stimulated at the intensity that elicited the maximum recurrent inhibition (Figure 7A).

Recordings were performed using glass micropipettes filled with K-acetate 2M (resistance 20-30 MΩ). In all cases, the presence of inhibition was visually confirmed in the recording, which sometimes required the injection of bias current solely for that purpose. To estimate the inhibitory conductance, we compared the total conductance of the motoneuron in control conditions (at rest) to the total conductance during the steady-state recurrent inhibitory post-synaptic potential, as previously described for *in vitro* electrophysiology^35^. The control conductance was measured using a series of 50 ms current pulses (∼1nA) 300 ms prior to nerve stimulation. Then, the root was stimulated with 20 shocks at 200 Hz to elicit recurrent inhibition. A second series of current pulses with the same duration and amplitude were delivered 40 ms after the first shock. Five to 10 sweeps were averaged for each current intensity, and a current-voltage curve was obtained by plotting the intensity of the current pulse against the control and test voltage deflection.

In our *in vivo* recordings, excitatory postsynaptic potentials were not detected following ventral root stimulation, which led only to recurrent inhibitory potentials. It is possible that general anaesthesia depresses recurrent excitation, which has been reported in adult chloralose cats^77^, or that in mice, these synapses are developmentally depressed beyond the first postnatal month, or that the recurrent excitation is masked in our recordings by the large shunting due to recurrent inhibition. As such, we focused solely on Renshaw inhibition in our *in vivo* studies.

#### Electromyographic (EMG) recordings

Mice were anaesthetized with an intraperitoneal injection of a mixture of ketamine and xylazine (100 mg/kg and 10 mg/kg, respectively) and tested for withdrawal reflex before proceeding with EMG recordings. Approximately one-sixth of the initial dose of ketamine and xylazine mixture was injected as required during the experimental procedures to supplement anaesthesia. We performed experiments in both male and female mice organized into three different groups: P18-23 (similar age to *in vitro* recordings), P57-67 (adult mice without clear signs of muscle weakness and of similar age as those used for *in vivo* recordings) and P87-108 (onset of signs of motor weakness).

Either the left or right skin of the hindlimb was shaved and eye ointment was applied on the eyes to prevent drying during the procedure. The experimental mouse was placed on a heating pad inside a homemade faraday cage, with internal (∼36°C) and skin temperature (∼31°C) monitored. The animal was grounded to the cage from the base of the tail. Two multi-stranded perfluoroalkoxy-coated stainless steel fine (25.4 µm strand diameter) wires (A-M Systems, USA) with their tip peeled to increase the active surface for recording, were inserted in the tibialis anterior (TA) (slightly lateral to the tibia and just distal to the patella) and quadriceps (Q) (middle portion – rectus femoris) muscles using a 25 G needle. The tip of the wires was fish-hooked before insertion to stabilise the filaments once inserted into the belly of the muscle. For peripheral nerve stimulation, two fine wires were positioned subcutaneously with a 25 G needle to 1) an area slightly proximal to popliteal fossa and mid-thigh for sciatic nerve stimulation and 2) the anteromedial aspect of the femur for femoral nerve stimulation (Figure 6D). The specificity of the stimulation was tested by checking the occurrence of knee extension following femoral nerve and knee flexion following sciatic nerve suprathreshold stimulations, along with the presence of specific direct motor response (M-response) in Q and TA muscles after small stimulus intensities (∼1-1.5x threshold intensity). Peripheral nerves were stimulated with a DS3 constant current stimulator (Digitimer, UK), bipolar EMG signals were obtained using a EXT-02F extracellular amplifier (NPi Electronic, Germany), filtered at 5 kHz and acquired at 20 kHz using a CED Power1401 using Spike 2 v8 software (Cambridge Electronic Design, UK).

Maximum M-responses, that were considered as the largest direct muscle responses induced by orthodromic nerve stimulation (M_max_) for both TA and Q muscles, were found using a 200 µs-long square pulse, whereas H-reflex responses were recorded from TA following sciatic nerve stimulation with 1 ms-long square pulses. Shorter pulse durations such as 200 µs can be used to minimize H-reflex contamination throughout the experimental procedure^78^. If the maximum H-reflex was smaller than 1 mV and/or threshold intensity for clear H-reflex was above 1 mA, experiment was stopped and discarded. After confirming the M_max_ in both muscles and the detection of H-reflex in TA, the experimental stimulus intensities were arranged, so that the TA H-reflex was around half of maximal H-reflex (H_max_) and Q M-response was 10% of M_max_ (in order to further minimise the stimulation of sensory fibres, see next paragraph). In each experiment lack of Q H-reflex was also confirmed before beginning any protocol.

To study recurrent inhibition, we used a method that tests heteronymous recurrent inhibition between Q and TA^79^. This is done by measuring the depression of TA H-reflex induced by a conditioning Q M-response that produces antidromic activation of Renshaw cells. Although studies using this method have claimed that the effect of sensory fibre stimulation (possible Ia inhibition) in their conditioning stimulus is minimal^55,79,80^, we set the intensity of femoral nerve stimulation to 10% of Q M_max_ and stimulus duration to 200 µs to favour stimulation of motor axons^78^. In addition, in our experimental setting, mice have been anaesthetized with ketamine which can act as a α-amino-3-hydroxy-5-methyl-4-isoxazolepropionic (AMPA) receptor blocker^81^ and is known to depress the H-reflex without affecting the M-response^82^, thus further minimizing the possible effect of small intensity and short duration conditioning on the recruitment of sensory afferents. For these experiments we obtained a single pulse control TA H-reflex, followed by a conditioning Q M-response 2, 5, 10, 30, 50 and 100 ms prior to a test H-response (Figure 7C). To study post-activation depression of H-reflex, we performed paired pulse stimulation of the sciatic nerve to evoke an initial control H-reflex followed by a second test H-reflex triggered 50 ms, 100 ms, 200 ms, 500 ms, 1 sec, 5 sec or 10 sec after the first H-response (Figure S15A). For all experiments we used a 20 sec interval between control and test H-reflex, regardless of any protocol, to prevent the effect of post-activation depression in our measurement. Each interval was randomly selected and repeated at least five times. Because of the variable nature of the H-reflex and possible effect of anaesthesia on it over time, we scaled each test H-reflex to its own control H-reflex, averaged these scaled responses for each interval and pooled mean values for each animal.

#### Immunohistochemistry and STED imaging

Immunohistochemistry was performed as previously reported^47^. Briefly, mice (both at P21 and P45) were, euthanized by cervical dislocation and were transcardially perfused with 50 mL ice-cold phosphate-buffered saline (PBS) followed by 2.5-3mL/g of freshly-prepared (<24h) 4% paraformaldehyde (PFA) in PBS. Upon dissection, spinal cords were post-fixed in 4% PFA/PBS for 18h and cryoprotected in 30% Sucrose/PBS for 36h, freeze-embedded in optimal-temperature-cutting (OCT, TissueTek) mix and sectioned at -18°C in a cryostat (Leica CM1950); 20µm sections were obtained across the lumbar spinal cord. Free-floating sections from L3-L5 metamers were blocked in 3% bovine serum albumin (BSA)/0.3 % Triton X-100 /PBS for 2h and then incubated (50h at 4°C on a rotary shaker) with the following primary antibodies (diluted in blocking buffer): chicken anti Calbindin (Calb, 1:500, Invitrogen PA5-143561), rabbit anti GlyT2 (1;500, Synaptic Systems 272 003), rat anti vesicular acetylcholine transporter (VAChT, 1:300, Synaptic Systems 139 017) and mouse anti glycine receptor (GlyR) alpha-1 (1:200, Synaptic Systems 272 003). Sections were thereafter washed 3x 45 min in 0.1% Triton X-100/PBS and incubated (2h, 24 °C, rotary shaker) in the following secondary antibodies: donkey anti rat 405 (Invitrogen) donkey anti chicken Cf488A (Sigma) FluoTag-X4 anti mouse aberrior star 580 and FluoTag-X4 anti rabbit aberrior star 635 (Nanotag). Sections were mounted in ProLong Gold Antifade (Thermo Fisher Scientific) and dried at room temperature for 24h before the imaging.

Super-resolution stimulated emission depletion (STED) imaging was performed on a Stedycon module (Abberior instruments, Germany) fitted to a Zeiss microscope with a 100× (NA 1.4) oil objective. Images were acquired at 8-bit depth in both confocal and STED modes; to limit photobleaching, only the GlyR channel was imaged in STED mode (whereas the Calb, GlyT2 and VAChT channels were only imaged in confocal mode). Motoneurons were identified by the moderate cytoplasmic VAChT staining and by the intense staining of C-boutons surrounding the cell body and proximal dendrites (the presence of C-boutons allows the exclusion of gamma-motoneurons). In line with our *in vitro* observations on recurrent inhibition, we preferentially targeted large (>300µm^2^) ventrolateral lumbar motoneurons. Imaging parameters in the confocal and STED modes were set to avoiding saturation. The laser power for the excitation laser was set to 20% for the 580 nm channel, with the depletion laser power set at 60% of the maximum output. Single optical sections crossing through the maximum diameter of each motoneurons were recorded at 5-8µm depth within the section, avoiding surface staining artefacts and reducing depth-and scattering-dependent variability in staining and fluorescence intensity.

For the quantification of the number of GlyT2^+^/Calb^+^ presynaptic terminals, confocal images were used. Processing was performed in ImageJ (National Institutes of Health, Bethesda, Maryland): confocal stacks were subject to rolling-ball background subtraction (50-pixels diameter) in all channels and then de-speckled. Confocal stacks composed of 7-8 optical sections spanning the maximum diameter of the motoneuron (including the nuclear shadow) were collapsed (maximum-intensity) and subject to a watershed filter to isolate different presynaptic terminal. The perimeter of the motoneuron was manually outlined and measured, and each presynaptic terminal was manually identified; the density of presynaptic terminals was calculated as number of terminals per 100µm of motoneuron perimeter.

For the quantification of the size and number of GlyR clusters, single-optical section STED images were subject to background subtraction (rolling-ball, 20-pixels diameter), de-speckled and thresholded at 1.5 times the background level (e.g. with a minimum signal-to-noise ratio of 2). GlyR clusters located on the inner or on the outer border of the non-STED-resolved GlyT2^+^/Calb^+^ double-positive presynaptic terminals were counted and quantified. Since the presynaptic terminals were imaged only in confocal mode (to avoid photobleaching), a complete separation of pre-and post-synaptic structures was not achieved and therefore some GlyR clusters appear inside the area of the presynaptic terminal.

For image analysis, healthy control and mSOD1 mice were processed in pairs and therefore this was also considered as an additional level in the hierarchical structure of the data obtained.

#### Data and statistical analysis

The analysis of mIPSCs raw traces and BQA from *in vitro* recordings were performed with PyClampsoftware (http://github.com/Bhumbra/Pyclamp), and the remaining *in vitro* raw data were analysed with Clampift 10.7 (Molecular Devices, USA). Spike 2 v8 was used for the analysis of the raw EMG and *in vivo* motoneuron recordings data. Statistical analyses were done using OriginPro 2022 (OriginLab Corporation, USA), Microsoft Excel version 2208 (Microsoft, USA), MATLAB R2022b (Mathworks, USA) and R Studio version 2022.07.1 (R Core Team, Austria). Statistical plots and figures were generated with Origin Pro 2022 and Microsoft PowerPoint version 2208 (Microsoft, USA).

The majority of the data produced in this study have a hierarchical structure, that is electrophysiological observations were taken from cells that were recorded from different animals, or several synaptic inputs (aIPSCs and mIPSCs) were taken from individual motoneurons (>50 per cell). This is a common feature in neuroscience datasets, and therefore data dependencies must be taken into consideration when analysing and reporting results to minimize the risk of false positives^35,83–85^. In order to decide on how to appropriately analyse and represent the data collected in this work, we first sought out to identify the variance component in our datasets. The intraclass correlation coefficient (ICC) provides a good measurement of reliability and partition of the sources of variability in our datasets^84,86,87^. For our data structure, we used the one-way random effects ICC model (1,1), which can be estimated through the mean squares (*MS*)^87,88^:

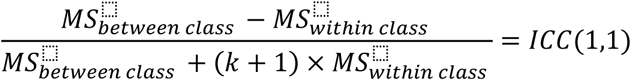

To calculate the ICC we used the function *ICCbare* from the *ICC* R package^89^. The output is generally a value between 0 and 1, and can be interpreted as an indicative of poor reliability if ≤0.50, moderate if between 0.50 and 0.75, good if between 0.75 and 0.90 and excellent if >0.90 ^87^. If ICC was 0.50 or less, meaning that 50% of more of the variance was within class and thus explained by the lowest level observation (e.g. motoneuron properties or synaptic currents), we treated those data as independent. In such cases, to make comparisons between groups, we computed two different effect sizes: 1) bootstrapped mean difference and 2) bootstrapped Hedges’ *g*. Briefly, random *n* values are resampled with replacement from each group (WT or mSOD1) and their means (µ) compared, or used to estimated Hedges’ *g* as follows:

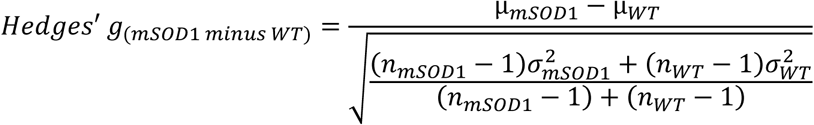

With *n* being the original sample size from each group and 10,000 bootstrap replicas were performed in total (paired resampling used for jitter and latency analyses). From the computed bootstrap we considered the resampling distributions, 95% confidence interval (CI) and mean. Group comparisons in which the 95% CI of the effect size did not include 0, would be interpreted as statistically meaningful. While mean difference permits to infer about absolute unitary changes between groups, it can be influenced by sample size, whereas Hedges’ *g* is less sensitive to it and provides a coefficient indicating by how many standard deviations the groups differ^90^. Guidelines for Hedges’ *g* refer to small, medium and large effects as 0.20, 0.50 and 0.80, respectively^90^. The interpretation of these benchmark values cannot be strict^91,92^ and results from Hedges’ *g* will be discussed appropriately taking also into consideration the mean difference effect size and the nature of biological variable studied.

Unsurprisingly, we found that for the majority of the data in this study ICC was smaller than 0.50, which probably reflects the high heterogenous properties of motoneurons^35^. But for datasets in which ICC was >0.50 we considered either a hierarchical bootstrap or a linear mixed-model (LMM). The hierarchical bootstrap has a smaller false negative rate than LMMs and is a more adequate statistical method for nested data in which the number of observations is large^93^. This method would be appropriate for the analysis of synaptic inputs (mIPSCs and aIPSCs), for which the ICC was >0.50 and the number of observations per motoneuron were ∼100 or more. However, since for mIPSCs and aIPSCs the number of total observations is in the thousands, and the number of observations per cell greatly varies (in some cases ranging between 100 and 1000), we decided to implement the hierarchical bootstrap for aIPSCs and mIPSCs analyses independently of the ICC value. We performed a two level resampling -1^st^ level - motoneuron and 2^nd^ level - aIPSCs or mIPSCs - where for each sampling replica (10,000 in total) we extracted *n* motoneurons with replacement followed by *k* aIPSCs or mIPSCs from each cell, with *n* defined as the total number of recorded cells per group and *k* as the maximum number of inputs obtained per cell in each group. From each hierarchical bootstrap replica we then computed the mean difference and Hedges’ *g* like previously described.

Hierarchical bootstraps are not ideal for datasets with small number of observations per subject since the resampling would not accurately represent the population distribution^93^, and therefore for the remaining data in which ICC was >0.50 and observations were less than 10-20 per mouse we employed a LMM. This was only the case for some recordings obtained from early firing motoneurons, Renshaw cells BQA and some dorsal root-evoked synaptic currents. We fitted a LMM with a fixed-effect coefficient for genotype (WT or mSOD1) and a random intercept that varies by animal. We used the *lmer* function within the package *lme4* in R ^94^ to fit the model as follows:

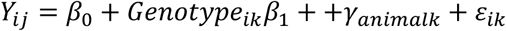

Y_ijk_ represents the datapoint pertaining to the i^th^ observation obtained from the k^th^ animal; β_0_ is the intercept; Genotype_ik_ is the predictor for observation i within k and β_1_ is its coefficient; γ_animalk_ is the random effect term and ε_ik_ is the residual error. From the *lmer* output we reported the predicted value for the WT group (intercept) and the estimated difference for the mSOD1 group along with respective 95% CIs. The variance of the random effects is also shown and used to estimate the ICC from the LMM. We have also reported partial eta squared (η_p_^2^) and its 90% CI,^95^, calculated with the *t_to_eta2* function in the *effectsize* R package using the *tback* method^96^. As an effect size, although slightly conservative for LMM, the η_p_^2^ output can be generally interpreted as small (0.01), medium (0.06) and large (0.14)^97,98^. For most of the data in which we employed a LMM, a considerable percentage of animals only had one single observation which may have led to inappropriate fitting of the model thus possibly affecting any inferences about the biologically relevance of the data. Therefore, in addition to the results from the LMM, we decided to also treat datapoints relating to those data as independent and report bootstrapped mean difference and Hedges’ *g.* Any outstanding differences between the LMM output and mean difference and Hedges’ *g* effect sizes are discussed in detail.

The data pertaining to the GlyR cluster immunohistochemistry contain thousands of observations within a hierarchical structure with numerous levels (e.g. bouton, motoneuron, animal, experimental pair id) and therefore we decided to employ a LMM in order to take into consideration the multiple different random effects. For cases in which most of the variability was within observations (i.e. ICC Residuals ≥0.50) we also report appropriate bootstrapped effect sizes considering observations as independent.

To infer about correlations between cell intrinsic properties and root-evoked synaptic conductances, we performed a non-linear Spearman rank test to explore if synaptic strength correlates directly or indirectly with cell size and/or conductance. The Spearman rank coefficient (ρ) is a good indicator of the strength of the relationship and values between 0.40 and 0.69 are associated with a moderate correlation whereas those larger than 0.7 are interpreted as strong correlations^99^.

For the *in vivo* and *in vitro* data, all datapoints are shown on the left side of top panel next to respective box-plots depicting the minimum, first quartile, median, third quartile, and maximum value for each group. On the right side of the top panel, we have depicted respective bootstrapped mean difference distribution (Kernel Smooth filled curve), mean (dot) and 95% CI (whiskers), with ‘0’ value aligned with the mean of the WT group and predicted mean difference with the mSOD1 group by dotted horizontal lines. In cases in which data dispersion required the use of a log_10_ scale, the bootstrapped mean difference effect size had to be shifted, with the ‘0’ value now centered on the mean of the WT group (‘Mean difference centered’). On a bottom panel we have represented bootstrapped Hedges’ *g* distribution (Kernel Smooth filled curve), mean (dot) and 95% CI (whiskers). For easier interpretation of the data pertaining to the 18 motoneuron properties we analysed, for Figure 1, mean bootstrapped Hedges *g*’ effect size is represented in a heatmap format, with the parameters for which 95% CI did not cross ‘0’ being highlighted. For the EMG data, both bootstrapped mean difference and Hedges’ *g* are shown below the data plots. To better interpret the EMG results on recurrent inhibition and post-activation depression, data are shown as connecting line series plots with each dot representing the mean for each interval and the shaded area the first and third quartiles. For the GlyR cluster data we displayed the LMM effect size estimate adjacent to the plot with respective η_p_^2^ and 90% CI on top, and due to the high number of observations and for better visual interpretation of the data, we also added density plots (Kernel smooth filled curves) and in some cases a log base 10 (log_10_) was used to transform data. Correlation graphics are shown as scatter plots depicting all the individual points plus linear regression lines with 95% CIs (shaded area) for illustration purposes; Density plots (Kernel smooth or lognormal filled curves) of the data are also shown in the margins. Traditional descriptive statistics such as the absolute mean±standard deviation (SD) plus number of observations (n) and mice are also reported throughout the article and supporting tables.

## Acknowledgements

We would like to thank Pascal Branchereau and Hongmei Zhu for preliminary efforts and discussions on GlyR cluster imaging. F.N. was supported by a Sir Henry Wellcome Postdoctoral Fellowship 221610/Z/20/Z; M.G.O. was supported by Royal Society Newton International Fellowship NIF\R1\192316; F.R. was supported by the Deutsche Forschungsgemeinschaft (in the context of the ANR/DFG cooperation program) with the grant no. 446067541 and with the grant no. 431995586 (individual grant). F.R. was also supported by the German Center for Neurodegenerative Diseases-Ulm. M.M. and D.Z. were supported by National Institutes of Health, Institute of Neurological Disorders and Stroke (R01NS110953) and the Fondation Thierry Latran (SPIN-ALS project). D.Z. was also supported by the Agence Nationale de la Recherche (in the context of the ANR/DFG cooperation program) with the grant no. ANR-20-CE92-0029-01). R.M.B. was supported by Medical Research Council Research Grant MR/V003607/1; M.B. was funded by Biotechnology and Biological Sciences Research Council Research Grant BB/S005943/1, M. Bą. Was supported by the Polish National Science Centre Grant 2017/26/D/NZ7/00728

