## Supplemental figures, tables and appendix for "Spinal microcircuits go through multiphasic homeostatic compensations in a mouse model of motoneuron degeneration"

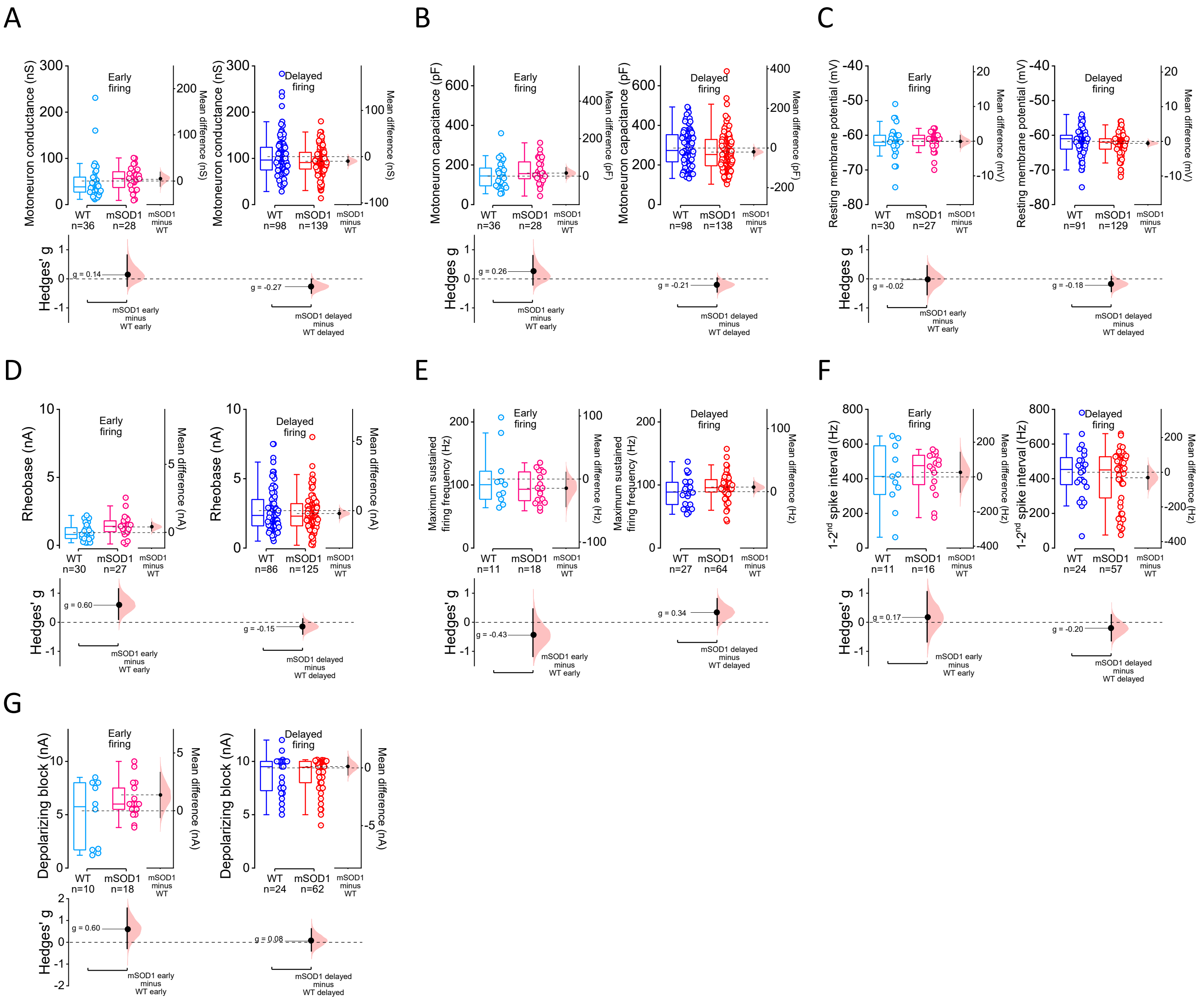

**Figure S1 – Motoneuron subthreshold and repetitive firing properties from early juvenile WT and mSOD1 mice.** Group data for **(A)** resting input conductance, **(B)** capacitance, **(C)** membrane potential, **(D)** rheobase, **(E)** maximum firing frequency, **(F)** 1-2$^{nd}$ spike interval and **(G)** depolarizing block. Estimation plots with all individual values and respective box-plots shown along with respective bootstrapped mean difference and bootstrapped Hedges' *g***.** See also Table S1.

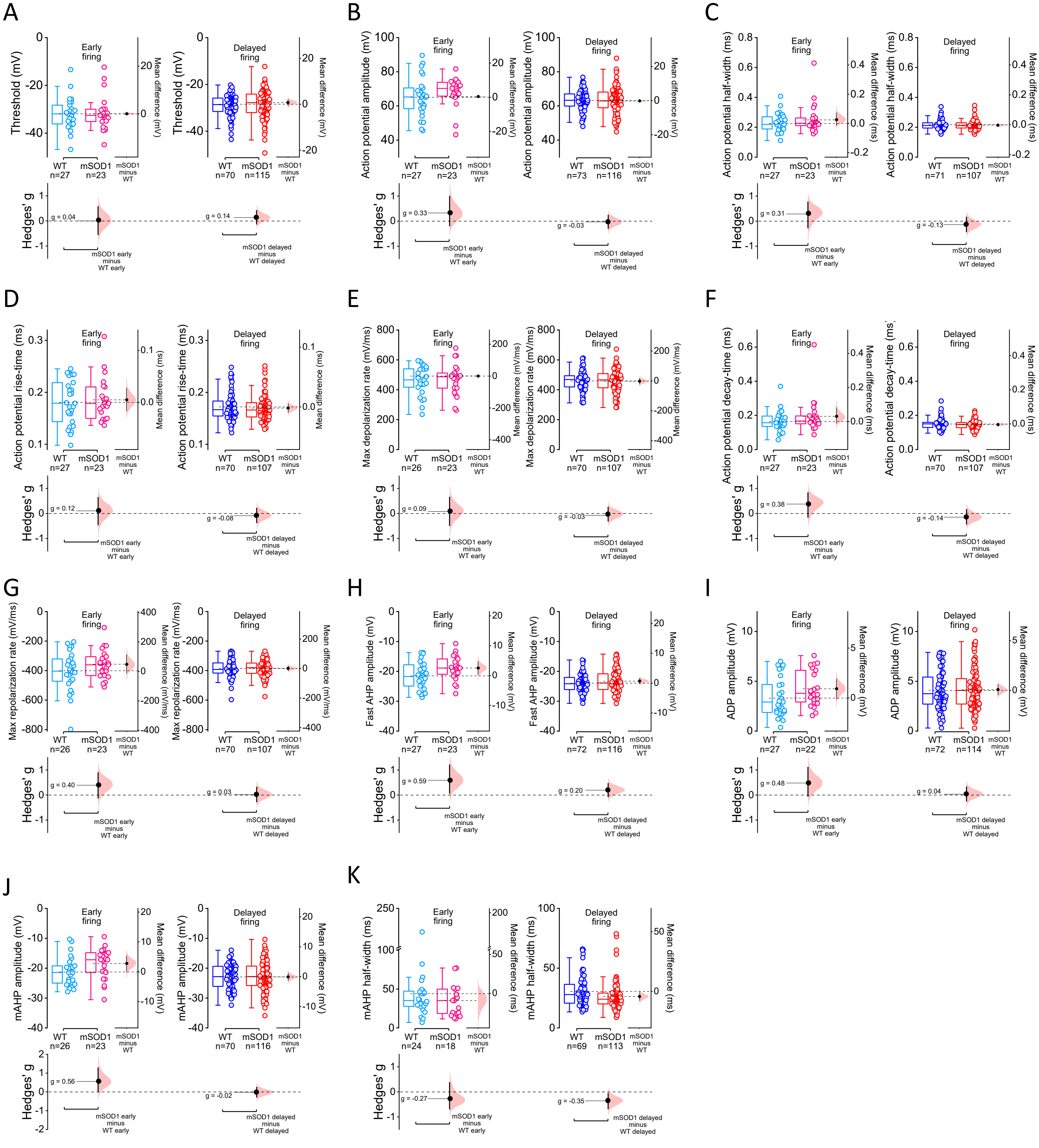

**Figure S2 – Motoneuron action potential properties from early juvenile WT and mSOD1 mice.** Plots for action potential **(A)** threshold, **(B)** amplitude, **(C)** half-width, **(D)** rise time, **(E)** maximum depolarization rate, **(F)** decay time, **(G)** maximum repolarization rate, **(H)** fast afterhyperpolarization (AHP) amplitude, **(I)** afterdepolarization (ADP) amplitude and medium AHP **(J)** amplitude and **(K)** half-width. Estimation plots with all individual values and respective box-plots shown along with respective bootstrapped mean difference and bootstrapped Hedges' g. See also Table S2.

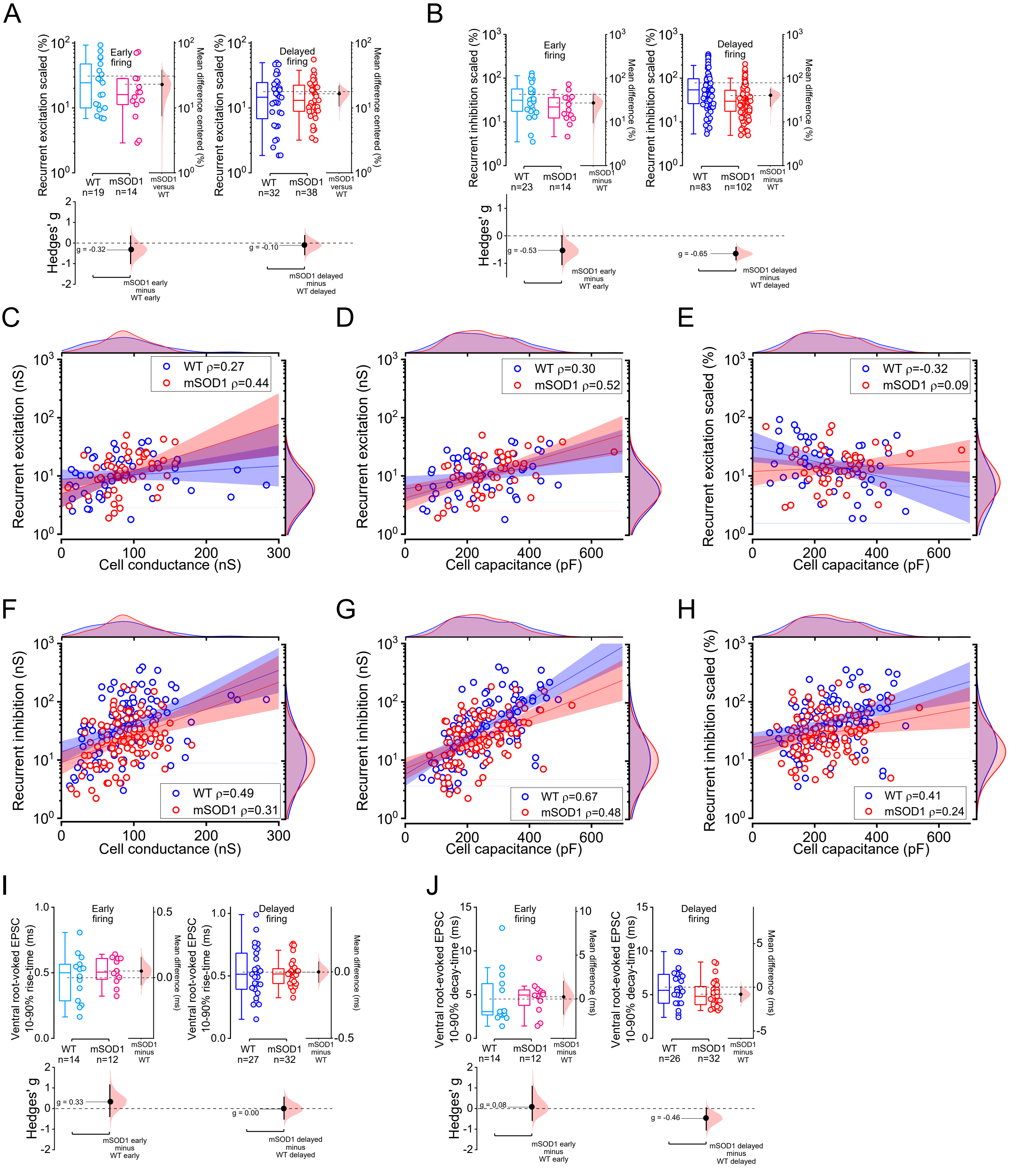

**Figure S3 – Scaled recurrent synaptic conductances, correlation between synaptic conductances of recurrent circuits and cell capacitance or conductance and kinetics of ventral-root evoked currents.** Group data for scaled conductances for **(A)** recurrent excitation and **(B)** recurrent inhibition. Scatter diagrams depicting the correlation between recurrent excitation and motoneuron **(C)** conductance and **(D)** capacitance, and **(E)** recurrent excitation scaled and cell capacitance. Correlation plots between synaptic conductance for recurrent inhibition and **(F)** cell conductance and **(G)** capacitance, and **(H)** recurrent inhibition scaled versus cell capacitance. Estimation plots for **(I)** rise and **(J)** decay times of ventral-root evoked excitatory EPSCs. For the correlation plots, kernel smooth (for linear scale) or lognormal (for log10 axis) distributions are shown next to respective axes, Spearman's rank-order correlation coefficient (ρ) is reported on top of the plot and linear fit (thick line) and 95% confidence interval (shaded area) are shown for visual guidance purposes only. Estimation plots with all individual values and respective box-plots shown along with respective bootstrapped mean difference and bootstrapped Hedges' g. See also Table S3.

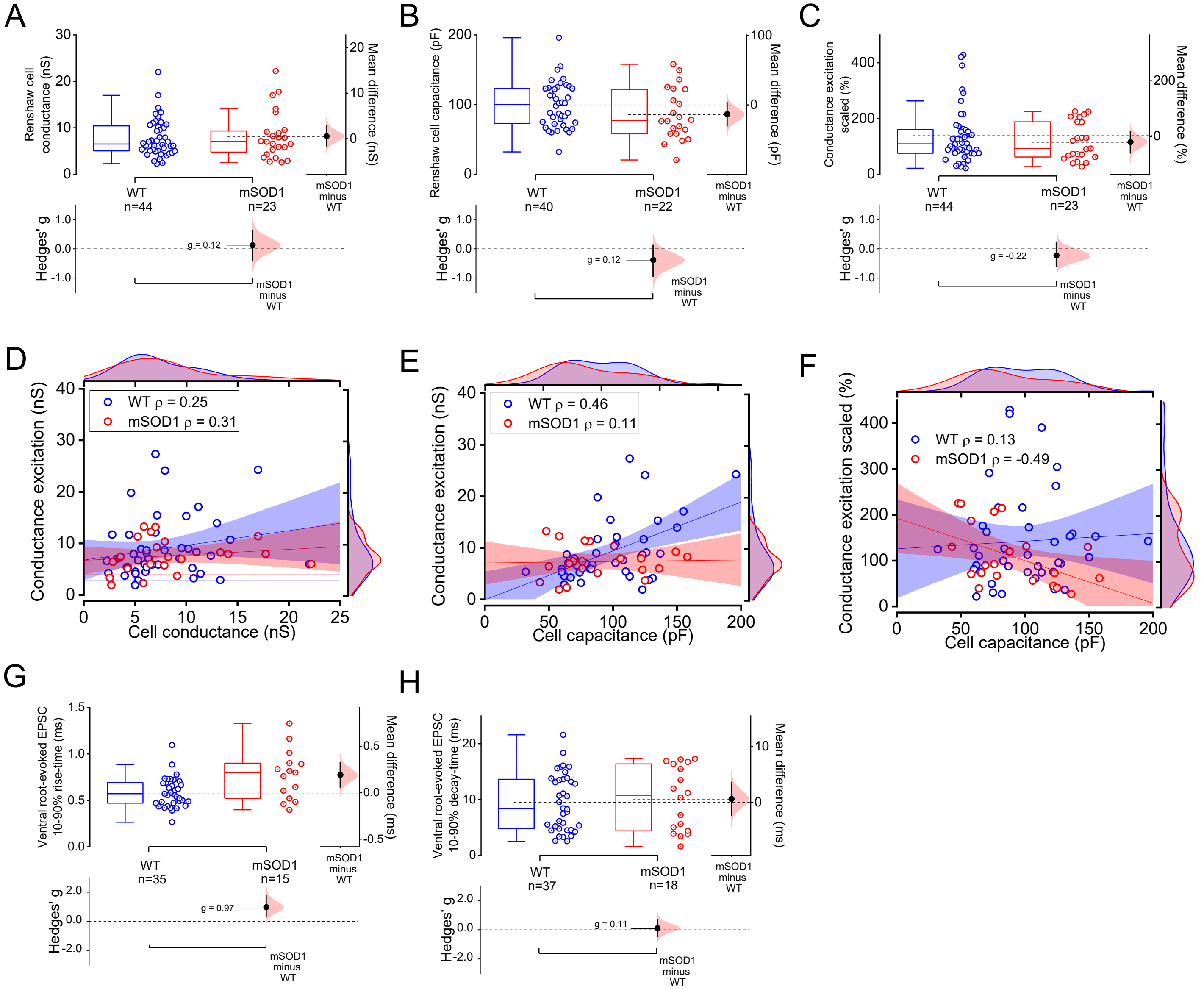

**Figure S4 - Intrinsic properties of Renshaw cells, scaled synaptic conductance and correlation between cell properties and synaptic conductance and kinetics of ventral root-evoked currents.** Estimation plots showing data obtained on Renshaw cell **(A)** resting conductance and **(B)** capacitance and **(C)** scaled synaptic conductance. Relationship between excitation received by Renshaw cells and **(D)** cell conductance and **(E)** capacitance and **(F)** excitation scaled against capacitance. Group data for the **(G)** rise and **(H)** decay time of the ventral root-evoked EPSCs measured from Renshaw cells from WT and mSOD1 mice. For the correlation plots, Kernel smooth distributions are shown next to respective axes, Spearman's rank-order correlation coefficient (ρ) is reported on top of the plot and linear fit (thick line) and 95% confidence interval (shaded area) are shown for visual guidance purposes only. Estimation plots with all individual values and respective box-plots shown along with respective bootstrapped mean difference and bootstrapped Hedges' g. See also Table S4.

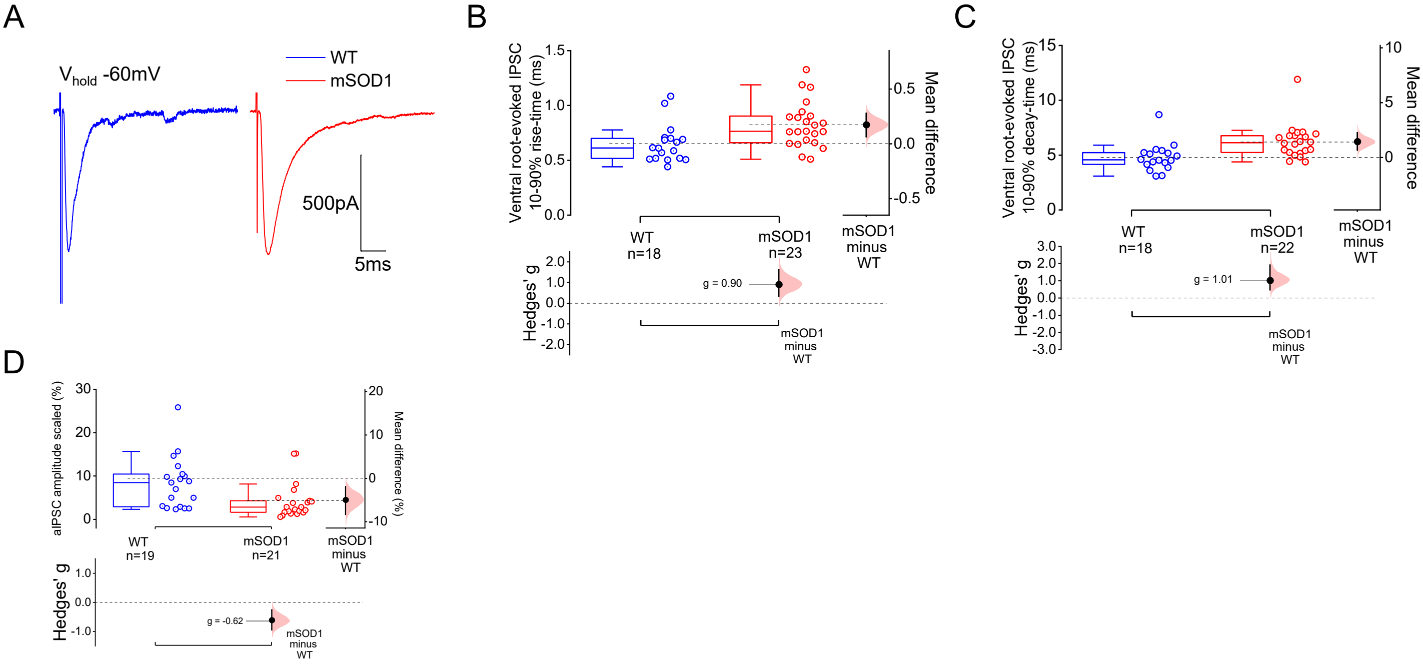

**Figure S5 –Kinetics of ventral-root evoked glycinergic currents and scaled conductance from asynchronous IPSCs (aIPSCs) . (A)** Example of individual IPSCs obtained from WT and mSOD1 mouse motoneurons before replacing $Ca^{2+}$ with $Sr^{2+}$ in the perfusion solution. Group data for IPSC **(B)** rise and **(C)** decay times. **(D)** scaled conductances from aIPSCs. Estimation plots with all individual values and respective box-plots shown along with respective bootstrapped mean difference and bootstrapped Hedges' g. Hierarchical resampling used for figure (D) with each dot representing the mean per animal. See also Table S5.

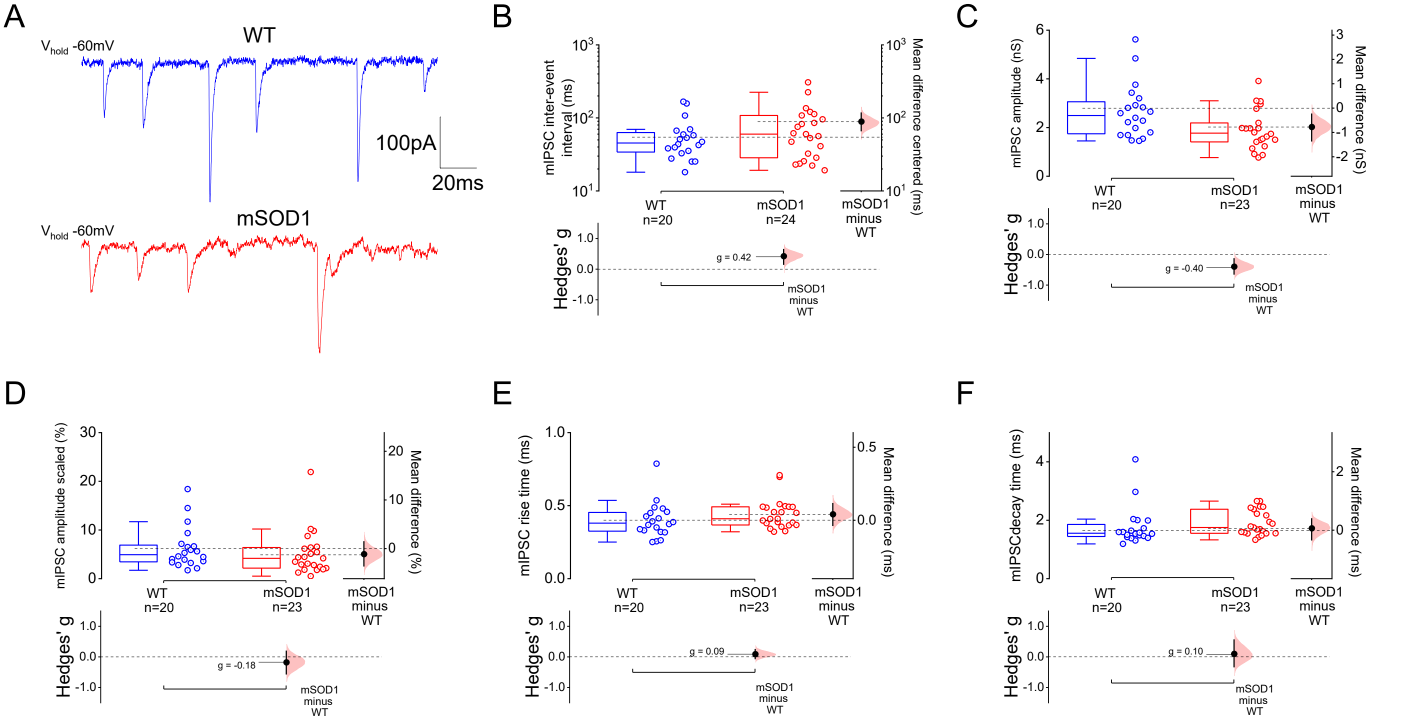

**Figure S6 – Glycinergic mIPSCs are mildly affected in motoneurons from early juvenile mSOD1 mice. (A)** examples of mIPSCs recorded from motoneurons from WT and mSOD1 mice. Estimation plots with data for mIPSC **(B)** inter-event interval, amplitude **(C)** absolute conductance and **(D)** scaled conductance, **(E)** rise and **(F)** decay times. Each dot represents the mean per animal, and mean difference and Hedges' *g* effect sizes were computed using hierarchical resampling. See also Table S5.

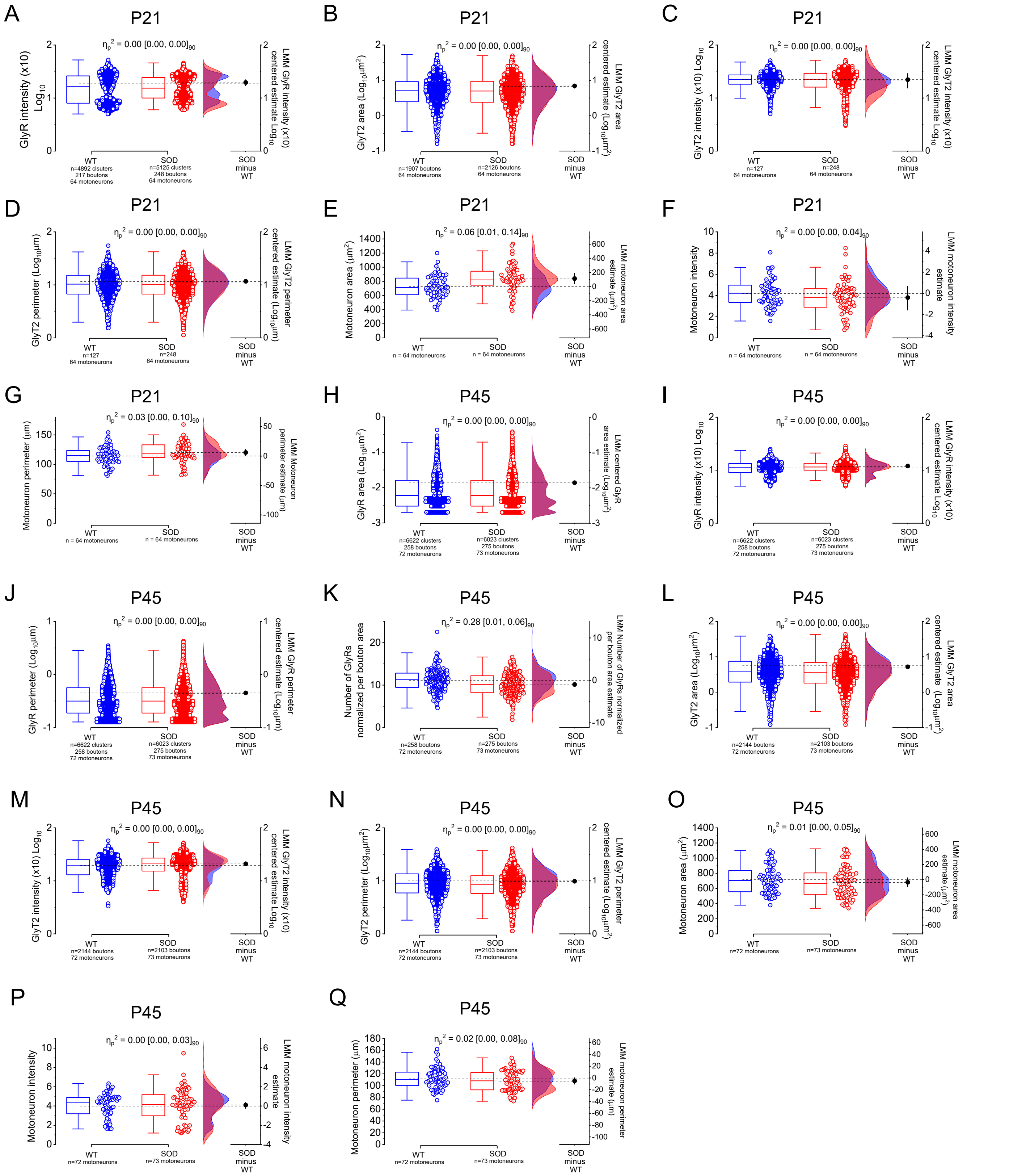

**Figure S7 – Data from P21 and P45 mice immunohistochemistry.** Group data from P21 mice for synaptic markers such as **(A)** GlyR intensity, GlyT2 **(B)** area, **(C)** intensity and **(D)** perimeter and motoneuron **(E)** area, **(F)** intensity and **(G)** perimeter. Group data obtained from P45 mice for GlyR **(H)** area, **(I)** number per bouton, **(J)** perimeter and **(K)** density, presynaptic GlyT2 **(L)** area, **(M)** number per bouton and **(N)** perimeter and motoneuron **(O)** area, **(P)** intensity and **(Q)** perimeter. Box-plots with all individual values and Kernel smooth distribution shown along with respective Linear Mixed Model estimates. See also Tables S6-9.

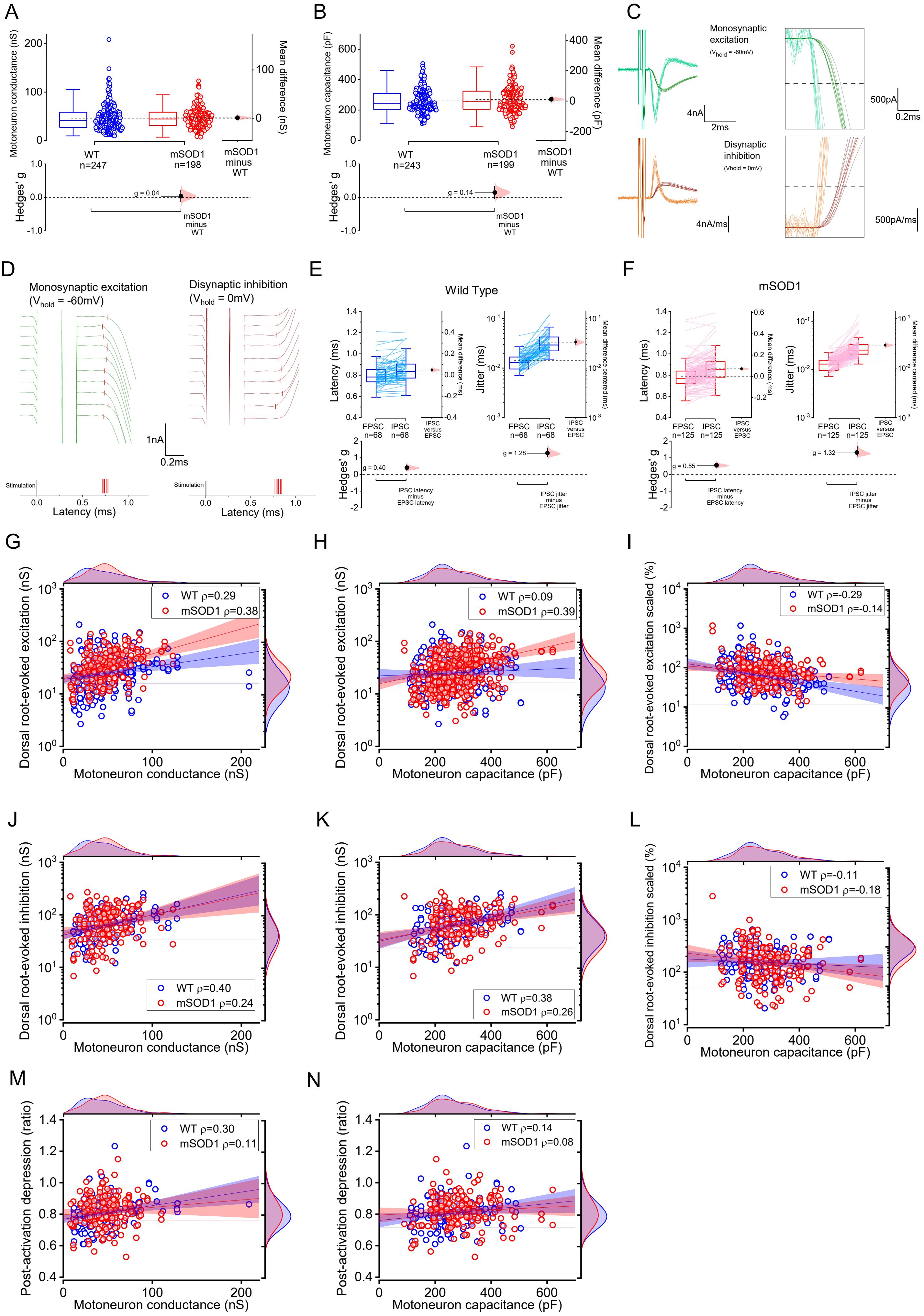
A
Motoneuron conductance (nS)
Mean difference (nS)
WT
n=247
mSOD1
n=198
mSOD1
minus
WT
Hedges' g
g = 0.04
B
Motoneuron capacitance (pF)
Mean difference (pF)
WT
n=243
mSOD1
n=199
g = 0.14
C
Monosynaptic
excitation
($V_{hold}$ = -60mV)
4nA
2ms
500pA
0.2ms
Disynaptic
inhibition
(Vhold = 0mV)
4nA/ms
500pA/ms
D
Monosynaptic excitation
($V_{hold}$ = -60mV)
Disynaptic inhibition
($V_{hold}$ = 0mV)
1nA
0.2ms
Stimulation
Latency (ms)
E
Wild Type
Latency (ms)
Mean difference (ms)
Jitter (ms)
Mean difference centered (ms)
EPSC
n=68
IPSC
n=68
IPSC
versus
EPSC
Hedges' g
g = 0.40
g = 1.28
IPSC latency
minus
EPSC latency
IPSC jitter
minus
EPSC jitter
F
mSOD1
EPSC
n=125
IPSC
n=125
g = 0.55
g = 1.32
G
Dorsal root-evoked excitation (nS)
WT ρ=0.29
mSOD1 ρ=0.38
Motoneuron conductance (nS)
H
WT ρ=0.09
mSOD1 ρ=0.39
Motoneuron capacitance (pF)
I
Dorsal root-evoked excitation scaled (%)
WT ρ=-0.29
mSOD1 ρ=-0.14
J
Dorsal root-evoked inhibition (nS)
WT ρ=0.40
mSOD1 ρ=0.24
K
WT ρ=0.38
mSOD1 ρ=0.26
L
Dorsal root-evoked inhibition scaled (%)
WT ρ=-0.11
mSOD1 ρ=-0.18
M
Post-activation depression (ratio)
WT ρ=0.30
mSOD1 ρ=0.11
N
WT ρ=0.14
mSOD1 ρ=0.08

**Figure S8 – Intrinsic properties of motoneurons obtained from ventral-horn ablated in vitro spinal cords, latency and jitter of dorsal root responses and correlations between motoneuron properties and sensory-evoked synaptic conductances and post-activation depression.** Motoneuron **(A)** conductance and **(B)** capacitance obtained from recordings performed from ventral horn-ablated *in vitro* spinal cords. **(C)** Examples of dorsal-root evoked EPSCs (dark green) and IPSCs (brown) with respective derivative for each trace (lighted colours), with boxes representing zoomed in sections illustrating the 5x standard deviation of the baseline noise from the derivative (dashed line) used as proxy for the onset of the synaptic current. **(D)** EPSCs and IPSCs from (C) vertically aligned, with synaptic current onset demarked by red line and latency rug at the bottom of each series of traces. Jitter and latency obtained from Wild Type **(E)** and mSOD1 **(F)** dorsal root evoked responses. Correlation plots for dorsal root-evoked excitation and **(D)** cell conductance or **(E)** capacitance, **(F)** excitation scaled and capacitance, disynaptic inhibition and **(G)** motoneuron conductance or **(H)** capacitance, inhibition scaled against capacitance **(I)** and post-activation depression ratio versus **(J)** cell conductance or **(K)** capacitance. For the correlation plots, kernel smooth (for linear scale) or lognormal (for log10 axis) distributions are shown next to respective axes, Spearman's rank-order correlation coefficient (ρ) is reported on top of the plot and linear fit (thick line) and 95% confidence interval (shaded area) are shown for visual guidance purposes only. Estimation plots with all individual values and respective box-plots shown along with respective bootstrapped mean difference and bootstrapped Hedges' g. See also Tables S11-S18.

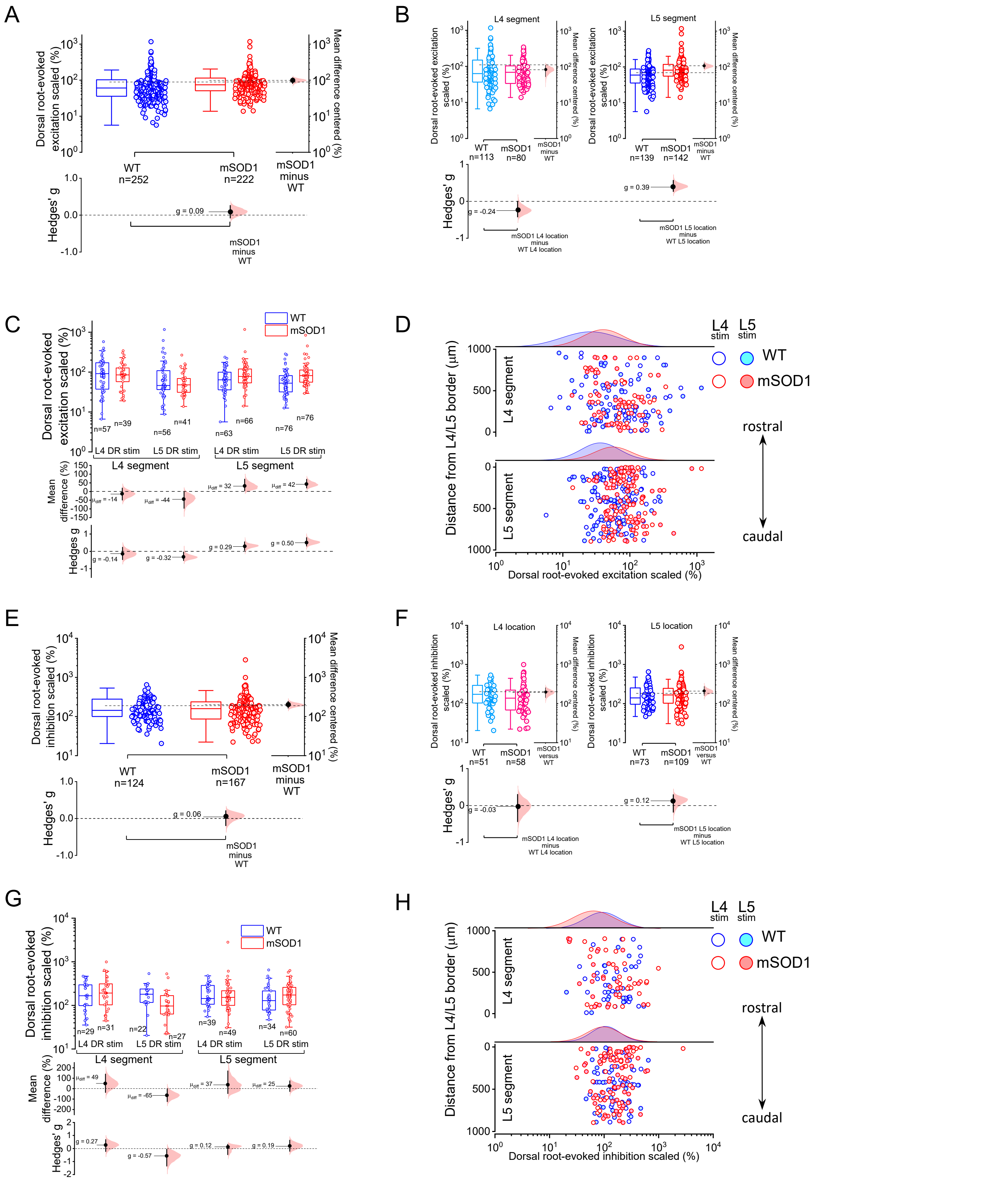

**Figure S9 – Dorsal root-evoked monosynaptic (Ia) excitation and disynaptic (Ia/Ib) inhibition scaled conductances.** **(A)** Data for monosynaptic Ia excitation scaled to resting cell conductance, **(B)** organized according to segment position and **(C)** according to stimulated root and location. **(D)** the values split by cell location (left) and with scaled conductances plotted against distance from L4/L5 border (right). Data for absolute and scaled conductance of disynaptic Ia/Ib inhibition organized according to segment and **(E and F)** grouped based on stimulated root and location **(G and H)**, and values organized by cell location (left) and with scaled inhibitory conductances plotted against distance from L4/L5 border (right). Estimation plots with all individual values and respective box-plots shown along with respective bootstrapped mean difference and bootstrapped Hedges' g. See also Tables S11-S17.

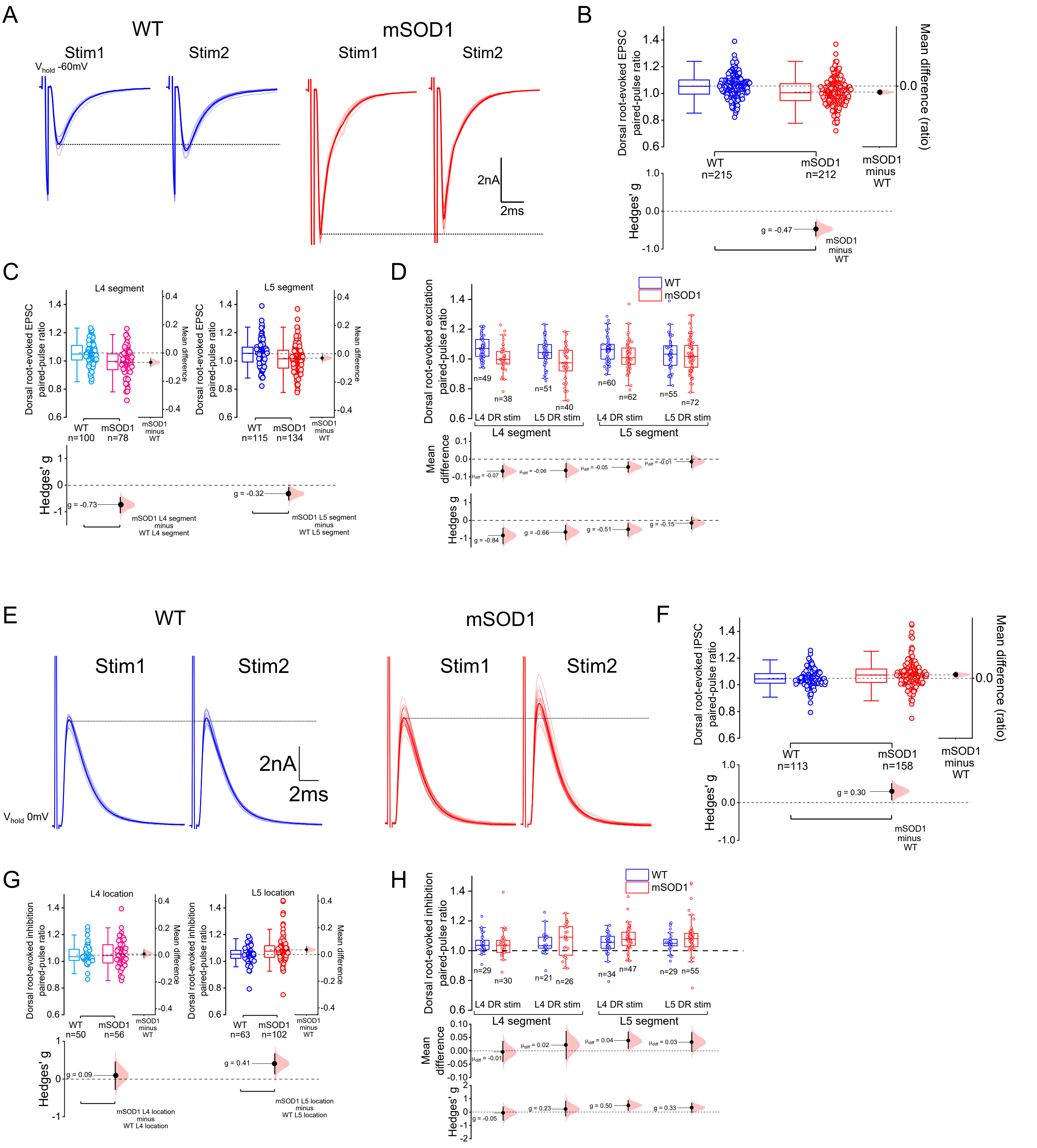

**Figure S10 – Dorsal root-evoked monosynaptic (Ia) excitation and disynaptic (Ia/Ib) inhibition paired-pulse ratios. (A)** Examples of dorsal-root evoked paired pulse responses (33 Hz) for WT and mSOD1 mice, with group data for **(B)** all observations, **(C)** recordings organized per location and **(D)** ratios grouped per segment and stimulating root. **(E)** Example traces of dorsal-root evoked disynaptic inhibitory paired pulse responses (33 Hz) for control and mutant mice, with group data for **(F)** all observations, **(G)** recordings organized per anatomical position and **(H)** ratios grouped per location and stimulating root. Estimation plots with all individual values and respective box-plots shown along with respective bootstrapped mean difference and bootstrapped Hedges' g. See also Tables S11-S17.

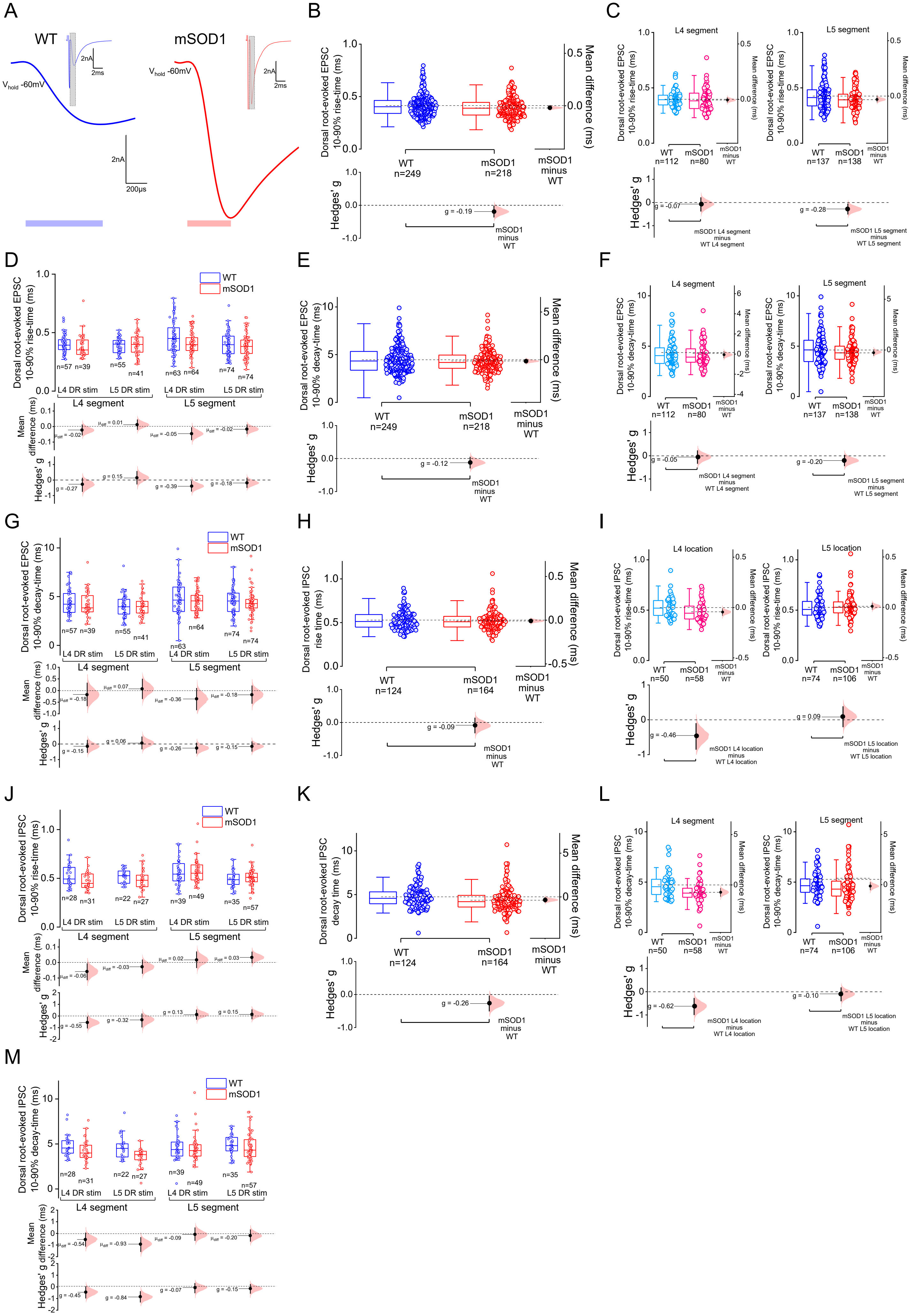

A
WT
mSOD1
V hold -60mV
2nA
2ms
200μs
B
Dorsal root-evoked EPSC 10-90% rise-time (ms)
Mean difference (ms)
WT n=249
mSOD1 n=218
mSOD1 minus WT
Hedges' g
g = -0.19
C
L4 segment
L5 segment
Dorsal root-evoked EPSC 10-90% rise-time (ms)
WT n=112
mSOD1 n=80
WT n=137
mSOD1 n=138
g = -0.07
g = -0.28
mSOD1 L4 segment minus WT L4 segment
mSOD1 L5 segment minus WT L5 segment
D
Dorsal root-evoked EPSC 10-90% rise-time (ms)
WT
mSOD1
n=57 n=39
n=55 n=41
n=63 n=64
n=74 n=74
L4 DR stim
L5 DR stim
L4 segment
L5 segment
Mean difference (ms)
μdiff = -0.02
μdiff = 0.01
μdiff = -0.05
μdiff = -0.02
Hedges' g
g = -0.27
g = 0.15
g = -0.39
g = -0.18
E
Dorsal root-evoked EPSC 10-90% decay-time (ms)
WT n=249
mSOD1 n=218
mSOD1 minus WT
g = -0.12
F
L4 segment
L5 segment
Dorsal root-evoked EPSC 10-90% decay-time (ms)
WT n=112
mSOD1 n=80
WT n=137
mSOD1 n=138
g = -0.05
g = -0.20
mSOD1 L4 segment minus WT L4 segment
mSOD1 L5 segment minus WT L5 segment
G
Dorsal root-evoked EPSC 10-90% decay-time (ms)
n=57 n=39
n=55 n=41
n=63 n=64
n=74 n=74
μdiff = -0.18
μdiff = 0.07
μdiff = -0.36
μdiff = -0.18
g = -0.15
g = 0.06
g = -0.26
g = -0.15
H
Dorsal root-evoked IPSC rise time (ms)
WT n=124
mSOD1 n=164
mSOD1 minus WT
g = -0.09
I
L4 location
L5 location
Dorsal root-evoked IPSC 10-90% rise-time (ms)
WT n=50
mSOD1 n=58
WT n=74
mSOD1 n=106
g = -0.46
g = 0.09
mSOD1 L4 location minus WT L4 location
mSOD1 L5 location minus WT L5 location
J
Dorsal root-evoked IPSC 10-90% rise-time (ms)
n=28 n=31
n=22 n=27
n=39 n=49
n=35 n=57
μdiff = -0.06
μdiff = -0.03
μdiff = 0.02
μdiff = 0.03
g = -0.55
g = -0.32
g = 0.13
g = 0.15
K
Dorsal root-evoked IPSC decay time (ms)
WT n=124
mSOD1 n=164
mSOD1 minus WT
g = -0.26
L
L4 segment
L5 segment
Dorsal root-evoked IPSC 10-90% decay-time (ms)
WT n=50
mSOD1 n=58
WT n=74
mSOD1 n=106
g = -0.62
g = -0.10
mSOD1 L4 location minus WT L4 location
mSOD1 L5 location minus WT L5 location
M
Dorsal root-evoked IPSC 10-90% decay-time (ms)
n=28 n=31
n=22 n=27
n=39 n=49
n=35 n=57
μdiff = -0.54
μdiff = -0.93
μdiff = -0.09
μdiff = -0.20
g = -0.45
g = -0.84
g = -0.07
g = -0.15

**Figure S11– Dorsal root-evoked monosynaptic (Ia) EPSC and disynaptic (Ia/Ib) IPSC rise and decay times. (A)** Close up of the initial phase of the example dorsal root-evoked EPSCs from WT and mSOD1 with the duration of the rise phase represented at the bottom. Estimation plots showing data for all root responses **(B and E)**, recordings grouped by segment **(C and F)** and results shown by location and stimulating root **(D and G)** for rise and decay phases of the dorsal-root evoked Ia monosynaptic EPSC. Group data for all disynaptic root responses **(H and K)**, recordings grouped by segment **(I and L)** and results shown by location and stimulating root **(J and M)** for rise and decay phases of the dorsal-root evoked Ia/Ib disynaptic EPSC. Estimation plots with all individual values and respective box-plots shown along with respective bootstrapped mean difference and bootstrapped Hedges' g. See also Tables S11-S17.

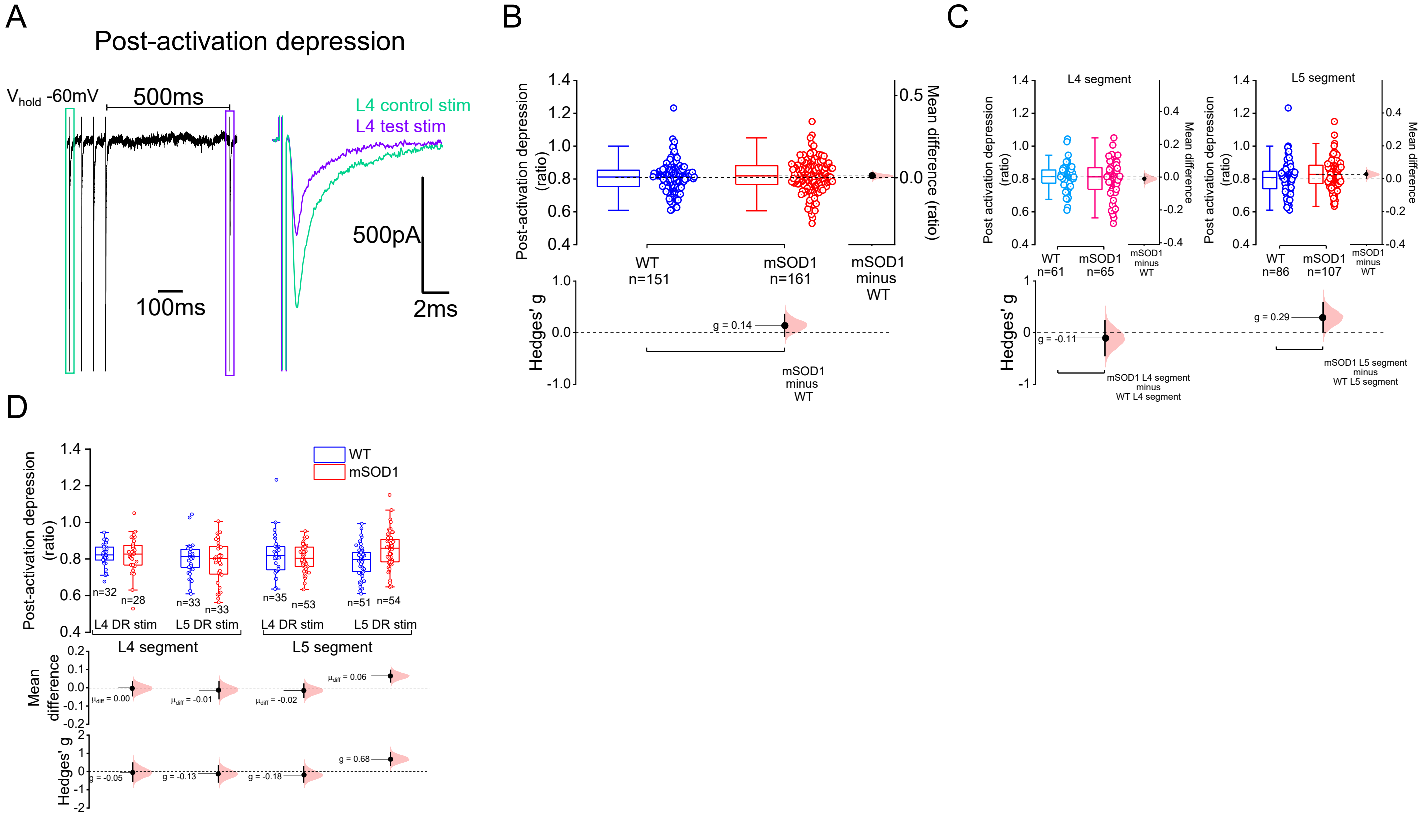

**Figure S12 – Post-activation depression responses organized according to segment location and stimulating root. (I)** Example of the conditioning protocol used to test post-activation depression *in vitro*, with the amplitude of a first EPSC from a train of 4 responses at 20 Hz (L4 control stim) is compared to that of a fifth EPSC triggered 500 ms after the 4$^{th}$ repetitive stimuli (L4 test stim). **(J)** Group data for post-activation depression.**(A)** Data for post-activation depression grouped according to segment position and **(B)** to stimulated root and location**.** Estimation plots with all individual values and respective box-plots shown along with respective bootstrapped mean difference and bootstrapped Hedges' g. See also Tables S11-S17.

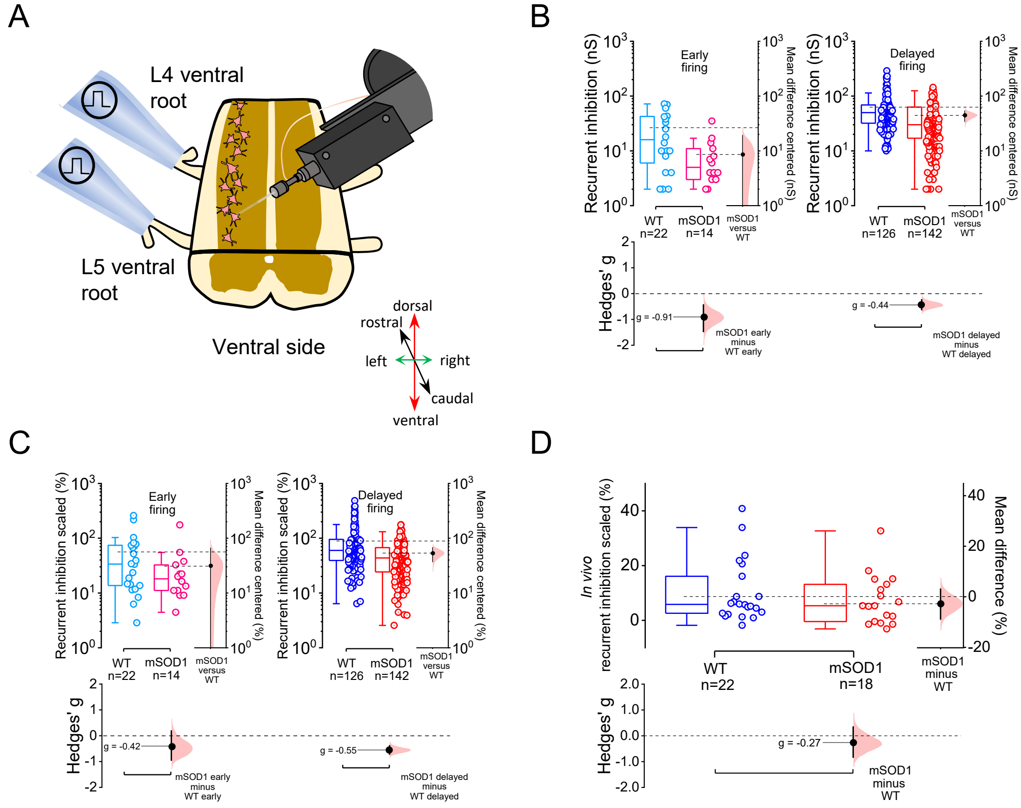

**Figure S13 – *In vitro* recurrent inhibition from distal muscle-innervating lumbar motoneurons and scaled recurrent inhibition conductances from *in vivo* recordings. (A)** Representation of the dorsal horn-ablated *in vitro* spinal cord preparation used to estimate recurrent inhibition from favorably targeted dorsolateral motoneurons (each *n* corresponds to a root response), with respective data obtained on **(B)** absolute and **(C)** scaled recurrent inhibition. **(D)** Group data for scaled recurrent inhibition estimated from *in vivo* motoneuron recordings. Estimation plots with all individual values and respective box-plots shown along with respective bootstrapped mean difference and bootstrapped Hedges' *g*. See also Table S22.

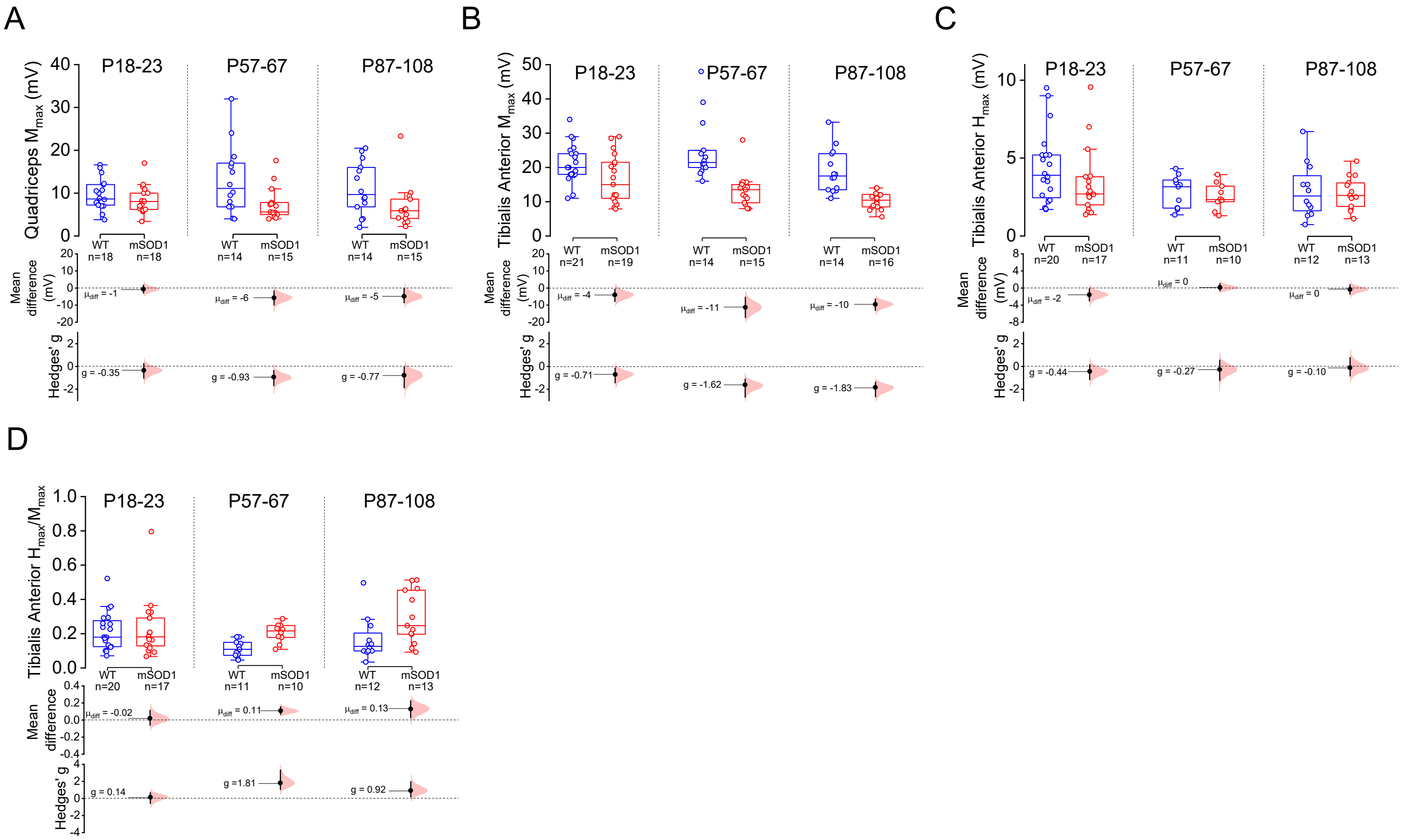

**Figure S14 – $M_{max}$ and $H_{max}$ values obtained from *in vivo* EMG experiments.** Data obtained on **(A)** quadriceps $M_{max}$ and Tibialis Anterior **(B)** $M_{max}$ , **(C)** $H_{max}$ and **(D)** $H_{max}/M_{max}$ ratio obtained from different age ranges (each *n* is an animal). Estimation plots with all individual values and respective box-plots shown along with respective bootstrapped mean difference and bootstrapped Hedges' *g*. See also Table S20.

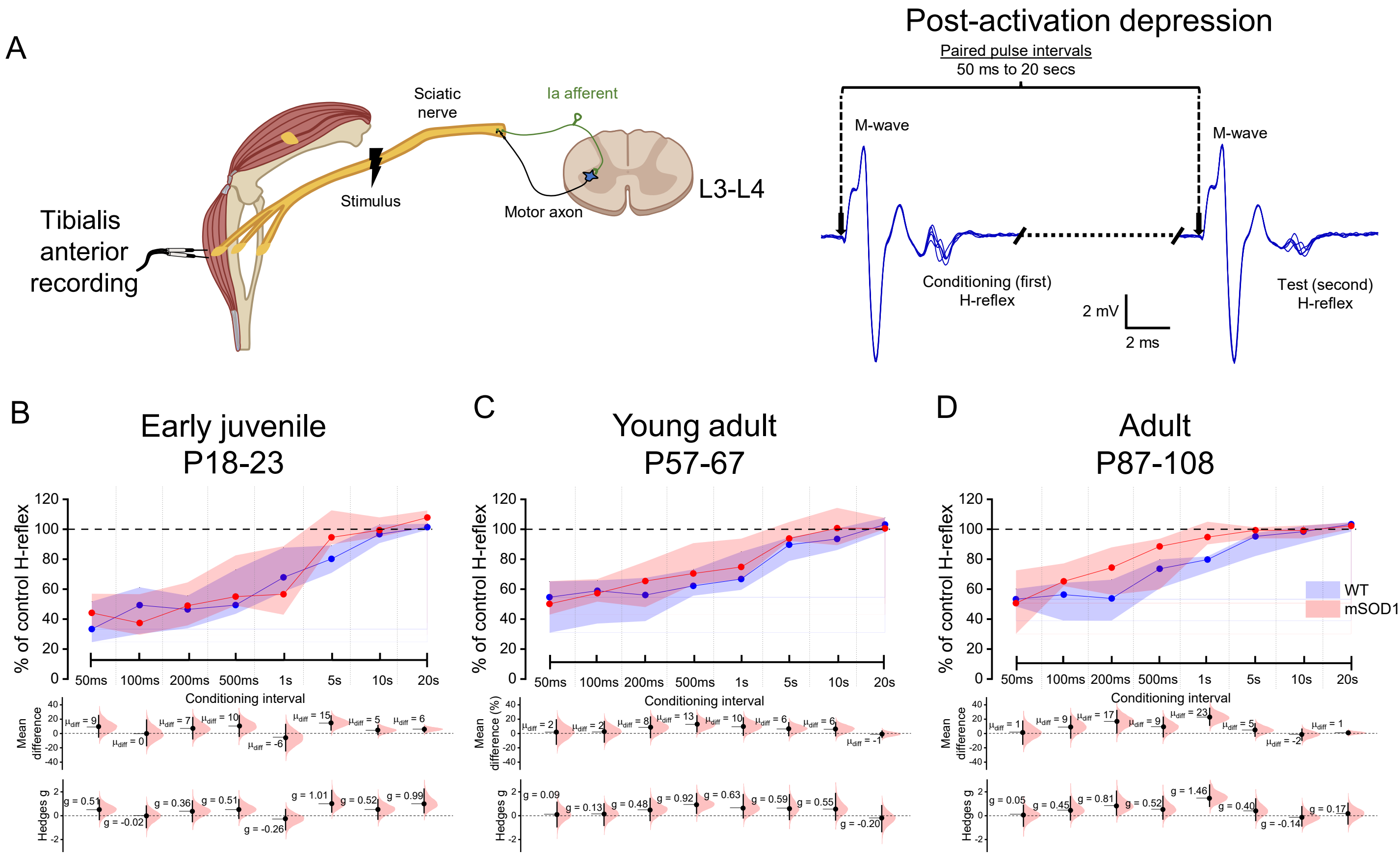

**Figure S15 – Post-activation depression from *in vivo* EMG experiments.** Schematic and example traces illustrating the EMG recordings used to obtain motor and H-reflex responses from tibialis anterior muscles, and the conditioning protocols used to estimate **(A)** post-activation depression. Data obtained for **(B-D)** post-activation depression (each *n* is an animal). Estimation plots with box-plots shown as median (dot) and interquartile range (shaded area) for post-activation depression along with respective bootstrapped mean difference and bootstrapped Hedges' ***g*****.** See also Table S21.

**Table S1 –** Mean and standard deviation, intraclass correlation coefficient (ICC), bootstrapped mean difference and Hedges' *g* for intrinsic properties and firing output from early and delayed firing motoneurons obtained from oblique slices

| Early firing motoneurons | | | | | | |
|---|---|---|---|---|---|---|
| Parameter | Mean±standard deviation | | ICC | | Bootstrapped Mean difference [95% CI] | Bootstrapped Hedges' *g* [95% CI] |
| | WT | mSOD1 | WT | mSOD1 | | |
| Conductance (ns) | 51±43 n=36 19 mice | 56±24 n=28 15 mice | 0.06 | **0.65** | 5 [-12, 20] | 0.14 [-0.27, 0.84] |
| Capacitance (pF) | 151±66 n=36 19 mice | 168±63 n=28 15 mice | -0.32 | -0.33 | 17 [-14, 49] | 0.26 [-0.23, 0.82] |
| Membrane potential (mv) | -62±4 n=30 18 mice | -62±3 n=27 15 mice | **0.60** | -0.27 | 0 [-2, 2] | -0.02 [-0.56, 0.47] |
| Rheobase (nA) | 0.95±0.60 n=30 17 mice | 1.36±0.76 n=27 15 mice | -0.14 | 0.19 | **0.41 [0.06, 0.77]** | **0.60 [0.08, 1.17]** |
| Maximum sustained firing (Hz) | 110±46 n=11 5 mice | 95±24 n=18 10 mice | 0.00 | -0.18 | -15 [-45, 11] | -0.43 [-1.20, 0.48] |
| 1-2nd spike interval (Hz) | 411±181 n=11 5 mice | 436±122 n=16 9 mice | **0.67** | 0.16 | 26 [-92, 144] | 0.17 [-0.69, 1.08] |
| Depolarizing block (nA) | 4.96±3.10 n=10 5 mice | 6.32±1.69 n=18 10 mice | -0.08 | -0.17 | 1.36 [-0.63, 3.36] | 0.60 [-0.31, 1.60] |
| Delayed firing motoneurons | | | | | | |
| Parameter | Mean±standard deviation | | ICC | | Bootstrapped Mean difference [95% CI] | Bootstrapped Hedges' *g* [95% CI] |
| | WT | mSOD1 | WT | mSOD1 | | |
| Conductance (ns) | 104±44 n=98 31 mice | 94±30 n=139 29 mice | 0.33 | 0.11 | -10 [-20, 0.1] | -0.27 [-0.52, 0.01] |
| Capacitance (pF) | 286±93 n=98 31 mice | 266±94 n=138 29 mice | 0.05 | 0.40 | -20 [-42, 5] | -0.21 [-0.47, 0.05] |
| Membrane potential (mv) | -62±4 n=91 29 mice | -63±3 n=129 28 mice | 0.00 | -0.04 | -0.5 [-1.4, 0.3] | -0.18 [-0.46, 0.10] |
| Rheobase (nA) | 2.69±1.54 n=86 29 mice | 2.49±1.21 n=125 28 mice | 0.29 | 0.07 | -0.20 [-0.59, 0.19] | -0.15 [-0.43, 0.14] |
| Maximum sustained firing (Hz) | 89±22 n=27 9 mice | 97±20 n=64 12 mice | 0.06 | 0.21 | 7 [-3, 17] | 0.34 [-0.12, 0.83] |
| 1-2nd spike interval (Hz) | 437±151 n=24 9 mice | 406±156 n=57 11 mice | 0.21 | 0.19 | -31 [-102, 40] | -0.20 [-0.66, 0.28] |
| Depolarizing block (nA) | 8.67±1.86 n=24 9 mice | 8.79±1.44 n=62 12 mice | 0.03 | 0.07 | 0.12 [-0.66, 0.96] | 0.08 [-0.43, 0.65] |

CI - confidence interval; for mixed model results when ICC>0.50 see table S24

**Table S2 -** Mean and standard deviation, intraclass correlation coefficient (ICC), bootstrapped mean difference and Hedges' *g* for action potential properties extracted from early and delayed firing motoneuron recordings from oblique slices

| Early firing motoneurons | | | | | | |
|---|---|---|---|---|---|---|
| Parameter | Mean±standard deviation | | ICC | | Bootstrapped Mean difference [95% CI] | Bootstrapped Hedges' *g* [95% CI] |
| | WT | mSOD1 | WT | mSOD1 | | |
| Threshold (mV) | -32±7 n=27 15 mice | -32±7 n=23 14 mice | 0.12 | 0.21 | 0 [-1, 1] | 0.04 [-0.56, 0.59] |
| Action potential amplitude (mV) | 65±11 n=27 15 mice | 68±8 n=23 14 mice | 0.07 | 0.18 | 3 [-2, 9] | 0.33 [-0.20, 1.01] |
| Action potential half-width (ms) | 0.23±0.06 n=27 15 mice | 0.26±0.10 n=23 14 mice | **0.58** | **0.92** | 0.03 [-0.02, 0.07] | 0.31 [-0.28, 0.78] |
| Action potential rise time (ms) | 0.18±0.04 n=27 15 mice | 0.19±0.04 n=23 14 mice | 0.37 | **0.51** | 0.01 [-0.02, 0.03] | 0.12 [-0.47, 0.66] |
| Action potential max depolarization rate (mV/ms) | 461±97 n=26 15 mice | 470±103 n=23 14 mice | - 0.05 | 0.36 | 9 [-46, 63] | 0.09 [-0.49, 0.67] |
| Action potential decay time (ms) | 0.16±0.06 n=27 15 mice | 0.19±0.10 n=23 14 mice | **0.54** | **0.94** | 0.03 [-0.01, 0.08] | 0.38 [-0.17, 0.85] |
| Action potential max repolarization rate (mV/ms) | -406±136 n=26 15 mice | -358±95 n=23 14 mice | 0.33 | 0.46 | 48 [-13, 113] | 0.40 [-0.14, 0.92] |
| Fast AHP amplitude (mV) | -21±5 n=27 15 mice | -19±4 n=23 14 mice | 0.49 | 0.20 | **3 [0.2, 5]** | **0.59 [0.06, 1.22]** |
| ADP amplitude (mV) | 3.3±2.0 n=27 15 mice | 4.2±1.8 n=22 14 mice | 0.43 | -0.34 | 0.9 [-0.1, 1.9] | 0.48 [-0.06, 1.13] |
| mAHP amplitude (mV) | -21±5 n=26 15 mice | -18±5 n=23 14 mice | 0.15 | 0.44 | 3 [-0.03, 5] | 0.56 [-0.01, 1.31] |
| mAHP half-width (ms) | 44±41 n=24 15 mice | 35±20 n=18 14 mice | 0.33 | -0.25 | -9 [-30, 7] | -0.27 [-0.70, 0.39] |
| Delayed firing motoneurons | | | | | | |
| Parameter | Mean±standard deviation | | ICC | | Bootstrapped Mean difference [95% CI] | Bootstrapped Hedges' *g* [95% CI] |
| | WT | mSOD1 | WT | mSOD1 | | |
| Threshold (mV) | -29±5 n=70 28 mice | -28±6 n=115 25 mice | 0.03 | 0.05 | 1 [-1, 2] | 0.14 [-0.14, 0.43] |
| Action potential amplitude (mV) | 63±6 n=73 28 mice | 63±8 n=116 25 mice | 0.24 | 0.12 | 0 [-2, 2] | -0.03 [-0.31, 0.25] |
| Action potential half-width (ms) | 0.22±0.03 n=71 | 0.21±0.03 n=107 | 0.12 | 0.31 | 0.00 [-0.01, 0.01] | -0.13 [-0.44, 0.18] |

| | | | | | | |
|---|---|---|---|---|---|---|
| | 28 mice | 23 mice | | | | |
| Action potential rise time (ms) | 0.17±0.03<br>n=70<br>28 mice | 0.17±0.02<br>n=107<br>23 mice | 0.03 | 0.22 | 0.00<br>[-0.01, 0.01] | -0.08<br>[-0.38, 0.22] |
| Action potential max depolarization rate (mV/ms) | 459±66<br>n=70<br>28 mice | 457±75<br>n=107<br>23 mice | -<br>0.13 | 0.27 | -2<br>[-23, 19] | -0.03<br>[-0.32, 0.27] |
| Action potential decay time (ms) | 0.15±0.03<br>n=70<br>28 mice | 0.15±0.03<br>n=107<br>23 mice | 0.37 | 0.40 | 0.00<br>[-0.01, 0.004] | -0.14<br>[-0.44, 0.17] |
| Action potential max repolarization rate (mV/ms) | -386±60<br>n=70<br>28 mice | -384±55<br>n=107<br>23 mice | 0.01 | 0.26 | 2<br>[-16, 19] | 0.03<br>[-0.29, 0.33] |
| Fast AHP amplitude (mV) | -24±3<br>n=72<br>28 mice | -23±4<br>n=116<br>25 mice | -<br>0.08 | 0.13 | 1<br>[-0.3, 2] | 0.20<br>[-0.08, 0.48] |
| ADP amplitude (mV) | 4.1±1.9<br>n=72<br>28 mice | 4.2±1.9<br>n=114<br>25 mice | 0.12 | 0.41 | 0.1<br>[-0.5, 0.6] | 0.04<br>[-0.26, 0.34] |
| Medium AHP amplitude (mV) | -23±4<br>n=70<br>28 mice | -23±5<br>n=116<br>25 mice | 0.17 | 0.24 | 0<br>[-1, 1] | -0.02<br>[-0.30, 0.26] |
| Medium AHP half-width (ms) | 30±13<br>n=69<br>28 mice | 26±11<br>n=113<br>25 mice | 0.44 | 0.00 | **-4<br>[-8, -1]** | **-0.35<br>[-0.69, -0.05]** |

CI - confidence interval; ADP - afterdepolarization; AHP - afterhyperpolarization; for mixed model results when ICC>0.50 see table S24

**Table S3 –** Mean and standard deviation, intraclass correlation coefficient (ICC), bootstrapped mean difference and Hedges' *g* for recurrent excitatory and inhibitory conductances measured from early and delayed firing motoneurons from oblique slices

| Early firing motoneurons | | | | | | |
|---|---|---|---|---|---|---|
| Parameter | Mean±standard deviation | | ICC | | Bootstrapped Mean difference [95% CI] | Bootstrapped Hedges' *g* [95% CI] |
| | WT | mSOD1 | WT | mSOD1 | | |
| Recurrent excitation (ns) | 10±7 n=19 10 mice | 9±6 n=14 7 mice | **0.71** | -0.63 | 2 [-2, 6] | -0.06 [-0.69, 0.65] |
| Recurrent excitation scaled (%) | 31±26 n=19 10 mice | 23±22 n=14 7 mice | 0.34 | **0.53** | -3 [-18, 13] | -0.32 [-1.02, 0.37] |
| Ventral root-evoked EPSC rise-time | 0.46±0.18 n=14 10 mice | 0.51±0.10 n=12 7 mice | 0.17 | **0.67** | 0.05 [-0.06, 0.16] | 0.33 [-0.41, 1.18] |
| Ventral root-evoked EPSC decay-time | 4.52±3.10 n=14 10 mice | 4.74±2.03 n=12 7 mice | **0.91** | **0.88** | 0.22 [-1.81, 2.05] | 0.08 [-0.60, 1.11] |
| Recurrent inhibition (nS) | 18±17 n=23 15 mice | 15±12 n=14 9 mice | -1.18 | -0.54 | -5 [-15, 5] | -0.21 [-0.78, 0.46] |
| Recurrent inhibition scaled (%) | 44±36 n=23 15 mice | 27±19 n=14 9 mice | -0.87 | -0.62 | -16 [-34, 1] | -0.53 [-1.07, 0.02] |
| Delayed firing motoneurons | | | | | | |
| Parameter | Mean±standard deviation | | ICC | | Bootstrapped Mean difference [95% CI] | Bootstrapped Hedges' *g* [95% CI] |
| | WT | mSOD1 | WT | mSOD1 | | |
| Recurrent excitation (ns) | 16±10 n=32 15 mice | 16±11 n=38 13 mice | 0.17 | 0.08 | 1 [-4, 6] | 0.07 [-0.41, 0.55] |
| Recurrent excitation scaled (%) | 18±14 n=32 15 mice | 17±11 n=38 13 mice | 0.34 | 0.04 | -1 [-7, 5] | -0.10 [-0.59, 0.39] |
| Ventral root-evoked EPSC rise-time | 0.52±0.20 n=27 13 mice | 0.52±0.11 n=32 11 mice | **0.54** | 0.48 | 0.00 [-0.08, 0.08] | 0.00 [-0.54, 0.58] |
| Ventral root-evoked EPSC decay-time | 5.84±1.99 n=26 13 mice | 5.04±1.48 n=32 11 mice | **0.52** | 0.29 | -0.80 [-1.71, 0.10] | -0.46 [0.05, -1.07] |
| Recurrent inhibition (nS) | 78±78 n=83 26 mice | 36±32 n=102 23 mice | 0.45 | 0.29 | **-42 [-61, -25]** | **-0.74 [-0.98, -0.50]** |
| Recurrent inhibition scaled (%) | 79±78 n=83 26 mice | 41±36 n=102 23 mice | 0.47 | 0.42 | **-38 [-56, -21]** | **-0.65 [-0.90, -0.40]** |

CI - confidence interval; EPSC - excitatory postsynaptic current; for mixed model results when ICC>0.50 see table S24

**Table S4** – Mean and standard deviation, intraclass correlation coefficient (ICC), bootstrapped mean difference and Hedges' *g* for Renshaw cell intrinsic properties, ventral root excitation and Bayesian Quantal Analysis (BQA)

| Renshaw cells | | | | | | |
|---|---|---|---|---|---|---|
| Parameter | Mean±standard deviation | | ICC | | Bootstrapped Mean difference [95% CI] | Bootstrapped Hedges' *g* [95% CI] |
| | WT | mSOD1 | WT | mSOD1 | | |
| Cell conductance (nS) | 7.8±4.1 n=44 15 mice | 8.3±5.1 n=24 9 mice | 0.06 | 0.09 | 0.5 [-1.7, 3.0] | 0.12 [-0.42, 0.66] |
| Cell capacitance (pF) | 99±33 n=40 15 mice | 86±37 n=22 9 mice | 0.37 | 0.18 | -13 [-31, 5] | -0.39 [-0.96, 0.14] |
| Ventral root excitation (ns) | 9.0±6.0 n=44 15 mice | 7.6±3.3 n=23 9 mice | 0.05 | 0.11 | -1.8 [-4.0, 0.3] | -0.27 [-0.64, 0.16] |
| Ventral root excitation scaled (%) | 137±100 n=44 15 mice | 117±68 n=23 9 mice | 0.07 | 0.36 | -20 [-61, 19] | -0.22 [-0.62, 0.25] |
| Ventral root-evoked EPSC rise-time (ms) | 0.58±0.16 n=35 14 mice | 0.77±0.27 n=15 7 mice | 0.27 | 0.42 | **0.19 [0.05, 0.33]** | **0.97 [0.30, 1.80]** |
| Ventral root-evoked EPSC decay-time (ms) | 9.49±5.15 n=37 14 mice | 10.09±5.74 n=18 7 mice | 0.27 | 0.50 | 0.60 [-2.40, 3.69] | 0.11 [-0.49, 0.73] |
| BQA quantal size (mV) | 0.82±0.55 n=13 8 mice | 0.84±0.65 n=11 5 mice | -0.44 | 0.35 | 0.02 [-0.43, 0.50] | 0.04 [-0.88, 0.87] |
| BQA number of release sites | 31±26 n=13 8 mice | 36±27 n=11 5 mice | 0.04 | -0.72 | 5 [-16, 25] | 0.19 [-0.63, 1.13] |
| BQA probability of release $[Ca^{2+}]$ = 2mM | 0.63±0.29 n=13 8 mice | 0.81±0.20 n=11 5 mice | **0.66** | -0.23 | 0.16 [-0.03, 0.34] | 0.71 [-0.04, 1.66] |
| BQA probability of release $[Ca^{2+}]$ = 1mM | 0.21±0.17 n=13 8 mice | 0.31±0.12 n=11 5 mice | **0.92** | -0.08 | 0.10 [-0.02, 0.20] | 0.66 [-0.10, 1.97] |

CI - confidence interval; EPSC - excitatory postsynaptic current; for mixed model results when ICC>0.50 see table S24

**Table S5 –** Mean and standard deviation, intraclass correlation coefficient (ICC), bootstrapped mean difference and Hedges' *g* for Bayesian Quantal Analysis (BQA), kinetics of ventral-root evoked EPSCs, aIPSCs amplitude and mIPSCs amplitude, frequency and kinetics obtained from recordings performed from motoneurons from oblique slices

| Motoneurons BQA | | | | | | |
|---|---|---|---|---|---|---|
| Parameter | Mean±standard deviation | | ICC | | Bootstrapped Mean difference [95% CI] | Bootstrapped Hedges' *g* [95% CI] |
| | WT | mSOD1 | WT | mSOD1 | | |
| BQA quantal size (mV) | -0.040±0.017 n=23 11 mice | -0.028±0.012 n=22 10 mice | 0.13 | -0.04 | **0.012 [0.004, 0.020]** | **0.81 [0.26, 1.47]** |
| BQA number of release sites | 258±159 n=23 11 mice | 344±275 n=22 10 mice | -0.09 | 0.33 | 86 [-42, 218] | 0.39 [-0.19, 1.02] |
| BQA probability of release [$Ca^{2+}$] = 2mM | 0.15±0.11 n=23 11 mice | 0.19±0.13 n=22 10 mice | -0.04 | 0.13 | 0.03 [-0.04, 0.10] | 0.26 [-0.34, 0.86] |
| BQA probability of release [$Ca^{2+}$] = 4mM | 0.44±0.25 n=23 11 mice | 0.53±0.28 n=22 10 mice | -0.29 | 0.32 | 0.08 [-0.07, 0.24] | 0.32 [-0.26, 0.94] |
| Ventral root-evoked IPSC rise-time (ms) | 0.65±0.17 n=18 8 mice | 0.82±0.21 n=23 5 mice | -0.20 | 0.11 | **0.17 [0.06, 0.29]** | **0.90 [0.30, 1.65]** |
| Ventral root-evoked IPSC decay-time (ms) | 4.78±1.25 n=18 18 mice | 6.22±1.55 n=22 23 mice | -0.34 | -0.25 | **1.44 [0.62, 2.31]** | **1.01 [0.44, 1.95]** |
| aIPSC amplitude (nS) | 2.20±1.84 n=1903 19 motoneurons | 1.55±1.36 n=2173 21 motoneurons | 0.16 | 0.19 | **-0.62 [-1.06, -0.21]*** | **-0.39 [-0.64, -0.14]*** |
| aIPSC amplitude scaled (%) | 9.47±10.60 n=1903 19 motoneurons | 4.12±5.31 n=2173 21 motoneurons | 0.37 | 0.47 | **-4.98 [-8.46, -1.73]*** | **-0.62 [-0.97, -0.24]*** |
| mIPSC amplitude (nS) | 2.80±2.21 n=10743 20 motoneurons | 2.04±1.72 n=8745 23 motoneurons | 0.29 | 0.21 | **-0.78 [-1.37, -0.22]*** | **-0.40 [-0.66, -0.12]*** |
| mIPSC amplitude scaled (%) | 6.17±6.08 n=10743 20 motoneurons | 4.70±5.74 n=8745 23 motoneurons | 0.42 | 0.35 | -1.14 [-3.68, 1.53]* | -0.18 [-0.58, 0.21]* |
| mIPSC inter-event interval (ms) | 55±60 n=10670 20 motoneurons | 67±82 n=10399 23 motoneurons | **0.52** | 0.47 | **31 [7, 61]*** | **0.42 [0.14, 0.66]*** |
| mIPSC rise-time | 0.40±0.44 n=10582 20 motoneurons | 0.46±0.48 n=8832 23 motoneurons | 0.06 | 0.06 | 0.04 [-0.04, 0.12]* | 0.09 [-0.08, 0.26]* |
| mIPSC decay-time | 1.66±0.84 n=10026 20 motoneurons | 1.83±0.85 n=8118 23 motoneurons | 0.40 | 0.20 | 0.06 [-0.35, 0.41]* | 0.10 [-0.34, 0.57]* |

CI - confidence interval; EPSC - excitatory postsynaptic current; aIPSC - asynchronous inhibitory postsynaptic current; mIPSC - miniature inhibitory postsynaptic current; *2-level hierarchical bootstrapping was used for effect size estimation

**Table S6 –** Linear mixed model fixed and random-effects variables and partial eta squared ($\eta_p^2$) for glycine receptor (GlyR) clusters immunohistochemistry from P21 mice

| P21 mice Glycine receptor clusters | | | | | | | | | | |
|---|---|---|---|---|---|---|---|---|---|---|
| | Mean±SD (total observations) | | Fixed effects Estimate [95% CI] | | Random effects $\sigma^2$ (ICC) | | | | | Effect size |
| Parameter | WT | mSOD 1 | WT (intercept) | mSOD1 | Pair | Animal | MN | Bouton | Residuals | $\eta_p^2$ [90% CI] |
| GlyR area ($\mu m^2$) | 1.44E-02±1.94 E-02 | 1.39E-02±1.8 5E-02 | 1.49E-02 [ 1.32E-02 , 1.67E-02] | -6.71E-04 [-1.73E-03, 3.88E-04] | 3.19E -06 (0.01) | 0.00 (0.00) | 0.00 (0.00) | 9.60E-06 (0.03) | 3.48E-04 (0.96) | 0.00 [0.00, 0.00] |
| GlyR intensity | 17.2±7. 9 | 17.7±1 0.0 | 18.6 [ 10.7 , 26.8] | 1 [ -1.7 , 3.8] | 79 (0.81) | 4 (0.04) | 9 (0.09) | 2 (0.02) | 3 (0.04) | 0.00 [0.00, 0.00] |
| GlyR perimeter (μm) | 0.45±0. 37 | 0.45±0. 36 | 0.46 [ 0.43 , 0.48] | -0.01 [ -0.03 , 0.01 | 5.85E -04 (0.00 4) | 0.00 (0.00) | 4.65E-04 (0.004 ) | 2.49E-03 (0.02) | 1.28E-01 (0.97) | 0.00 [0.00, 0.00] |
| GlyR per bouton area (count/$\mu m^2$) | 10.0±3. 30 | 9.06±3. 18 | 9.77 [7.53, 12.01] | **-0.85 [-1.35, - 0.35]** | 6.30 (0.54) | 0.00 (0.00) | 0.71 (0.06) | N/A | 4.72 (0.40) | **0.024 [0.01, 0.05]** |

CI - confidence interval; ICC - intraclass correlation coefficient;
for WT group: 5 animals, 64 motoneurons, 217 boutons and 4892 clusters; for mSOD1 group: 5 animals, 64 motoneurons, 248 boutons and 5125 clusters. For alternative statistical analysis when $\sigma^2_{Residuals}$>0.50 $\sigma^2_{total}$ see Table S10.

**Table S7 –** Linear mixed model fixed and random-effects variables and partial eta squared ($\eta_p^2$) for glycine transporter 2 (GlyT2) immunohistochemistry from P21 mice

| P21 mice Glycine transporter 2 and motoneuron | | | | | | | | | |
|---|---|---|---|---|---|---|---|---|---|
| | Mean±SD (total observations) | | Fixed effects Estimate [95% CI] | | Random effects $\sigma^2$ (ICC) | | | | Effect size |
| Parameter | WT | mSOD1 | WT (intercept) | mSOD1 | Pair | Animal | MN | Residuals | $\eta_p^2$ [90% CI] |
| GlyT2 area (µm²) | 6.90±5.08 | 6.89±5.00 | 6.95 [6.18, 7.73] | 0.03 [-1.06, 1.11] | 0.00 (0.00) | 0.54 (0.01) | 1.76 (0.04) | 38.32 (0.94) | 0.00 [0.00, 0.00] |
| GlyT2 intensity | 22.9±6.98 | 22.3±9.14 | 22.7 [17.0, 28.4] | -0.5 [-7.5, 6.6] | 10.0 (0.13) | 30.8 (0.41) | 20.3 (0.27) | 13.4 (0.18) | 0.00 [0.00, 0.00] |
| GlyT2 perimeter (µm) | 11.60±6.53 | 11.60±6.59 | 11.61 [10.96, 12.26] | 0.05 [-0.86, 0.97] | 0.00 (0.00) | 0.32 (0.01) | 1.42 (0.03) | 41.39 (0.96) | 0.00 [0.00, 0.00] |
| Motoneuron area (µm²) | 726±159 | 838±188 | 726 [669, 783] | **113 [33, 193]** | 0.00 (0.00) | 1863 (0.06) | | 29043 (0.94) | **0.06 [0.01, 0.14]** |
| Motoneuron intensity | 4.21±1.29 | 3.82±1.52 | 4.2 [3.3, 5.2] | -0.4 [-1.6, 0.7] | 0.31 (0.14) | 0.79 (0.36) | | 1.09 (0.50) | 0.00 [0.00, 0.04] |
| Motoneuron perimeter (µm) | 114±15 | 120±17 | 114 [110, 118] | 6 [0.2, 12] | 0.00 (0.00) | 0.00 (0.00) | | 266 (1.00) | 0.03 [0.00, 0.10] |

CI - confidence interval; ICC - intraclass correlation coefficient;
for WT group: 5 animals, 64 motoneurons and 217 boutons; for mSOD1 group: 5 animals, 64 motoneurons and 248 boutons. For alternative statistical analysis when $\sigma^2_{Residuals}$>0.50 $\sigma^2_{total}$ see Table S10.

**Table S8 –** Linear mixed model fixed and random-effects variables and partial eta squared ($\eta_p^2$) for glycine receptor (GlyR) clusters immunohistochemistry from P45 mice

| P45 mice Glycine receptor clusters | | | | | | | | | | |
|---|---|---|---|---|---|---|---|---|---|---|
| | Mean±SD (total observations) | | Fixed effects Estimate [95% CI] | | Random effects $\sigma^2$ (ICC) | | | | | Effect size |
| Parameter | WT | mSOD1 | WT (intercept) | mSOD1 | Pair | Animal | MN | Bouton | Residuals | $\eta_p^2$ [90% CI] |
| GlyR area (μm$^2$) | 1.37E-02±2.08E-02 | 1.38E-02±2.13E-02 | 1.40E-02 [1.19E-02, 1.61E-02] | -3.37E-05 [-1.33E-03, 1.27E-03] | 4.62E-06 (0.01) | 0.00 (0.00) | 6.35E-06 (0.01) | 1.29E-05 (0.03) | 4.24E-04 (0.95) | 0.00 [0.00, 0.00] |
| GlyR intensity | 11.4±3.0 | 11.7±3.2 | 11.4 [9.2, 13.7] | 0.6 [-0.07, 1.2] | 6.5 (0.54) | 0.00 (0.00) | 3.4 (0.29) | 0.6 (0.05) | 1.4 (0.12) | 0.00 [0.00, 0.00] |
| GlyR perimeter (μm) | 0.44±0.38 | 0.44±0.38 | 0.45 [0.41, 0.48] | -1.34E-03 [-0.02, 0.02] | 1.44E-03 (0.01) | 0.00 (0.00) | 1.13E-03 (0.01) | 2.73E-03 (0.02) | 1.42E-0.1 (0.96) | 0.00 [0.00, 0.00] |
| GlyR per bouton area (count/μm$^2$) | 11.13±2.57 | 10.15±2.59 | 11.11 [10.26, 11.97] | **-0.98 [-1.46, -0.49]** | 0.78 (0.11) | 0.00 (0.00) | 0.67 (0.10) | | 5.38 (0.79) | **0.028 [0.01, 0.06]** |

CI - confidence interval; ICC - intraclass correlation coefficient;
for WT group: 5 animals, 72 motoneurons, 258 boutons and 6622 clusters; for mSOD1 group: 5 animals, 73 motoneurons, 275 boutons and 6203 clusters. For alternative statistical analysis when $\sigma^2_{Residuals}$>0.50 $\sigma^2_{total}$ see Table S10.

**Table S9 –** Linear mixed model fixed and random-effects variables and partial eta squared ($\eta_p^2$) for glycine receptor (GlyR) clusters immunohistochemistry from P45 mice

| P45 mice Glycine transporter 2 (GlyT2) and motoneuron | | | | | | | | | |
|---|---|---|---|---|---|---|---|---|---|
| | Mean±SD (total observations) | | Fixed effects Estimate [95% CI] | | Random effects $\sigma^2$ (ICC) | | | | Effect size |
| Parameter | WT | mSOD1 | WT (intercept) | mSOD1 | Pair | Animal | MN | Residuals | $\eta_p^2$ [90% CI] |
| GlyT2 area ($\mu m^2$) | 5.52±5.12 | 5.18±5.07 | 5.54 [5.09, 6.00] | -0.34 [-0.99, 0.30] | 0.00 (0.00) | 0.11 (0.00) | 1.44 (0.06) | 24.52 (0.94) | 0.00 [0.00, 0.00] |
| GlyT2 intensity | 19.6±7.9 | 21.0±8.8 | 19.5 [13.3, 25.6] | 1.5 [-0.5, 3.6] | 6.9 (0.43) | 1.2 (0.07) | 4.4 (0.28) | 3.6 (0.22) | 0.00 [0.00, 0.00] |
| GlyT2 perimeter (μm) | 10.28±5.90 | 9.81±5.75 | 10.30 [9.80, 10.80] | -0.48 [-1.06, 0.11] | 6.85 (0.43) | 1.18 (0.07) | 4.41 (0.28) | 3.56 (0.22) | 0.00 [0.00, 0.00] |
| Motoneuron area ($\mu m^2$) | 715±184 | 681±205 | 716 [664, 767] | -35 [-98, 29] | 822 (0.02) | 0.00 (0.00) | | 37365 (0.98) | 0.01 [0.00, 0.05] |
| Motoneuron intensity | 4.04±1.27 | 4.10±1.69 | 4.00 [2.8, 5.2] | 0.1 [-0.3, 0.4] | 1.75 (0.66) | 0.01 (0.00) | | 0.89 (0.33) | 0.00 [0.00, 0.03] |
| Motoneuron perimeter (μm) | 113±18 | 108±18 | 113 [108, 118] | -5 [-11, 1] | 11.1 (0.04) | 0.00 (0.00) | | 305 (0.96) | 0.02 [0.00, 0.08] |

CI - confidence interval; ICC - intraclass correlation coefficient;
for WT group: 5 animals, 72 motoneurons and 2144 boutons; for mSOD1 group: 5 animals, 73 motoneurons and 2103 boutons. For alternative statistical analysis when $\sigma^2_{Residuals}$>0.50 $\sigma^2_{total}$ see Table S10.

**Table S10** - Mean and standard deviation, intraclass correlation coefficient (ICC), bootstrapped mean difference and Hedges' *g* for glycine receptor (GlyR) immunohistochemistry parameters from P21 and P45 mice

| P21 mice Glycine receptor clusters | | |
|---|---|---|
| Parameter | Bootstrapped<br>Mean difference<br>Estimate [95% CI] | Bootstrapped Hedges' *g*<br>Estimate [95% CI] |
| GlyR area (μm$^2$) | -4.48E-04<br>[-1.2E-03, 3.00E-04] | -0.02<br>[-0.06, 0.02] |
| GlyR perimeter (μm) | 6.49E-03<br>[-2.07E-02, 7.54E-03] | -0.02<br>[-0.06, 0.02] |
| P21 mice Glycine transporter 2 (GlyT2) and motoneuron | | |
| GlyT2 perimeter (μm) | -2.28E-03<br>[-3.94E-01, 3.84E-01] | 0.00<br>[-0.06, 0.06] |
| Motoneuron area (μm$^2$) | **112**<br>**[53, 171]** | **0.65**<br>**[0.30, 1.02]** |
| Motoneuron intensity | -0.39<br>[-0.87, 0.10] | -0.28<br>[-0.63, 0.07] |
| Motoneuron perimeter (μm) | **6**<br>**[0.40, 12]** | **0.36**<br>**[0.03, 0.73]** |
| P45 mice Glycine receptor clusters | | |
| GlyR area (μm$^2$) | 5.34E-05<br>[-6.77E-04, 7.77E-04] | 0.00<br>[-0.03, 0.04] |
| GlyR perimeter (μm) | **-0.21**<br>**[-0.27, -0.15]** | **-0.12**<br>**[-0.16, -0.09]** |
| GlyR per bouton area (count/μm$^2$) | **-0.98**<br>**[-1.42, -0.54]** | **-0.3**<br>**[-0.55, -0.21]** |
| P45 mice Glycine transporter 2 and motoneuron | | |
| GlyT2 area (μm$^2$) | **-0.33**<br>**[-0.64, -0.03]** | **-0.07**<br>**[-0.13, -0.01]** |
| Motoneuron area (μm$^2$) | -35<br>[-97, 29] | -0.18<br>[-0.51, 0.15] |
| Motoneuron perimeter (μm) | -5<br>[-11, 1] | -0.29<br>[-0.62, 0.04] |

CI - confidence interval

**Table S11 -** Mean and standard deviation, intraclass correlation coefficient (ICC), bootstrapped mean difference and Hedges' *g* for all data gathered on dorsal root-evoked monosynaptic Ia excitation, disynaptic Ia/Ib inhibition and post-activation depression

| Ventral horn-ablated *in vitro* spinal cord | | | | | | |
|---|---|---|---|---|---|---|
| All motoneurons | | | | | | |
| Parameter | Mean±standard deviation | | ICC | | Bootstrapped Mean difference [95% CI] | Bootstrapped Hedges' *g* [95% CI] |
| | WT | mSOD1 | WT | mSOD1 | | |
| Dorsal root-evoked excitation (nS) | 33±28 n=253 25 mice | 41±26 n=222 23 mice | 0.32 | 0.40 | **8 [3, 13]** | **0.28 [0.10, 0.49]** |
| Dorsal root-evoked excitation scaled (%) | 90±108 n=252 25 mice | 99±108 n=222 23 mice | 0.17 | 0.10 | 10 [-10, 30] | 0.09 [-0.09, 0.27] |
| Dorsal root-evoked EPSC paired-pulse ratio | 1.05±0.09 n=215 23 mice | 1.01±0.10 n=212 23 mice | 0.09 | 0.23 | **-0.04 [-0.06, -0.03]** | **-0.47 [-0.28, -0.67]** |
| Dorsal root-evoked EPSC rise-time (ms) | 0.41±0.10 n=249 25 mice | 0.39±0.09 n=218 23 mice | 0.28 | 0.25 | **-0.02 [-0.04, -0.001]** | **-0.19 [-0.01, -0.37]** |
| Dorsal root-evoked EPSC decay-time (ms) | 4.47±1.38 n=249 25 mice | 4.32±1.12 n=218 23 mice | 0.41 | 0.06 | -0.15 [-0.38, -0.08] | -0.12 [-0.30, 0.05] |
| Post-activation depression (ratio) | 0.81±0.09 n=151 16 mice | 0.82±0.10 n=169 17 mice | 0.12 | 0.10 | 0.01 [-0.01, 0.03] | 0.14 [-0.08, 0.37] |
| Dorsal root-evoked inhibition (nS) | 76±45 n=126 18 mice | 82±51 n=167 18 mice | 0.21 | 0.20 | 6 [-5, 17] | 0.13 [-0.10, 0.36] |
| Dorsal root-evoked inhibition scaled (%) | 190±122 n=124 18 mice | 206±249 n=167 18 mice | 0.14 | 0.14 | 15 [-25, 62] | 0.06 [-0.20, 0.22] |
| Dorsal root-evoked IPSC paired-pulse ratio | 1.05±0.07 n=113 17 mice | 1.08±0.10 n=158 17 mice | -0.04 | 0.26 | **0.03 [0.01, 0.05]** | **0.30 [0.07, 0.52]** |
| Dorsal root-evoked IPSC rise-time (ms) | 0.53±0.11 n=124 18 mice | 0.52±0.11 n=164 18 mice | 0.40 | 0.34 | -0.01 [-0.03, 0.02] | -0.09 [-0.33, 0.15] |
| Dorsal root-evoked IPSC decay-time (ms) | 4.73±1.24 n=124 18 mice | 0.52±0.11 n=163 18 mice | 0.26 | 0.26 | **-0.34 [-0.63, -0.04]** | **-0.26 [-0.51, -0.03]** |

CI - confidence interval; ICC - intraclass correlation coefficient; EPSC - excitatory postsynaptic current; IPSC - inhibitory postsynaptic current.

**Table S12 -** Mean and standard deviation, intraclass correlation coefficient (ICC), bootstrapped mean difference and Hedges' *g* for all data gathered on dorsal root-evoked monosynaptic Ia excitation, disynaptic Ia/Ib inhibition and post-activation depression recorded from motoneurons within L4 segments

| Ventral horn-ablated *in vitro* spinal cord | | | | | | |
|---|---|---|---|---|---|---|
| L4 segment | | | | | | |
| Parameter | Mean±standard deviation | | ICC | | Bootstrapped Mean difference [95% CI] | Bootstrapped Hedges' *g* [95% CI] |
| | WT | mSOD1 | WT | mSOD1 | | |
| Dorsal root-evoked excitation (nS) | 40±36 n=113 15 mice | 36±25 n=80 13 mice | 0.35 | 0.43 | -4 [-12, 5] | -0.14 [-0.38, 0.15] |
| Dorsal root-evoked excitation scaled (%) | 114±148 n=113 16 mice | 85±67 n=80 13 mice | 0.29 | 0.18 | **-29 [-62, -0.4]** | -0.24 [-0.44, 0.01] |
| Dorsal root-evoked EPSC paired-pulse ratio | 1.06±0.08 n=100 15 mice | 0.99±0.10 n=78 13 mice | 0.18 | 0.20 | **-0.07 [-0.09, -0.04]** | **-0.73 [-1.06, -0.44]** |
| Dorsal root-evoked EPSC rise-time (ms) | 0.40±0.07 n=112 15 mice | 0.39±0.10 n=80 13 mice | 0.15 | 0.30 | -0.01 [-0.03, 0.02] | -0.07 [-0.39, 0.22] |
| Dorsal root-evoked EPSC decay-time (ms) | 4.22±1.21 n=112 15 mice | 4.15±1.14 n=80 13 mice | 0.49 | 0.20 | -0.06 [-0.40, 0.28] | -0.05 [-0.34, 0.23] |
| Post-activation depression (ratio) | 0.81±0.08 n=65 12 mice | 0.80±0.11 n=61 11 mice | 0.10 | 0.35 | -0.01 [-0.04, 0.02] | -0.11 [-0.46, 0.25] |
| Dorsal root-evoked inhibition (nS) | 87±50 n=51 13 mice | 80±58 n=58 11 mice | 0.20 | 0.16 | -7 [-27, 13] | -0.12 [-0.51, 0.25] |
| Dorsal root-evoked inhibition scaled (%) | 201±127 n=51 13 mice | 196±187 n=58 11 mice | 0.28 | 0.09 | -5 [-62, 56] | -0.03 [-0.45, 0.31] |
| Dorsal root-evoked IPSC paired-pulse ratio | 1.05±0.07 n=50 13 mice | 1.06±0.10 n=56 11 mice | 0.29 | **0.55** | 0.01 [-0.02, 0.04] | 0.09 [-0.29, 0.46] |
| Dorsal root-evoked IPSC rise-time (ms) | 0.53±0.10 n=50 13 mice | 0.48±0.09 n=58 11 mice | **0.63** | **0.60** | **-0.04 [-0.08, 0.01]** | **-0.46 [-0.86, -0.10]** |
| Dorsal root-evoked IPSC decay-time (ms) | 4.72±1.24 n=50 13 mice | 3.99±1.10 n=58 11 mice | 0.36 | **0.54** | **-0.73 [-1.17, -0.30]** | **-0.62 [-1.00, -0.27]** |

CI - confidence interval; ICC - intraclass correlation coefficient; EPSC - excitatory postsynaptic current; IPSC - inhibitory postsynaptic current; for linear mixed model results when ICC>0.50 see table S25

**Table S13 -** Mean and standard deviation, intraclass correlation coefficient (ICC), bootstrapped mean difference and Hedges' *g* for all data gathered on dorsal root-evoked monosynaptic Ia excitation, disynaptic Ia/Ib inhibition and post-activation depression recorded from motoneurons within L5 segments

| Ventral horn-ablated *in vitro* spinal cord | | | | | | |
|---|---|---|---|---|---|---|
| L5 segment | | | | | | |
| Parameter | Mean±standard deviation | | ICC | | Bootstrapped Mean difference [95% CI] | Bootstrapped Hedges' *g* [95% CI] |
| | WT | mSOD1 | WT | mSOD1 | | |
| Dorsal root-evoked excitation (nS) | 26±17 n=140 21 mice | 43±27 n=142 22 mice | 0.34 | 0.36 | **17 [11, 22]** | **0.73 [0.52, 0.95]** |
| Dorsal root-evoked excitation scaled (%) | 70±50 n=139 21 mice | 107±125 n=142 22 mice | 0.14 | 0.42 | **38 [18, 62]** | **0.39 [0.26, 0.58]** |
| Dorsal root-evoked EPSC paired-pulse ratio | 1.05±0.09 n=115 19 mice | 1.02±0.10 n=134 22 mice | 0.06 | 0.22 | **-0.03 [-0.05, -0.01]** | **-0.32 [-0.57, -0.06]** |
| Dorsal root-evoked EPSC rise-time (ms) | 0.43±0.12 n=137 21 mice | 0.40±0.09 n=138 22 mice | 0.38 | 0.31 | **-0.03 [-0.06, -0.01]** | **-0.28 [-0.52, -0.05]** |
| Dorsal root-evoked EPSC decay-time (ms) | 4.67±1.48 n=137 21 mice | 4.41±1.10 n=138 22 mice | 0.44 | 0.22 | -0.26 [-0.57, 0.04] | -0.20 [-0.44, 0.23] |
| Post-activation depression (ratio) | 0.80±0.10 n=86 15 mice | 0.83±0.09 n=107 16 mice | 0.28 | 0.04 | **0.27 [0.00, 0.05]** | **0.29 [0.01, 0.59]** |
| Dorsal root-evoked inhibition (nS) | 68±39 n=75 15 mice | 83±47 n=109 15 mice | 0.16 | 0.27 | **15 [2, 27]** | **0.33 [0.05, 0.62]** |
| Dorsal root-evoked inhibition scaled (%) | 183±119 n=73 15 mice | 210±278 n=109 15 mice | 0.14 | **0.86** | 28 [-24, 92] | 0.12 [-0.19, 0.30] |
| Dorsal root-evoked IPSC paired-pulse ratio | 1.05±0.07 n=63 14 mice | 1.09±0.10 n=102 14 mice | -0.13 | 0.31 | **0.04 [0.01, 0.06]** | **0.41 [0.13, 0.68]** |
| Dorsal root-evoked IPSC rise-time (ms) | 0.53±0.11 n=74 15 mice | 0.54±0.11 n=106 15 mice | 0.32 | 0.19 | 0.01 [-0.02, 0.04] | 0.09 [-0.21, 0.39] |
| Dorsal root-evoked IPSC decay-time (ms) | 4.74±1.25 n=74 15 mice | 4.62±1.43 n=105 15 mice | 0.31 | 0.29 | -0.13 [-0.54, 0.26] | -0.10 [-0.41, 0.19] |

CI - confidence interval; ICC - intraclass correlation coefficient; EPSC - excitatory postsynaptic current; IPSC - inhibitory postsynaptic current; for linear mixed model results when ICC>0.50 see table S25

**Table S14 -** Mean and standard deviation, intraclass correlation coefficient (ICC), bootstrapped mean difference and Hedges' *g* for all data gathered on L4 dorsal root-evoked monosynaptic Ia excitation, disynaptic Ia/Ib inhibition and post-activation depression recorded from motoneurons within L4 segments

| Ventral horn-ablated *in vitro* spinal cord | | | | | | |
|---|---|---|---|---|---|---|
| L4 segment and L4 dorsal root stimulation | | | | | | |
| Parameter | Mean±standard deviation | | ICC | | Bootstrapped Mean difference [95% CI] | Bootstrapped Hedges' *g* [95% CI] |
| | WT | mSOD1 | WT | mSOD1 | | |
| Dorsal root-evoked excitation (nS) | 44±36 n=57 15 mice | 44±26 n=39 11 mice | 0.33 | 0.44 | 0 [-13, 12] | -0.01 [-0.36, 0.41] |
| Dorsal root-evoked excitation scaled (%) | 119±109 n=57 15 mice | 105±76 n=39 11 mice | 0.35 | 0.02 | -14 [-51, 22] | -0.14 [-0.49, 0.25] |
| Dorsal root-evoked EPSC paired-pulse ratio | 1.07±0.07 n=49 15 mice | 1.00±0.09 n=38 11 mice | 0.23 | 0.25 | **0.07 [-0.10, -0.03]** | -0.84 [-1.34, 0.42] |
| Dorsal root-evoked EPSC rise-time (ms) | 0.40±0.08 n=57 15 mice | 0.38±0.10 n=39 11 mice | 0.04 | 0.26 | -0.02 [-0.06, 0.01] | -0.27 [-0.76, 0.15] |
| Dorsal root-evoked EPSC decay-time (ms) | 4.45±1.25 n=57 15 mice | 4.27±1.24 n=39 11 mice | 0.45 | 0.06 | -0.18 [-0.67, 0.34] | -0.15 [-0.57, 0.27] |
| Post-activation depression (ratio) | 0.82±0.06 n=32 12 mice | 0.82±0.10 n=28 10 mice | 0.13 | **0.65** | 0.00 [-0.05, 0.04] | -0.05 [-0.57, 0.51] |
| Dorsal root-evoked inhibition (nS) | 85±54 n=29 13 mice | 103±63 n=31 10 mice | 0.00 | 0.34 | 18 [-11, 47] | 0.31 [-0.19, 0.86] |
| Dorsal root-evoked inhibition scaled (%) | 207±139 n=29 13 mice | 255±216 n=31 10 mice | -0.02 | 0.23 | 49 [-39, 143] | 0.27 [-0.26, 0.71] |
| Dorsal root-evoked IPSC paired-pulse ratio | 1.05±0.06 n=29 13 mice | 1.04±0.09 n=30 10 mice | 0.43 | **0.68** | -0.01 [-0.04, 0.04] | -0.05 [-0.64, 0.42] |
| Dorsal root-evoked IPSC rise-time (ms) | 0.53±0.12 n=28 13 mice | 0.47±0.09 n=31 10 mice | **0.63** | **0.64** | **-0.06 [-0.11, -0.01]** | **-0.55 [-1.07, -0.06]** |
| Dorsal root-evoked IPSC decay-time (ms) | 4.83±1.20 n=28 13 mice | 4.30±1.17 n=31 10 mice | 0.04 | 0.45 | -0.54 [-1.13, 0.07] | -0.45 [-0.99, 0.05] |

CI - confidence interval; ICC - intraclass correlation coefficient; EPSC - excitatory postsynaptic current; IPSC - inhibitory postsynaptic current; for linear mixed model results when ICC>0.50 see table S25

**Table S15 -** Mean and standard deviation, intraclass correlation coefficient (ICC), bootstrapped mean difference and Hedges' *g* for all data gathered on L5 dorsal root-evoked monosynaptic Ia excitation, disynaptic Ia/Ib inhibition and post-activation depression recorded from motoneurons within L4 segments

| Ventral horn-ablated *in vitro* spinal cord | | | | | | |
|---|---|---|---|---|---|---|
| L4 segment and L5 dorsal root stimulation | | | | | | |
| Parameter | Mean±standard deviation | | ICC | | Bootstrapped Mean difference [95% CI] | Bootstrapped Hedges' *g* [95% CI] |
| | WT | mSOD1 | WT | mSOD1 | | |
| Dorsal root-evoked excitation (nS) | 36±36<br>n=56<br>14 mice | 29±22<br>n=41<br>13 mice | 0.28 | **0.56** | -8<br>[-20, 3] | -0.27<br>[-0.59, 0.11] |
| Dorsal root-evoked excitation scaled (%) | 108±180<br>n=56<br>14 mice | 65±50<br>n=41<br>13 mice | 0.26 | 0.40 | **-44**<br>**[-99, -1]** | **-0.32**<br>**[-0.54, -0.02]** |
| Dorsal root-evoked EPSC paired-pulse ratio | 1.04±0.09<br>n=51<br>14 mice | 0.98±0.11<br>n=40<br>13 mice | 0.09 | 0.29 | **-0.06**<br>**[-0.11, -0.02]** | **-0.66**<br>**[-1.11, -0.25]** |
| Dorsal root-evoked EPSC rise-time (ms) | 0.39±0.06<br>n=55<br>14 mice | 0.40±0.10<br>n=41<br>13 mice | 0.25 | **0.54** | 0.01<br>[-0.02, 0.05] | 0.15<br>[-0.28, 0.58] |
| Dorsal root-evoked EPSC decay-time (ms) | 3.98±1.12<br>n=55<br>14 mice | 4.05±1.05<br>n=41<br>13 mice | 0.50 | 0.36 | 0.07<br>[-0.37, 0.50] | 0.06<br>[-0.34, 0.48] |
| Post-activation depression (ratio) | 0.80±0.09<br>n=33<br>10 mice | 0.78±0.12<br>n=33<br>10 mice | **0.57** | 0.28 | -0.01<br>[-0.06, 0.04] | -0.13<br>[-0.61, 0.37] |
| Dorsal root-evoked inhibition (nS) | 90±47<br>n=22<br>10 mice | 54±39<br>n=27<br>9 mice | 0.20 | 0.40 | **-36**<br>**[-59, -12]** | **-0.83**<br>**[-1.51, -0.30]** |
| Dorsal root-evoked inhibition scaled (%) | 194±111<br>n=22<br>10 mice | 129±116<br>n=27<br>9 mice | 0.50 | -0.11 | **-65**<br>**[-128, -3]** | **-0.57**<br>**[-1.38, -0.03]** |
| Dorsal root-evoked IPSC paired-pulse ratio | 1.05±0.08<br>n=21<br>10 mice | 1.07±0.11<br>n=26<br>9 mice | 0.36 | **0.53** | 0.02<br>[-0.03, 0.07] | 0.23<br>[-0.32, 0.84] |
| Dorsal root-evoked IPSC rise-time (ms) | 0.52±0.07<br>n=22<br>10 mice | 0.49±0.10<br>n=27<br>9 mice | 0.43 | **0.52** | -0.03<br>[-0.07, 0.02] | -0.32<br>[-0.98, 0.21] |
| Dorsal root-evoked IPSC decay-time (ms) | 4.58±1.31<br>n=22<br>10 mice | 3.64±0.92<br>n=27<br>9 mice | 0.27 | **0.62** | **-0.93**<br>**[-1.60, -0.34]** | **-0.84**<br>**[-1.34, -0.37]** |

CI - confidence interval; ICC - intraclass correlation coefficient; EPSC - excitatory postsynaptic current; IPSC - inhibitory postsynaptic current; for linear mixed model results when ICC>0.50 see table S25

**Table S16 -** Mean and standard deviation, intraclass correlation coefficient (ICC), bootstrapped mean difference and Hedges' *g* for all data gathered on L4 dorsal root-evoked monosynaptic Ia excitation, disynaptic Ia/Ib inhibition and post-activation depression recorded from motoneurons within L5 segments

| Ventral horn-ablated *in vitro* spinal cord | | | | | | |
|---|---|---|---|---|---|---|
| L5 segment and L4 dorsal root stimulation | | | | | | |
| Parameter | Mean±standard deviation | | ICC | | Bootstrapped Mean difference [95% CI] | Bootstrapped Hedges' *g* [95% CI] |
| | WT | mSOD1 | WT | mSOD1 | | |
| Dorsal root-evoked excitation (nS) | 30±19 n=63 17 mice | 42±26 n=66 19 mice | **0.55** | 0.19 | **12 [4, 20]** | **0.54 [0.22, 0.87]** |
| Dorsal root-evoked excitation scaled (%) | 76±49 n=63 17 mice | 108±143 n=66 19 mice | 0.30 | 0.37 | **32 [2, 73]** | **0.29 [0.04, 0.57]** |
| Dorsal root-evoked EPSC paired-pulse ratio | 1.06±0.08 n=60 17 mice | 1.02±0.10 n=62 19 mice | 0.19 | 0.15 | **-0.05 [-0.08, -0.01]** | **-0.51 [-0.91, -0.14]** |
| Dorsal root-evoked EPSC rise-time (ms) | 0.46±0.14 n=63 17 mice | 0.41±0.09 n=64 19 mice | 0.46 | 0.10 | **-0.05 [-0.09, -0.01]** | **-0.39 [-0.75, -0.05]** |
| Dorsal root-evoked EPSC decay-time (ms) | 4.82±1.68 n=63 17 mice | 4.46±1.04 n=64 19 mice | 0.45 | 0.29 | -0.36 [-0.85, 0.10] | -0.26 [-0.61, 0.08] |
| Post-activation depression (ratio) | 0.82±0.11 n=35 11 mice | 0.81±0.07 n=53 14 mice | 0.17 | 0.03 | -0.02 [-0.06, 0.02] | -0.18 [-0.61, 0.29] |
| Dorsal root-evoked inhibition (nS) | 72±40 n=39 11 mice | 80±47 n=49 13 mice | 0.17 | 0.36 | 8 [-10, 26] | 0.18 [-0.24, 0.59] |
| Dorsal root-evoked inhibition scaled (%) | 195±106 n=39 11 mice | 232±392 n=49 13 mice | -0.07 | **0.94** | 37 [-51, 173] | 0.12 [-0.50, 0.35] |
| Dorsal root-evoked IPSC paired-pulse ratio | 1.05±0.07 n=34 11 mice | 1.09±0.08 n=47 13 mice | 0.19 | 0.26 | **0.04 [0.01, 0.07]** | **0.50 [0.10, 0.90]** |
| Dorsal root-evoked IPSC rise-time (ms) | 0.56±0.13 n=39 11 mice | 0.57±0.12 n=49 13 mice | 0.47 | -0.06 | 0.02 [-0.04, 0.07] | 0.13 [-0.32, 0.55] |
| Dorsal root-evoked IPSC decay-time (ms) | 4.63±1.39 n=39 11 mice | 4.54±1.39 n=49 13 mice | 0.26 | 0.09 | -0.09 [-0.65, 0.50] | -0.07 [-0.54, 0.33] |

CI - confidence interval; ICC - intraclass correlation coefficient; EPSC - excitatory postsynaptic current; IPSC - inhibitory postsynaptic current; for linear mixed model results when ICC>0.50 see table S25

**Table S17 –** Mean and standard deviation, intraclass correlation coefficient (ICC), bootstrapped mean difference and Hedges' *g* for all data gathered on L5 dorsal root-evoked monosynaptic Ia excitation, disynaptic Ia/Ib inhibition and post-activation depression recorded from motoneurons within L5 segments

| Ventral horn-ablated *in vitro* spinal cord | | | | | | |
|---|---|---|---|---|---|---|
| L5 segment and L5 dorsal root stimulation | | | | | | |
| Parameter | Mean±standard deviation | | ICC | | Bootstrapped Mean difference Estimate [95% CI] | Bootstrapped Hedges' *g* Estimate [95% CI] |
| | WT | mSOD1 | WT | mSOD1 | | |
| Dorsal root-evoked excitation (nS) | 24±16<br>n=77<br>19 mice | 44±27<br>n=76<br>20 mice | 0.25 | 0.47 | **20<br>[13, 27]** | **0.90<br>[0.61, 1.20]** |
| Dorsal root-evoked excitation scaled (%) | 65±50<br>n=76<br>19 mice | 107±107<br>n=76<br>20 mice | 0.07 | 0.33 | **42<br>[18, 72]** | **0.50<br>[0.29, 0.81]** |
| Dorsal root-evoked EPSC paired-pulse ratio | 1.04±0.11<br>n=55<br>17 mice | 1.02±0.10<br>n=72<br>20 mice | 0.20 | 0.19 | -0.01<br>[-0.05, 0.02] | -0.15<br>[-0.49, 0.21] |
| Dorsal root-evoked EPSC rise-time (ms) | 0.40±0.10<br>n=74<br>19 mice | 0.38±0.09<br>n=74<br>20 mice | 0.28 | **0.52** | -0.02<br>[-0.05, 0.01] | -0.18<br>[-0.50, 0.15] |
| Dorsal root-evoked EPSC decay-time (ms) | 4.55±1.29<br>n=74<br>19 mice | 4.37±1.17<br>n=74<br>20 mice | 0.42 | 0.28 | -0.18<br>[-0.57, 0.21] | -0.15<br>[-0.49, 0.17] |
| Post-activation depression (ratio) | 0.79±0.09<br>n=51<br>14 mice | 0.85±0.10<br>n=54<br>14 mice | 0.23 | -0.06 | **0.06<br>[0.03, 0.10]** | **0.68<br>[0.31, 1.08]** |
| Dorsal root-evoked inhibition (nS) | 64±38<br>n=36<br>15 mice | 85±47<br>n=60<br>12 mice | 0.28 | 0.25 | **21<br>[4, 38]** | **0.48<br>[0.09, 0.89]** |
| Dorsal root-evoked inhibition scaled (%) | 169±133<br>n=34<br>15 mice | 193±123<br>n=60<br>12 mice | 0.19 | 0.06 | 25<br>[-31, 77] | 0.19<br>[-0.24, 0.66] |
| Dorsal root-evoked IPSC paired-pulse ratio | 1.06±0.06<br>n=29<br>14 mice | 1.09±0.11<br>n=55<br>12 mice | -0.30 | 0.31 | 0.03<br>[-0.01, 0.07] | 0.33<br>[-0.05, 0.69] |
| Dorsal root-evoked IPSC rise-time (ms) | 0.50±0.08<br>n=35<br>15 mice | 0.51±0.08<br>n=57<br>12 mice | 0.18 | **0.61** | 0.03<br>[-0.01, 0.07] | 0.15<br>[-0.28, 0.57] |
| Dorsal root-evoked IPSC decay-time (ms) | 4.86±1.09<br>n=35<br>15 mice | 4.69±1.47<br>n=56<br>12 mice | **0.51** | 0.32 | -0.20<br>[-0.71, 0.33] | -0.15<br>[-0.58, 0.24] |

CI - confidence interval; ICC - intraclass correlation coefficient; EPSC - excitatory postsynaptic current; IPSC - inhibitory postsynaptic current

**Table S18 –** Mean and standard deviation, bootstrapped mean difference and Hedges' *g* for motoneuron resting conductance and capacitance and latency and jitter obtained from dorsal root-evoked monosynaptic EPSCs and disynaptic IPSCs from *in vitro* partially-ventral horn ablated spinal cord preparations

| Parameter | Mean±standard deviation | | Bootstrapped Mean difference Estimate [95% CI] | Bootstrapped Hedges' *g* Estimate [95% CI] |
|---|---|---|---|---|
| | WT | mSOD1 | | |
| Motoneuron conductance (nS) | 46±28<br>n=247<br>35 mice | 47±22<br>n=198<br>30 mice | 0.9<br>[-4, 5] | 0.04<br>[-0.14, 0.23] |
| Motoneuron capacitance (pF) | 260±79<br>n=243<br>35 mice | 271±86<br>n=199<br>30 mice | 11<br>[-4, 27] | 0.14<br>[-0.05, 0.33] |
| | EPSC | IPSC | | |
| WT latency (ms) | 0.80±0.11<br>n=68<br>16 mice | 0.85±0.12<br>n=68<br>16 mice | **0.05**<br>**[0.03, 0.07]** | **0.40**<br>**[0.24, 0.59]** |
| WT jitter (ms) | 0.014±0.006<br>n=68<br>16 mice | 0.036±0.020<br>n=68<br>16 mice | **0.019**<br>**[0.015, 0.024]** | **1.28**<br>**[1.05, 1.61]** |
| mSOD1 latency (ms) | 0.79±0.12<br>n=125<br>16 mice | 0.86±0.14<br>n=125<br>16 mice | **0.07**<br>**[0.06, 0.08]** | **0.55**<br>**[0.41, 0.71]** |
| mSOD1 jitter (ms) | 0.013±0.004<br>n=125<br>16 mice | 0.029±0.017<br>n=125<br>16 mice | **0.015**<br>**[0.013, 0.019]** | **1.32**<br>**[1.07, 1.67]** |

CI - confidence interval; EPSC - excitatory postsynaptic current; IPSC - inhibitory postsynaptic current

**Table S19 -** Mean and standard deviation, intraclass correlation coefficient (ICC), bootstrapped mean difference and Hedges' *g* for Bayesian Quantal Analysis (BQA) performed on monosynaptic Ia excitation

| *In vitro* ventral horn-ablated spinal cord | | | | | | |
|---|---|---|---|---|---|---|
| Bayesian Quantal Analysis | | | | | | |
| Parameter | Mean±standard deviation | | ICC | | Bootstrapped Mean difference Estimate [95% CI] | Bootstrapped Hedges' *g* Estimate [95% CI] |
| | WT | mSOD1 | WT | mSOD1 | | |
| Quantal size (nS) | 0.38±0.14<br>n=36<br>13 mice | 0.37±0.19<br>n=27<br>9 mice | -0.10 | 0.22 | -0.01<br>[-0.09, 0.07] | -0.09<br>[-0.64, 0.44] |
| Quantal size scaled (%) | 1.46±1.15<br>n=36<br>13 mice | 1.58±1.46<br>n=27<br>9 mice | **0.52** | 0.09 | 0.11<br>[-0.52, 0.80] | 0.09<br>[-0.46, 0.58] |
| Number of release sites | 245±123<br>n=37<br>14 mice | 201±132<br>n=27<br>9 mice | -0.03 | 0.13 | -44<br>[-106, 18] | -0.35<br>[-0.87, 0.15] |
| Probability of release $[Ca^{2+}]$ = 2mM | 0.23±0.10<br>n=37<br>14 mice | 0.33±0.15<br>n=27<br>9 mice | -0.05 | 0.06 | **0.11**<br>**[0.05, 0.17]** | **0.83**<br>**[0.32, 1.42]** |
| Probability of release $[Ca^{2+}]$ = 4mM | 0.48±0.17<br>n=36<br>14 mice | 0.63±0.23<br>n=27<br>9 mice | 0.05 | -0.22 | **0.16**<br>**[0.06, 0.26]** | **0.79**<br>**[0.31, 1.40]** |

CI - confidence interval;  ICC - intraclass correlation coefficient

**Table S20 -** Mean and standard deviation, bootstrapped mean difference and Hedges' *g* for recurrent inhibition studied through electromyography (EMG) recordings

| | Parameter | Mean±standard deviation | | Bootstrapped Mean difference Estimate [95% CI] | Bootstrapped Hedges' *g* Estimate [95% CI] |
|---|---|---|---|---|---|
| Age range | Conditioning interval | WT | mSOD1 | | |
| P18-23 | 2 ms (% control H-reflex) | 101±28 n=13 mice | 107±27 n=11 mice | 6 [-15, 27] | 0.22 [-0.60, 1.08] |
| | 5 ms (% control H-reflex) | 102±37 n=15 mice | 101±20 n=12 mice | -1 [-22, 19] | -0.04 [-0.69, 0.90] |
| | 10 ms (% control H-reflex) | 89±14 n=14 mice | 105±18 n=10 mice | **16 [4, 29]** | **1.02 [0.27, 1.98]** |
| | 20 ms (% control H-reflex) | 82±25 n=14 mice | 99±8 n=12 mice | **17 [3, 30]** | **0.85 [0.21, 1.70]** |
| | 30 ms (% control H-reflex) | 82±13 n=14 mice | 99±11 n=11 mice | **17 [8, 26]** | **1.41 [0.71, 2.58]** |
| | 50 ms (% control H-reflex) | 80±22 n=14 mice | 98±11 n=10 mice | **17 [5, 31]** | **0.94 [0.42, 1.63]** |
| | 100 ms (% control H-reflex) | 89±16 n=14 mice | 98±16 n=8 mice | 9 [-3, 22] | 0.60 [-0.25, 1.53] |
| P57-67 | 2 ms (% control H-reflex) | 102±30 n=11 mice | 84±22 n=11 mice | -18 [-41, 2] | -0.67 [-1.56, 0.11] |
| | 5 ms (% control H-reflex) | 85±23 n=11 mice | 83±23 n=11 mice | -2 [-20, 16] | -0.09 [-0.97, 0.78] |
| | 10 ms (% control H-reflex) | 82±18 n=11 mice | 85±23 n=10 mice | 3 [-14, 20] | 0.14 [-0.81, 1.05] |
| | 20 ms (% control H-reflex) | 94±17 n=11 mice | 84±19 n=10 mice | -10 [-26, 3] | -0.58 [-1.39, 0.27] |
| | 30 ms (% control H-reflex) | 96±16 n=11 mice | 96±8 n=10 mice | 0 [-10, 10] | 0.01 [-1.00, 0.83] |
| | 50 ms (% control H-reflex) | 92±13 n=11 mice | 102±4 n=10 mice | **9 [1, 16]** | **0.94 [0.10, 2.75]** |
| P87-108 | 2 ms (% control H-reflex) | 101±17 n=10 mice | 81±24 n=11 mice | **-20 [-38, -4]** | **-0.93 [-1.88, -0.20]** |
| | 5 ms (% control H-reflex) | 96±24 n=10 mice | 85±18 n=11 mice | -10 [-28, 6] | -0.49 [-1.37, 0.38] |
| | 10 ms (% control H-reflex) | 87±23 n=9 mice | 78±29 n=11 mice | -10 [-31, 12] | -0.36 [-1.44, 0.47] |
| | 20 ms (% control H-reflex) | 88±21 n=10 mice | 98±20 n=11 mice | 10 [-6, 28] | 0.50 [-0.37, 1.32] |
| | 30 ms (% control H-reflex) | 103±19 n=9 mice | 93±19 n=11 mice | -9 [-26, 5] | -0.49 [-1.29, 0.45] |
| | 50 ms (% control H-reflex) | 93±16 n=9 mice | 97±10 n=11 mice | 3 [-7, 15] | 0.24 [-0.91, 1.05] |

CI - confidence interval;  ICC - intraclass correlation coefficient

**Table S21 -** Mean and standard deviation, bootstrapped mean difference and Hedges' *g* for post-activation depression studied through EMG recordings

| | Parameter | Mean±standard deviation | | Bootstrapped Mean difference Estimate [95% CI] | Bootstrapped Hedges' *g* Estimate [95% CI] |
|---|---|---|---|---|---|
| Age range | Conditioning interval | WT | mSOD1 | | |
| P18-23 | 50 ms (% control H-reflex) | 37±16 n=15 mice | 47±22 n=9 mice | 9 [-6, 26] | 0.51 [-0.38, 1.37] |
| | 100 ms (% control H-reflex) | 46±20 n=15 mice | 45±24 n=8 mice | 0 [-18, 19] | -0.02 [-1.05, 0.87] |
| | 200 ms (% control H-reflex) | 47±15 n=14 mice | 54±23 n=10 mice | 7 [-9, 23] | 0.36 [-0.55, 1.29] |
| | 500 ms (% control H-reflex) | 54±19 n=14 mice | 64±22 n=11 mice | 10 [-5, 26] | 0.51 [-0.27, 1.40] |
| | 1 s (% control H-reflex) | 70±23 n=14 mice | 64±27 n=10 mice | -6 [-25, 13] | -0.26 [-1.24, 0.58] |
| | 5 s (% control H-reflex) | 83±15 n=15 mice | 97±13 n=10 mice | **15 [4, 25]** | **1.01 [0.25, 2.17]** |
| | 10 s (% control H-reflex) | 97±7 n=15 mice | 102±11 n=10 mice | 4 [-3, 12] | 0.52 [-0.35, 1.41] |
| | 20 s (% control H-reflex) | 102±5 n=13 mice | 107±6 n=11 mice | **6 [1, 10]** | **0.99 [0.19, 2.29]** |
| P57-67 | 50 ms (% control H-reflex) | 50±19 n=11 mice | 52±20 n=6 mice | 2 [-16, 20] | 0.09 [-0.98, 1.18] |
| | 100 ms (% control H-reflex) | 54±20 n=11 mice | 56±15 n=7 mice | 2 [-13, 17] | 0.13 [-0.86, 1.06] |
| | 200 ms (% control H-reflex) | 55±18 n=11 mice | 63±16 n=7 mice | 8 [-7, 24] | 0.48 [-0.45, 1.49] |
| | 500 ms (% control H-reflex) | 62±15 n=11 mice | 75±13 n=7 mice | **13 [1, 25]** | **0.92 [0.15, 1.91]** |
| | 1 s (% control H-reflex) | 70±16 n=11 mice | 80±14 n=7 mice | 10 [-4, 22] | 0.63 [-0.24, 1.82] |
| | 5 s (% control H-reflex) | 88±10 n=11 mice | 94±12 n=7 mice | 6 [-4, 16] | 0.59 [-0.37, 1.82] |
| | 10 s (% control H-reflex) | 94±10 n=11 mice | 100±12 n=7 mice | 6 [-4, 16] | 0.55 [-0.41, 1.92] |
| | 20 s (% control H-reflex) | 103±5 n=11 mice | 102±6 n=6 mice | -1 [-6, 4] | -0.20 [-1.39, 0.90] |
| P87-108 | 50 ms (% control H-reflex) | 49±20 n=9 mice | 51±21 n=11 mice | 1 [-16, 18] | 0.05 [-0.93, 0.93] |
| | 100 ms (% control H-reflex) | 54±21 n=9 mice | 62±19 n=13 mice | 9 [-7, 25] | 0.45 [-0.41, 1.65] |
| | 200 ms (% control H-reflex) | 53±18 n=9 mice | 70±22 n=12 mice | **17 [0.1, 33]** | **0.81 [0.03, 2.11]** |
| | 500 ms (% control H-reflex) | 70±16 n=9 mice | 79±19 n=11 mice | 9 [-6, 23] | 0.52 [-0.33, 1.68] |
| | 1 s (% control H-reflex) | 72±16 n=9 mice | 95±15 n=13 mice | **23 [10, 36]** | **1.46 [0.73, 2.63]** |
| | 5 s (% control H-reflex) | 92±13 n=9 mice | 97±11 n=13 mice | 5 [-5, 14] | 0.40 [-0.46, 1.47] |
| | 10 s (% control H-reflex) | 99±9 n=9 mice | 97±13 n=13 mice | -2 [-11, 7] | -0.14 [-0.92, 0.81] |
| | 20 s (% control H-reflex) | 101±4 n=9 mice | 102±4 n=13 mice | 1 [-2, 4] | 0.17 [-0.75, 1.10] |

CI - confidence interval

**Table S22 -** Mean and standard deviation, bootstrapped mean difference and Hedges' *g* for Mmax, Hmax and $H_{max}/M_{max}$ (ratio) for Quadriceps and Tibialis Anterior muscles studied through EMG recordings

| | Parameter | Mean±standard deviation | | Bootstrapped Mean difference Estimate [95% CI] | Bootstrapped Hedges' *g* Estimate [95% CI] |
|---|---|---|---|---|---|
| Age range | Conditioning interval | WT | mSOD1 | | |
| P18-23 | Quadriceps $M_{max}$ (mV) | 10±4 n=18 mice | 8±3 n=18 mice | -1 [-3, 2] | -0.35 [-1.08, 0.29] |
| | Tibialis Anterior $M_{max}$ (mV) | 21±6 n=21 mice | 17±7 n=19 mice | **-4 [-8, -0.5]** | **-0.71 [-1.48, -0.10]** |
| | Tibialis Anterior $H_{max}$ (mV) | 4±2 n=20 mice | 3±2 n=17 mice | **-2 [-3, -0.1]** | -0.44 [-1.19, 0.21] |
| | Tibialis Anterior $H_{max}/M_{max}$ (ratio) | 0.21±0.11 n=20 mice | 0.23±0.17 n=17 mice | 0.02 [-0.07, 0.12] | 0.14 [-0.61, 0.73] |
| P57-67 | Quadriceps $M_{max}$ (mV) | 13±8 n=14 mice | 7±4 n=15 mice | **-6 [-10, -1]** | **-0.93 [-1.73, -0.32]** |
| | Tibialis Anterior $M_{max}$ (mV) | 25±9 n=14 mice | 13±5 n=15 mice | **-11 [-18, -5]** | **-1.62 [-2.73, -1.08]** |
| | Tibialis Anterior $H_{max}$ (mV) | 3±1 n=11 mice | 3±1 n=10 mice | 0 [-1, 1] | -0.27 [-1.27, 0.58] |
| | Tibialis Anterior $H_{max}/M_{max}$ (ratio) | 0.11±0.05 n=11 mice | 0.21±0.05 n=10 mice | **0.11 [0.06, 0.16]** | **1.81 [1.00, 3.39]** |
| P87-108 | Quadriceps $M_{max}$ (mV) | 11±6 n=14 mice | 7±5 n=15 mice | **-5 [-9, -0.2]** | **-0.77 [-1.91, -0.02]** |
| | Tibialis Anterior $M_{max}$ (mV) | 19±6 n=14 mice | 10±2 n=16 mice | **-10 [-13, -6]** | **-1.83 [-2.71, -1.33]** |
| | Tibialis Anterior $H_{max}$ (mV) | 3±2 n=12 mice | 3±1 n=13 mice | 0 [-1, 1] | -0.10 [-0.88, 0.82] |
| | Tibialis Anterior $H_{max}/M_{max}$ (ratio) | 0.17±0.12 n=12 mice | 0.30±0.15 n=13 mice | **0.13 [0.02, 0.23]** | **0.92 [0.14, 2.02]** |

CI - confidence interval

**Table S23 -** Mean and standard deviation, intraclass correlation coefficient (ICC), bootstrapped mean difference and Hedges' *g* for recurrent inhibition estimated from *in vivo* and *in vitro* dorsal horn-ablated motoneuron recordings

| Ventral horn-ablated *in vitro* spinal cord | | | | | | |
|---|---|---|---|---|---|---|
| *In vivo* recordings | | | | | | |
| Parameter | Mean±standard deviation | | ICC | | Bootstrapped Mean difference Estimate [95% CI] | Bootstrapped Hedges' *g* Estimate [95% CI] |
| | WT | mSOD1 | WT | mSOD1 | | |
| Recurrent inhibition (nS) | 29±23 n=22 6 mice | 26±37 n=18 7 mice | 0.38 | **0.64** | -4 [-22, 16] | -0.12 [-0.98, 0.49] |
| Recurrent inhibition scaled (%) | 10±11 n=22 6 mice | 7±9 n=18 7 mice | 0.25 | **0.55** | -3 [-9, 3] | -0.27 [-0.85, 0.37] |
| Dorsal horn-ablated *in vitro* spinal cord | | | | | | |
| Early firing motoneurons | | | | | | |
| Parameter | Mean±standard deviation | | ICC | | Bootstrapped Mean difference Estimate [95% CI] | Bootstrapped Hedges' *g* Estimate [95% CI] |
| | WT | mSOD1 | WT | mSOD1 | | |
| Recurrent inhibition (nS) | 27±25 n=20 7 mice | 9±9 n=14 6 mice | 0.19 | 0.07 | **-18 [-30, -7]** | **-0.91 [-1.49, -0.41]** |
| Recurrent inhibition scaled (%) | 61±68 n=20 7 mice | 31±44 n=14 6 mice | -0.02 | -0.08 | -29 [-66, 8] | -0.42 [-0.97, 0.21] |
| Delayed firing motoneurons | | | | | | |
| Parameter | Mean±standard deviation | | ICC | | Bootstrapped Mean difference Estimate [95% CI] | Bootstrapped Hedges g Estimate [95% CI] |
| | WT | mSOD1 | WT | mSOD1 | | |
| Recurrent inhibition (nS) | 63±50 n=126 11 mice | 45±33 n=142 9 mice | 0.27 | 0.35 | **-18 [-29, -8]** | **-0.44 [-0.65, -0.21]** |
| Recurrent inhibition scaled (%) | 89±86 n=126 11 mice | 53±35 n=142 9 mice | 0.43 | 0.24 | **-36 [-53, -20]** | **-0.55 [-0.74, -0.36]** |

CI - confidence interval; ICC - intraclass correlation coefficient; for linear mixed model results when ICC>0.50 see table S26

**Table S24 –** Linear mixed model fixed and random-effects variables and partial eta squared ($\eta_p^2$) for electrophysiological parameters obtained from early and delayed firing motoneurons and Renshaw cells recorded from oblique spinal cord slices

| Oblique spinal cord slices | | | | | | |
|---|---|---|---|---|---|---|
| | | Fixed effects Estimate [95% CI] | | Random effects $\sigma^2$ (ICC) | | Effect size |
| | Parameter | WT (intercept) | mSOD1 | Animal | Residuals | $\eta_p^2$ [90% CI] |
| Early firing motoneurons | Conductance (nS) | 52 [39, 65] | 4 [-16, 24] | 198 (0.15) | 1102 (0.85) | 0.00 [0.00, 0.06] |
| | Membrane potential (mV) | -62 [-64, -60] | 0 [-2, 3] | 5.0 (0.34) | 9.8 (0.66) | 0.00 [0.00, 0.04] |
| | 1-2$^{nd}$ spike interval (Hz) | 391 [268, 514] | 35 [-120, 191] | 11131 (0.47) | 12746 (0.53) | 0.00 [0.00, 0.16] |
| | Action potential half-width (ms) | 0.23 [0.19, 0.28] | 0.03 [-0.04, 0.10] | 7.50E-03 (0.84) | 1.41E-03 (0.16) | 0.01 [0.00, 0.12] |
| | Action potential rise time (ms) | 0.18 [0.16, 0.20] | 0.00 [-0.02, 0.03] | 7.50E-03 (0.88) | 1.07E-03 (0.12) | 0.01 [0.00, 0.07] |
| | Action potential decay time (ms) | 0.16 [0.11, 0.21] | 0.04 [-0.04, 0.11] | 8.15E-03 (0.86) | 1.38E-03 (0.14) | 0.02 [0.00, 0.14] |
| | Recurrent excitation (ns) | 10 [6, 13] | 0 [-6, 6] | 17 (0.38) | 28 (0.62) | 0.00 [0.00, 0.00] |
| | Recurrent excitation scaled (%) | 32 [18, 46] | -10 [-31, 12] | 242 (0.40) | 355 (0.60) | 0.03 [0.00, 0.19] |
| | Ventral root-evoked EPSC rise-time (ms) | 0.46 [0.36, 0.55] | 0.07 [-0.07, 0.21] | 9.1E-04 (0.86) | 1.44E-02 (0.14) | 0.04 [0.00, 0.24] |
| | Ventral root-evoked EPSC decay-time (ms) | 4.58 [2.64, 6.53] | 0.62 [-2.41, 3.66] | 8.27 (0.92) | 0.71 (0.08) | 0.00 [0.00, 0.15] |
| Delayed firing motoneurons | Ventral root-evoked EPSC rise-time (ms) | 0.50 [0.42, 0.58] | 0.06 [-0.06, 0.18] | 1.54E-02 (0.58) | 1.13E-02 (0.42) | 0.02 [0.00, 0.11] |
| | Ventral root-evoked EPSC decay-time (ms) | 5.65 [4.69, 6.61] | -0.48 [-1.89, 0.94] | 1.75 (0.47) | 1.94 (0.53) | 0.00 [0.00, 0.07] |
| Renshaw cells | BQA probability of release [Ca2+] = 1mM | 0.19 [0.08, 0.30] | 0.10 [-0.08, 0.28] | 1.16E-02 (0.65) | 8.97E-03 (0.35) | 0.06 [0.00, 0.28] |
| | BQA probability of release [Ca2+] = 2mM | 0.58 [0.40, 0.76] | 0.26 [-0.03, 0.55] | 3.14E-02 (0.43) | 4.21E-03 (0.57) | 0.15 [0.00, 0.39] |

CI - confidence interval; ICC - intraclass correlation coefficient; EPSC - excitatory postsynaptic current; BQA - Bayesian Quantal Analysis

**Table S25 –** Linear mixed model fixed and random-effects variables and partial eta squared ($\eta_p^2$) for electrophysiological parameters obtained from ventral horn-ablated spinal cord slices

| *In vitro* ventral horn-ablated spinal cord | | | | | | |
|---|---|---|---|---|---|---|
| | | Fixed effects Estimate [95% CI] | | Random effects $\sigma^2$ (ICC) | | Effect size |
| | Parameter | WT (intercept) | mSOD1 | Animal | Residuals | $\eta_p^2$ [90% CI] |
| L4 segment L4 and L5 stimulation | Dorsal root-evoked IPSC paired-pulse ratio | 1.05 [1.01, 1.10] | 0.03 [-0.04, 0.09] | 4.71E-03 (0.51) | 4.44E-03 (0.49) | 0.01 [0.00, 0.06] |
| | Dorsal root-evoked IPSC rise-time (ms) | 0.55 [0.49, 0.60] | -0.05 [-0.13, 0.04] | 8.91E-03 (0.69) | 3.97E-03 (0.31) | 0.01 [0.00, 0.07] |
| | Dorsal root-evoked IPSC decay-time (ms) | 4.80 [4.14, 5.46] | -0.81 [-1.78, 0.16] | 1.13 (0.58) | 0.83 (0.42) | 0.03 [0.00, 0.10] |
| L5 segment L4 and L5 stimulation | Dorsal root-evoked inhibition scaled (%) | 179 [-60, 418] | 164 [-174, 503] | 21.7E04 (0.95) | 12.2E03 (0.05) | 0.01 [0.00, 0.04] |
| L4 segment L4 root stimulation | Post-activation depression (ratio) | 0.83 [0.78, 0.87] | -0.02 [-0.10, 0.05] | 5.35E-03 (0.58) | 3.931E-03 (0.42) | 0.00 [0.00, 0.02] |
| | Dorsal root-evoked IPSC paired-pulse ratio | 1.05 [1.00, 1.10] | 6.38E-03 [-0.06, 0.08] | 5.54E-03 (0.66) | 2.84E-03 (0.34) | 0.00 [0.00, 0.04] |
| | Dorsal root-evoked IPSC rise-time (ms) | 0.55 [0.49, 0.61] | -0.07 [-0.16, 0.03] | 9.76E-03 (0.69) | 4.36E-06 (0.31) | 0.04 [0.00, 0.15] |
| L4 segment L5 root stimulation | Dorsal root-evoked excitation (nS) | 35 [23, 47] | -6 [-24, 12] | 297 (0.31) | 650 (0.69) | 0.01 [0.00, 0.06] |
| | Dorsal root-evoked EPSC rise-time (ms) | 0.39 [0.35, 0.42] | 0.02 [-0.03, 0.07] | 3.08E-0.3 (0.47) | 3.49E-0.3 (0.53) | 0.01 [0.00, 0.06] |
| | Post-activation depression (ratio) | 0.81 [0.75, 0.87] | 0.05 [-0.12, 0.04] | 5.10E-0.3 (0.41) | 7.34E-0.3 (0.59) | 0.02 [0.00, 0.11] |
| | Dorsal root-evoked IPSC paired-pulse ratio | 1.05 [0.99, 1.10] | 0.03 [-0.05, 0.11] | 4.78E-0.3 (0.47) | 5.39E-0.3 (0.53) | 0.01 [0.00, 0.12] |
| | Dorsal root-evoked IPSC rise-time (ms) | 0.53 [0.48, 0.57] | -0.02 [-0.10, 0.05] | 3.82E-0.3 (0.48) | 4.20E-0.3 (0.52) | 0.01 [0.00, 0.11] |
| | Dorsal root-evoked IPSC decay-time (ms) | 4.68 [4.02, 5.35] | -1.15 [-2.09, -0.20] | 0.634 (0.43) | 0.833 (0.57) | 0.12 [0.01, 0.27] |
| L5 segment L4 root stimulation | Dorsal root-evoked excitation (nS) | 30 [22, 38] | 10 [-2, 22] | 173 (0.32) | 374 (0.68) | 0.02 [0.00, 0.08] |
| | Dorsal root-evoked inhibition scaled (%) | 191 [-129, 511] | 169 [-267, 604] | 281E03 (0.96) | 11E03 (0.04) | 0.01 [0.00, 0.06] |
| L5 segment L5 root stimulation | Dorsal root-evoked EPSC rise-time (ms) | 0.40 [0.36, 0.43] | -0.02 [-0.07, 0.03] | 3.43E-03 (0.39) | 5.45E-03 (0.61) | 0.00 [0.00, 0.04] |

| | | | | | | |
|---|---|---|---|---|---|---|
| | Dorsal root-evoked IPSC rise-time (ms) | 0.49 [0.45, 0.53] | 0.02 [-0.04, 0.08] | 4.50E-03 (0.54) | 3.86E-03 (0.46) | 0.00 [0.00, 0.05] |
| | Dorsal root-evoked IPSC decay-time (ms) | 4.85 [4.27, 5.43] | -0.16 [-0.97, 0.66] | 0.66 (0.34) | 1.26 (0.66) | 0.00 [0.00, 0.04] |

CI - confidence interval; ICC - intraclass correlation coefficient; EPSC - excitatory postsynaptic current; IPSC - inhibitory postsynaptic current; BQA - Bayesian Quantal Analysis

**Table S26 –** Linear mixed model fixed and random-effects variables and partial eta squared for quantal size estimated from Bayesian Quantal Analysis (BQA) experiments obtained from *in vitro* ventral horn ablated spinal cords and conductance of recurrent inhibition estimated from *in vivo* motoneuron recordings

| *In vitro* motoneuron recordings from ventral horn-ablated spinal cords | | | | | |
|---|---|---|---|---|---|
| | Fixed effects<br>Estimate<br>[95% CI] | | Random effects<br>$\sigma^2$<br>(ICC) | | Effect size |
| Parameter | WT<br>(intercept) | mSOD1 | Animal | Residuals | $\eta_p^2$<br>[90% CI] |
| Quantal size scaled (%) | 1.45<br>[0.89, 2.01] | 0.07<br>[-0.80, 0.94] | 0.49<br>(0.65) | 2.19<br>(0.35) | 0.00<br>[0.00, 0.03] |
| *In vivo* motoneuron recordings | | | | | |
| Recurrent inhibition (nS) | 28<br>[3, 53] | 10<br>[-25, 45] | 803<br>(0.65) | 440<br>(0.35) | 0.00<br>[0.00, 0.12] |
| Recurrent inhibition scaled (%) | 11<br>[4, 17] | -2<br>[-11, 7] | 36<br>(0.31) | 79<br>(0.69) | 0.01<br>[0.00, 0.10] |

CI - confidence interval; ICC - intraclass correlation coefficient;

**Appendix**

The framework for Bayesian quantal analysis implementation with improved discrete-grid exact inference (BQA-DGEI) is similar to that used previously[1] except evaluated as a single posterior distribution rather than across a family of posteriors that are subsequently subjected to conditional marginalisation. If $\Xi$ denotes data observations described by parameters $\Pi$, the posterior probability density function $f(\Pi|\Xi)$ can be expressed with respect Bayes rule:

$$f(\Pi|\,\Xi) \;= \frac{f(\Pi)f(\Xi|\Pi)}{f(\Xi)} \tag{A.1}$$

The data observations $\Xi$ comprise three quantities. First the synaptic responses $\boldsymbol{\mathcal{X}}$ comprising $c$ vectors $\mathbf{x}_i$, where $0 < i < c - 1$ and $c$ denotes the number of conditions of release probability. Second, the accompanying indices $\boldsymbol{\mathcal{K}}$ comprising $c$ vectors $\mathbf{k}_i$ where $k=i$ to inform the model which release condition of release probability is associated with each observed response. Finally the standard deviation of the baseline noise $\varepsilon$ is measured from the raw data without attempt of probabilistic treatment.

In order to characterise the joint probability density function for the prior $f(\Pi)$, marginal independent priors are assigned explicitly for the quantal parameters $\Pi$ comprising the number of release sites $n$, the quantal size $q$, and a shaping parameter $\gamma$ that functionally determines the coefficient of variation of the uniquantal distribution. For each parameter, independent priors are assigned in accordance with Jeffrey's rule, that when applied to the number of release sites $n$, a discrete variable, assigns a marginal prior explicitly as a Zipf proportionality: $f(n) \propto 1/\,n$. Since $q$ and $\gamma$ are continuously, they are most efficiently sampled on a log scale so that the corresponding marginal priors can be treated as uniform. Applying product rule, the resulting prior $f(\Pi) = f(n,\log(q),\log(\gamma))$ is three-dimensional with probabilities that change only with respect to $n$: $f(n,\log(q),\log(\gamma))$ ) $\propto 1/\,n$.

As previously adopted[1], the uniquantal distribution selected is the gamma probability $\Gamma$ density function $\boldsymbol{\mathcal{G}}(x|\gamma,\lambda)$ expressed with respect to a shaping parameter $\gamma$ and a scaling parameter $\lambda$:

$$\mathcal{G}(x|\gamma,\lambda) = \frac{1}{\lambda^{\gamma}\Gamma(\gamma)}x^{\gamma-1}e^{-x/\lambda}$$

where:

$$\Gamma(\gamma) = \int_0^{\infty} g^{\gamma-1}e^{-g}dg$$

Since the quantal size $q$ represents the mean of the uniquantal distribution with expectation $q = \gamma\lambda$, the gamma scaling parameter $\lambda$ constitutes a latent variable expressed by the relation $\lambda = q/\gamma$. The magnitude of baseline noise is measured by the standard deviation of failed responses $\varepsilon$ and its distribution is modelled using a zero-mean Gaussian probability density function $\boldsymbol{\mathcal{N}}\,(x|0,\,\varepsilon)$:

$$\mathcal{N}(x|0,\varepsilon) = \frac{1}{\sqrt{2\pi\varepsilon^2}}e^{-x^2/2\varepsilon^2}$$

Average responses for each condition of release probability were evaluated using the arithmetic means $\mu_i$. If $p_i$ denotes the corresponding probabilities of release, its relation with the mean response $\mu_i$ can be expressed as a product: $\mu_i = np_iq$. The probability of release thus constitutes another latent variable determined by the relation: $p_i = \mu_i/nq$. For a given release probability *p* a binomial probability mass function_$\mathcal{B}(j\ |n, p)$ is used to model the probability of observing *j* successful releases from *n* sites each with identical Bernoulli probabilities of *p*:

$$\mathcal{B}(j|n,p) = \frac{n!}{j!\,(n-j)!}p^j(1-p)^{n-1}$$

For each index *k* of release probability, the composite quantal amplitude function $\mathcal{Q}(x|k)$ is described by a summation that weights the binomial probability mass function $\mathcal{B}(j\ |n,p)$ over the baseline noise normal probability density function $\mathcal{N}\ (x|0, \varepsilon^2)$ and a series of gamma probability density functions $\mathcal{G}(x|j\gamma, \lambda)$ representing a sequential convolution over *n* Bernoulli trials:

$$\mathcal{Q}(x|k) = \mathcal{B}(0|n,p_k)\mathcal{N}(x|0,\varepsilon) + \sum_{j=1}^{n}\mathcal{B}(j|n,p_k)\mathcal{G}(x|j\gamma,\lambda)$$

Product rule, including multiplication of the composite quantal amplitude function with a Gaussian, to model the probability density function for the mean responses $\mu_i$ for each condition of release probability, completes the expression of likelihood function $f(\mathcal{X},\mathcal{K}|n,\log(q),\log(\gamma))$:

$$f(\mathcal{X},\mathcal{K}|n,\log(q),\log(\gamma)) = \prod_{i=0}^{c-1}\mathcal{N}(\mu_i|\mu_i,\mathrm{s.e.}[\mathbf{x}_i])\prod_{x\in\mathbf{x}_i}\mathcal{Q}(x|k)$$

where $\mathrm{s.e.}[\mathbf{x}_i]$ is the standard error for the observations $\mathbf{x}_i$ for each condition in release probability. Notably this likelihood function differs from the quantal likelihood function published previously[1] since this Gaussian term was absent from the original version. The main advantage of its inclusion is the improved numerical stability that arises from conferring greater weight to composite quantal amplitude distributions associated with larger numbers of observations compared to those with fewer. The procedure is analogous to the mean-variance parabolic fitting used for multiple probability fluctuation analysis weighting the moment data according the error bars on the variances.

In practice the product for the likelihood function is more efficiently computed with respect to the corresponding logarithms since this converts products to summations.

$$\mathcal{L} = \sum_{i=0}^{c-1}\log\big(\mathcal{N}(0|0,s.e.[\boldsymbol{x}_i])\big) + \sum_{x\in\mathbf{x}_i}\log\big(\mathcal{Q}(x|k)\big) + L$$

Where $\mathcal{L}$ is the log likelihood $\log\left(f(\mathcal{X},\mathcal{K}|n,\log(q),\log(\gamma))\right)$ offset by an arbitrary constant *L* set to particular value, such as the negative mean log likehood, to optimise the floating point 64-bit precision for addition operations. Using exponentation of the log likelihood totals and marginalisation of the denominator term $f(\mathcal{X},\mathcal{K})$, the final posterior $f(n,\log(q),\log(\gamma)\,|\mathcal{X},\mathcal{K})$ can be expressed by applying Bayes rule in Equation A.1:

$$f(n,\log(q),\log(\gamma)\,|\mathcal{X},\mathcal{K}) = \frac{f(n,\log(q),\log(\gamma))f(\mathcal{X},\mathcal{K}|n,\log(q),\log(\gamma))}{f(\mathcal{X},\mathcal{K})}$$

$$= \frac{f(n, \log(q), \log(\gamma)) f(\mathcal{X}, \mathcal{K} | n, \log(q), \log(\gamma))}{\int_{\mathcal{A}_n} \int_{\mathcal{S}_q} \int_{\mathcal{S}_\gamma} f(n, \log(q), \log(\gamma)) f(\mathcal{X}, \mathcal{K} | n, \log(q), \log(\gamma)) \, d\gamma dq dn} \qquad (A.2)$$

Where $\mathcal{A}_n$ denotes the alphabet set for the number of release sites *n* and $\mathcal{S}_q$ and $\mathcal{S}_\gamma$ denote the support sets of the quantal size and shaping parameter respectively. Markov-chain Monte-Carlo sampling of the posterior however cannot be performed efficiently, for example using Hamiltonian dynamics, since the discrete variable $n$ renders the probability density function non-differentiable for this term. While vanilla Metropolis-Hastings sampling is tractable, it is bedevilled by inefficient random walk behaviour. Fortunately since only two of the quantal parameters are continuous, *viz* the quantal size $q$ and shaping parameter $\gamma$, it is very efficient to perform discrete grid exact inference, for example with a resolution of 128 for $q$ and 64 for $\gamma$. The integrals of Equation A.2 are thus replaced with summations.

Prior limits for *n* are chosen explicitly in the range [1, $n_{max}$] where $n_{max}$ is set by the operator. The limits for $\gamma$ are set implicitly with respect to a range of coefficient of variation $v$ of [0.05, 1.0], since $\gamma = 1/v^2$. By combining the prior limits for *n* with the range of known mean responses $\mu_i$ across all *c* conditions of release probability, the limits for *q* are set implicitly in accordance with a tractable range in probability of release of [0.04, 0.96], assuming the relation $q=\mu_i/np$. Having computed the posterior $f(n, \log(q), \log(\gamma) | \mathcal{X}, \mathcal{K})$ over the entire grid, marginal posteriors are evaluated by summation:

$$f(n|\mathcal{X}, \mathcal{K}) = \sum_{q \in \mathbf{q}} \sum_{\gamma \in \boldsymbol{\gamma}} f(n, \log(q), \log(\gamma) | \mathcal{X}, \mathcal{K})$$

$$f(\log(q)|\mathcal{X}, \mathcal{K}) = \sum_{n \in \mathbf{n}} \sum_{\gamma \in \boldsymbol{\gamma}} f(n, \log(q), \log(\gamma) | \mathcal{X}, \mathcal{K})$$

$$f(\log(\gamma)|\mathcal{X}, \mathcal{K}) = \sum_{n \in \mathbf{n}} \sum_{q \in \mathbf{q}} f(n, \log(q), \log(\gamma) | \mathcal{X}, \mathcal{K})$$

Estimates for the parameters $(n, q, \gamma)$ are obtained from the marginal posteriors using the half-quantile of the distribution. The estimate for the gamma scaling parameter $\lambda$ is computer using the relation $\lambda = q/\gamma$ and the probabilities of release for each of the mean responses $\mu_i$ are estimated using the relation $p_i = \mu_i/nq$.
